# Enhanced Golden Gate Assembly: Evaluating Overhang Strength for Improved Ligation Efficiency

**DOI:** 10.1101/2022.09.09.507109

**Authors:** Patryk Strzelecki, Nicolas Joly, Pascal Hébraud, Elise Hoffmann, Grzegorz M. Cech, Anna Kloska, Florent Busi, Wilfried Grange

## Abstract

Molecular cloning, a routine yet essential technique, relies heavily on efficient ligation, which can be significantly improved using Golden Gate Assembly (GGA). A key component of GGA is the use of type IIS enzymes, which uniquely cleave downstream of their recognition sequences to generate various overhangs, including non-palindromic ones. Recent advancements in GGA include the development of newly engineered enzymes with enhanced activity. Additionally, high-throughput GGA assays, which allow for the simultaneous study of all possible overhangs, have identified optimal GGA substrates with high efficiencies and fidelities, greatly facilitating the design of complex assemblies. Interestingly, these assays reveal unexpected correlations between ligation efficiencies and overhang stabilities. One hypothesis for this observation is that newly hydrolyzed DNA fragments with strong overhangs can readily re-ligate, thereby slowing down the overall process. In this paper, we employ a combination of gel electrophoresis and numerical calculations to test this hypothesis, ultimately determining that it does not hold true under the conditions established by conventional GGA assays. Using an assembly of 10 fragments, we demonstrate that strong overhangs yield higher GGA efficiency, while weak overhangs result in lower efficiency. These findings enable us to propose optimal overhangs for efficient GGA assays, significantly increasing yield.

## INTRODUCTION

Type IIS restriction endonucleases recognize non-palindromic DNA sequences, typically 5 to 6 nucleotides in length, and cleave at a defined distance from these sites (1; 2). These enzymes have significantly advanced fast and reliable DNA cloning techniques, such as the Golden Gate Assembly (GGA) method (3). GGA has been utilized to insert large assemblies of up to 52 DNA fragments (4), generate libraries of engineered metabolic pathways (5), and greatly enhance the yield of DNA constructs in single-molecule assays (6). These methods employ relatively long primers (e.g., 30 to 40 nucleotides) to introduce the recognition sequence of type IIS enzymes at both ends of Polymerase Chain Reaction (PCR) products. Since the recognition sequence is lost after cleavage, the appropriate choice of non-palindromic overhangs prevents self-ligation, resulting in a high yield of target products in a one-pot reaction. This reaction involves mixing the ligase, restriction endonuclease, and all amplified DNA fragments together in a single tube (7; 8).

In recent years, high-throughput sequencing-based GGA assays have been developed to measure ligation frequencies of substrates digested by various type IIS enzymes (9). These studies assess both the efficiency (*i*.*e*., the ability of an overhang to ligate its complementary pair through Watson–Crick base pairing) and the fidelity (*i*.*e*., the ability of an overhang to ligate with a mismatch-containing sequence). These high-throughput assays generate a vast amount of data that cannot be obtained by analyzing individual sequences one at a time. For 4-base pair (bp) overhangs, this includes a total of 256 possible substrates and thousands of possible mismatches. Given that ligase recognizes nicked double-stranded DNA (dsDNA) (10; 11), it would be expected that strong overhangs would exhibit the highest ligation efficiencies, while weak overhangs would exhibit the lowest. However, it was found that ligation efficiencies are poorly correlated with GC content (12; 9). To explain this observation, the authors noted that GGA involves the melting of DNA prior to ligation and proposed that this step may slow down the overall process. Specifically, newly hydrolyzed DNA fragments with strong overhangs, which melt more slowly than weak ones, may hinder assembly as they are more prone to re-ligation. However, experiments conducted with significantly fewer substrates (13 or 35 fragments) and presumably suboptimal ligation efficiencies, as indicated by high-throughput assays, resulted in a substantial number of transformants (9). The authors suggest that high-throughput experiments exhibit a complex equilibrium that may not directly apply to practical GGA experiments, where a specific number of overhangs with minimal or no mismatch ligations are combined. Still, based on the results of their high-throughput assays, the authors report on the ligation efficiency values for assemblies consisting of significantly fewer fragments (8). The probability that high-throughput assays would erroneously detect mismatch ligations appears to be low. However, whether the reported relative efficiencies and fidelities are correct remains speculative at this stage. Additionally, whether the melting of strong overhangs impacts GGA assembly needs further investigation. This is critically important as online tools, developed from these assays and offered to the scientific community, recommend optimal substrates for GGA assembly (8).

In this study, we explore GGA from both experimental and theoretical perspectives. First, we conduct time-dependent GGA experiments using individual substrates to mitigate potential biases from substrate competition when determining reaction efficiencies. By integrating gel electrophoresis with quantitative and automated analysis, we assess the efficiency of GGA (*i*.*e*., Watson-Crick base pairing) across six specific substrates, which were identified by high-throughput GGA assays as having either exceptionally high or low ligation efficiencies. Based on these data, we develop a kinetic model indicating that strong overhangs, when isolated in the reaction mixture (*i*.*e*., without competitive partners), enhance ligation efficiencies at the enzyme concentrations and substrate molarities commonly recommended for GGA assays. Subsequently, we assemble 10 fragments in a GGA experiment using overhangs predicted to exhibit - according to high-throughput assays - either low or high ligation efficiencies, while maintaining high fidelity (*i*.*e*., the likelihood of any given overhang binding to another is nearly zero). In agreement with our model and hypothesis, we find that strong overhangs greatly increase ligation efficiency in experimental conditions proposed in GGA assays. These findings have profound implications for GGA, enabling us to propose optimal experimental conditions for GGA.

## MATERIALS AND METHODS

All enzymes were obtained from New England Biolabs (NEB). Unless otherwise noted, chemicals were obtained from Merck.

### Golden Gate Assembly-like assays with 2 fragments only

Experiments were performed with T4 DNA ligase (M0202) at a final concentrations of 12 U/*µ*L and BsaI-HFv2 (R3733) at a final concentration of 0.6 U/*µ*L in CutSmart or rCutSmart Buffer supplemented with 1 mM ATP (P0756, NEB) and 10 mM dithiothreitol (DTT, 43816) in a ∼10 *µ*L volume. All experiments were performed with DNA molecules at molarities equal or less than 21 nM. For cycling experiments (37°C for 5 minutes and 16°C for 5 minutes, 5 minutes GGA-cycling), tubes were placed in a thermal cycler. For static experiments (37°C), a dry bath was used. While the time is well defined for static experiments, it takes about 10 to 20 seconds for the thermal cycler to reach and stabilize the temperature and so there might be an uncertainty in time of 5 to 10% for those experiments. Reactions were stopped with the addition of sodium dodecyl sulfate (SDS, 71725) and an incubation at 65°C for at least 10 minutes. We have kept reaction volumes as low as possible (about 10 *µ*L) to ensure proper thermal exchange in the cycler. We also have performed additional experiments at varying ligase concentrations (3 U/*µ*L, 12 U/*µ*L and 48 U/*µ*L) and a fixed BsaI-HFv2 concentration of 0.65 U/*µ*L. Detailed protocols are provided in the Supplementary Text, where we also discuss the typical concentrations used in GGA assays and explain the rationale behind our decision to use different concentrations in our experiments involving 2 fragments only.

### 10 fragments Golden Gate Assembly

As in (9), we have cloned a cassette of the lac operon into the pGGAselect destination vector and screened blue and white colonies. Here, our assembly consists of 9 fragments of about 300 bp and 1 fragment of about 2,200 bp (simply because lacI may be dispensable for blue/white screening and so equally long fragments could yield biased results). Using a genetic algorithm (*i*.*e*. an adaptive heuristic algorithm), we selected 2 populations (defined as optimal and non-optimal) consisting of sets of 9 overhangs with a fidelity of ∼1 (see below). Those overhangs were chosen on the basis of previous sequencing-based experiments that have estimated the number of ligation frequencies for all possible overhang pairs, *i*.*e*. a 256 × 256 elements matrix for 4 bases overhangs. Here, we define the fidelity *F* of the assembly as:

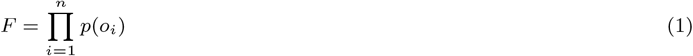

where *p*(*o*_*i*_) is the probability for a given overhang to correctly bind its complementary overhang and reads :

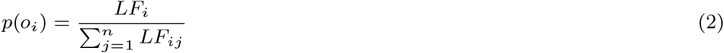

where *LF*_*ij*_ denotes the ligation frequency between the overhang *i* and all possible overhangs in the set (*n*); *LF*_*i*_ is the ligation frequency between the overhang *i* and its complementary overhang (diagonal element of the matrix). Assuming 9 overhangs, *n* is equal to (9*×*2 + 2*×*2) as complementary overhangs have to be taken into account (in addition, there are 4 extra overhangs. 2 for pGGAselect and 2 for the two fragments that directly ligate to the vector).

The optimal population consists of sets with overhangs that have a stability greater than 4.5 kcal/mol (absolute value). Their fidelities (approximately 1) have been estimated using the methods described in (4) and (9), which provide efficiency values for static and cycling GGA assays using BsaI-HFv2 and T4 DNA ligase. In contrast, the non-optimal population does not impose any stability restrictions. From this population, we estimated the frequency of overhangs with stabilities above -3.5 kcal/mol, between -3.5 and -4.5 kcal/mol, and below -4.5 kcal/mol. We found the frequencies to be 1/3 for each range. We then selected sets for the non-optimal population, ensuring that the chosen sets consisted of overhangs representative of the population’s stability distribution. Additionally, we discarded `TNNA` overhangs, as they have been previously found to significantly decrease ligation efficiencies (9). In both the optimal and non-optimal populations, multiple sets were available. We selected one set for each population based on two criteria: (*i*) the overhangs in the set could be found in the cassette and produce fragments of the desired length, and (*ii*) the overhangs in the optimal set have low ligation efficiency (according to high-throughput assays), while those in the non-optimal set have high ligation efficiency. In other words, our optimal set should produce significantly fewer fragments than the non-optimal set if the ligation efficiencies are accurate. Assemblies were performed using the NEBridge Golden Gate Assembly Kit (E1601) following the recommended protocol (Instruction Manual of the NEB^®^ Golden Gate Assembly Kit (BsaI-HF^®^v2), NEB #E1601S/L, Version 2.0 2/20, see Supplementary Text). As stated above, the assembly consists in 9 short fragments of about 300 bp and a long fragment of about 2,200 bp. Fragments were synthesized using a high-fidelity polymerase (Q5). Detailed protocols are provided in the Supplementary Text,

### Primers and sequence context

Note that the sequences upstream of the recognition site differ between the forward and reverse primers. This difference is evident in both the experiments conducted on 2 fragments and on 10 fragments. Indeed, the upstream sequences may need to be adjusted to prevent the formation of hairpin dimers or hairpins. For further details, please refer to the Supplementary Information.

### Gel assays and analysis (GGA with 2 fragments only)

1.2 % agarose gels (in Tris-acetate-EDTA buffer) were used to follow the kinetic of GGA experiments. Gels were post-stained in ethidium bromide (EtBr, 0.5 *µ*g/ml) for 15 minutes and destained in water for 10 minutes (twice). The quantification of DNA bands (intensity) was performed automatically using a code written in R. Details are also given in Supplementary Figure S3. To quantify the stoichiometry of GGA experiments, gels contain two lanes with *αα*′ and *β*′*β* fragments at time zero. From those, we calculate the ratio *S*_*β*_ defined as:

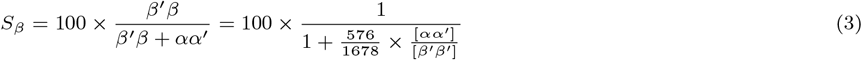

where *β*′*β* and *αα*′ are intensities (determined from gel assays). Here we have used the length of both PCR fragments (565 bp for *αα*′ and 1678 bp for *β*′*β*, see Supplementary Protocols). For our experiments, [*αα*′]=21 nM and [*β*′*β*]= 7 nM; therefore the expected *S*_*β*_ value is ∼49.7.

### Kinetic model

We solved differential equations using the R package deSolve and the R function lsoda. Details are given in the Supplementary Text.

### Figures

Violin plots were made using the ggstatsplot package (13).

## RESULTS AND DISCUSSION

### Two fragments reaction

We first present GGA experiments performed on various sequences with different overhang stabilities, as reported by (14) and different stacking energies, determined from previous nicking experiments (15). Here, we define a sequence as a succession of 6 nucleotides, where nucleotides in positions 2,3,4 and 5 form the non-palindromic overhang generated by the restriction endonuclease BsaI-HFv2. In contrast, nucleotides in positions 1 and 6 govern stacking energies (see Supplementary Tables S1 and S2 for energy values). These experiments were performed at a static temperature of 37°C or by cycling the temperature between 37°C and 16°C (5 minutes at each temperature) as in (9). The kinetics was determined by measuring the concentration of products at 10, 30, 70, 140 and 240 minutes. Specifically, nucleotides in positions 2,3,4 and 5 have been chosen by looking at the results of GGA sequencing-based assays (9). Based on this study, we selected 4 bp overhangs with the highest ligation efficiencies: `AGGA`, normalized ligation efficiency of 739 (reported values are normalized to 100,000 ligation events) and `AGAA` and `TTCT` (see Figure 2 of (9), reporting averaged ligation events for BsaI-HFv2, BsmBI-v2, Esp3I and BbsI-HF). These overhangs are considered to be equivalent and non discernable in (9) and yield an averaged ligation efficiency of 731. Similarly, we have selected the 2 lowest ligation efficiency values (247 for `CCGC` and 234 for `CACC`, respectively). Here the 2 fragments *αα*′ (565 bp) and *β*′*β* (1678 bp) synthesized by PCR (Figure 1 and Supplementary Figure S1) were mixed in a 3:1 ratio (final molarity of 21 and 7 nM, respectively) and T4 DNA ligase and BsaI-HFv2 were added to the reaction tubes. Both *α* and *β* do not contain the recognition site of BsaI-HFv2 (see Figure 1) so that the product *αβ* remains stable after ligation. In contrast, all other ligations yield non-stable products (*αα*′, *β*′*β* and *β*′*α*′), which can be re-digested as they do contain the recognition site of BsaI-HFv2. Note that nucleotides in positions 1 and 6, adjacent to the overhang, have been carefully chosen to mimic what happens in a real GGA experiment (e.g. assembling a PCR fragment in a destination vector) where stacking energies are usually not conserved among the different possible products (see Figure 1 and Supplementary Figure S2). Except for `GCCGCC` (see Supplementary Table S1), all our experiments reproduce this behavior.

**Fig. 1:**
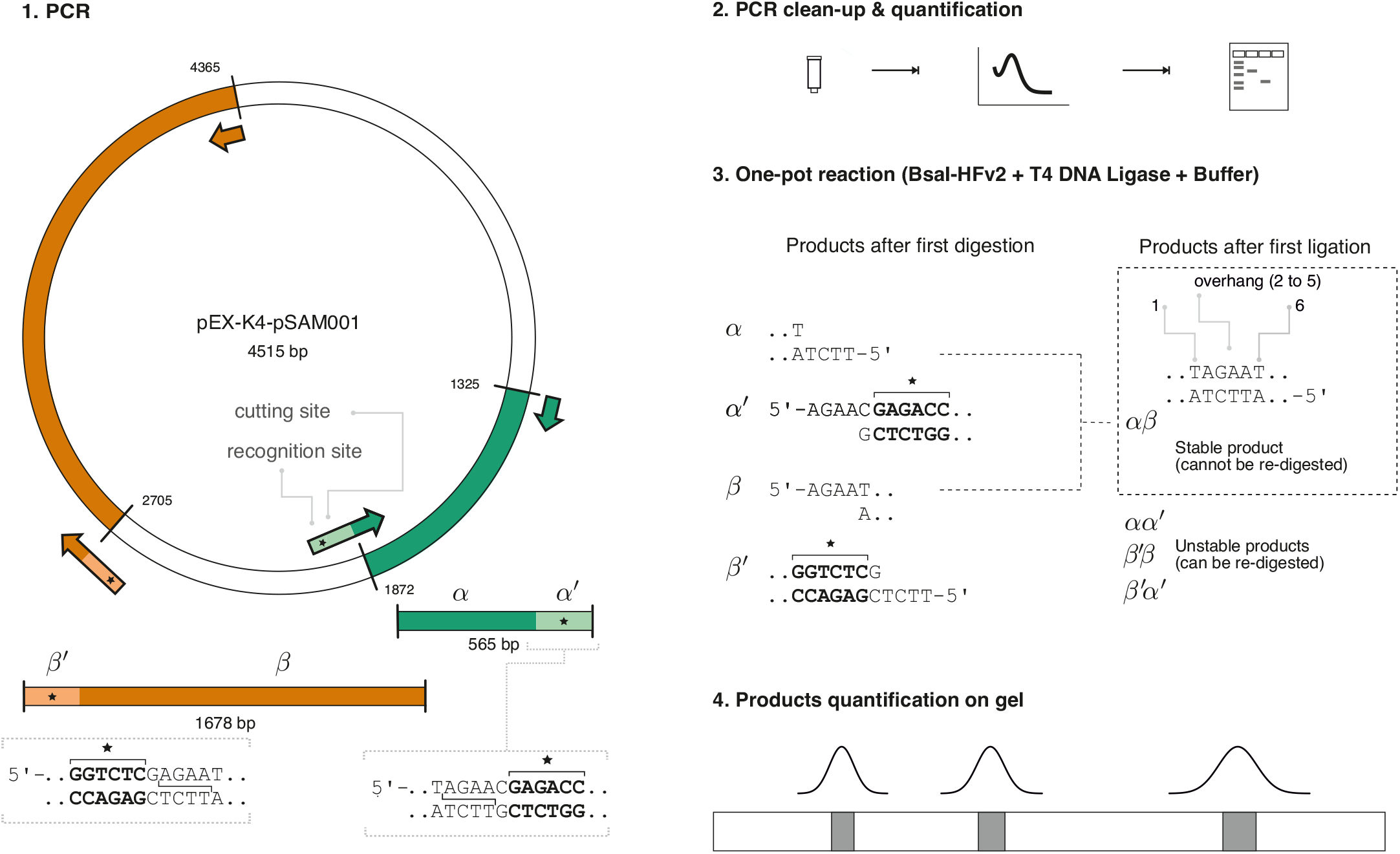
Experimental workflow for the Golden Gate Assembly (GGA) assay involving 2 substrates only. First, two PCR reactions are performed on the pEX-K4-pSAM001 plasmid (4515 bp) using primers that introduce both the recognition site of the type IIS restriction endonuclease BsaI-HFv2 (`GGTCTC`, asterisk) and specific sequences located downstream the recognition site (step 1, also refer to Supplementary Figure S2). After column purification of PCR products, determination of DNA concentration (at 260 nm) and visualization of DNA fragments by electrophoresis (step 2), a one-pot reaction is performed (step 3). Here, both PCR fragments (*αα*′, 565 bp and *β*′*β*, 1678 bp) are mixed at a given molarity and stoichiometry together with BsaI-HFv2 and T4 DNA ligase. Among the fragments produced by BsaI-HFv2, only those not containing BsaI-HFv2’s recognition sites yield a stable product after ligation. Shown here are nucleotides for the `TAGAAT` sequence, which is a sequence of six nucleotides containing a non palindromic 4 bp overhang at position 2. Note that nucleotides at positions 1 and 6 may also play a role as they govern stacking energies. Here, and except for the `GCCGCC` sequence, nucleotides at positions 1 and 6 vary among possible ligation products *αβ, αα*′, *β*′*β* and *β*′*α*′. To determine the percentage of product formed Ψ_*β*_ (defined as 100 *×* [*αβ*]*/*([*β*′*β*] + [*αβ*])), we perform a quantitative and robust analysis on agarose gels (see Supplementary Figure S3).

**Fig. 2:**
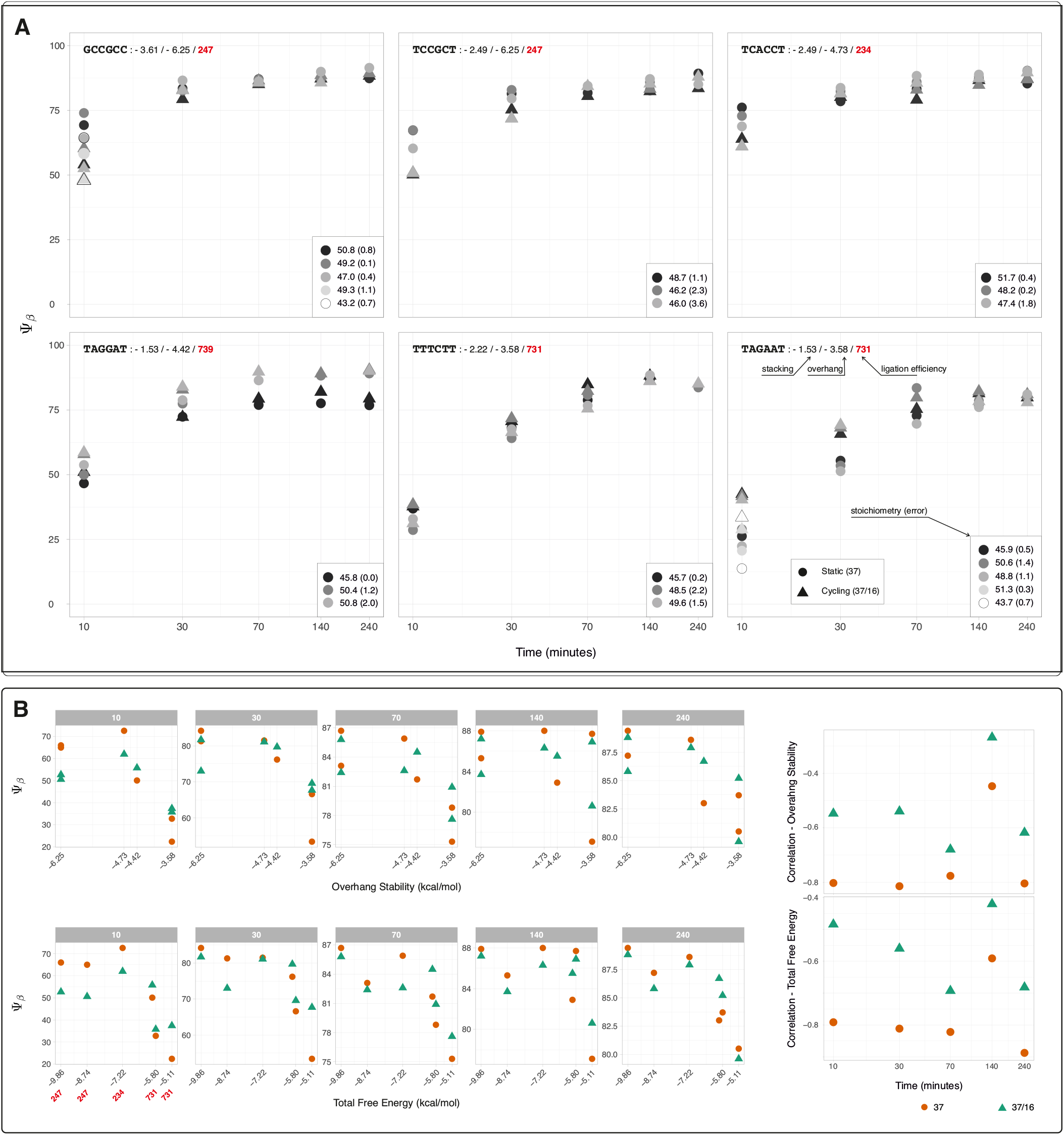
**A**. The percentage of product formed, Ψ_*β*_, as a function of time, was deduced from gel assays with [*αα*′] = 21 nM and [*β*′*β*] = 7 nM. The time points measured were 10, 30, 70, 140, and 240 minutes. Experiments were performed for different sequences and under various GGA assay conditions. Static assay (constant temperature of 37°C), data points are represented by circles; cycling assay (alternating between 37°C and 16°C every 5 minutes), upper triangles. For each sequence, stacking energies and overhang stabilities (in kcal/mol) are provided (e.g., -3.61 and -6.25 for `GCCGCC`, respectively). Ligation efficiencies (e.g., 247 for `CCGC`) deduced from sequencing-based assays are also shown (9). The insets display *S*_*β*_ values, reporting on the stoichiometry (the value in brackets is the corresponding error). The expected value of *S*_*β*_ is 49.7 (see Materials and Methods). **B**. Left panel: Using the same data as in A, we plot averaged Ψ_*β*_ values as a function of the overhang stability (upper sub-panel) or the total free energy (lower sub-panel). Each sub-panel contains a total of 6 plots corresponding to data obtained at 10, 30, 70, 140, and 240 minutes. Also shown are normalized ligation efficiencies as determined by sequencing-based assays (red). Right panel: Pearson correlation coefficients calculated for averaged Ψ_*β*_ values at different time points, evaluated for both overhang stability and total free energy. For all panels, circles refer to static assays, and upper triangles refer to cycling assays as in A.

Figure 2.A reports the percentage Ψ_*β*_ of product formed (*αβ*) over the course of the experiment (five time points up to 240 minutes) for all six overhangs under study: `GCCGCC`, `TCCGCT`, `TCACCT`, `TAGGAT`, `TTTCTT` and `TAGAAT`. Here, we have defined Ψ_*β*_ as:

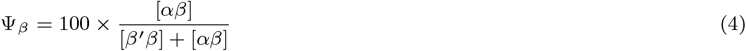

where brackets indicate molar concentrations. In addition, we report the initial stoichiometry *S*_*β*_, as defined in Materials and Methods under the paragraph Gel assays and analysis. To accurately determine the concentrations of products and reactants ([*αβ*], [*β*′*β*] and [*αβ*]), we perform agarose gel electrophoresis combined with an automated and quantitative analysis (Supplementary Figure S3). Ψ_*β*_ values represented as a function of time, overhang, and cycling in Figure 2.A can be displayed as in Figure 2.B. Here, at given time, we plot averaged Ψ_*β*_ (over three or five replicates) as a function of the stability of the overhangs (determined from nucleotides in positions 2, 3, 4 and 5, Supplementary Table S1) or the total energy (determined from nucleotides in positions 1 to 6, Supplementary Tables S1 and S2). At a given time, we first observe that the percentage of product formed increases with the stability of the overhangs: `CCGC` or `CACC` overhangs (energy of -6.25 and -4.73 kcal/mol, Table S1) are far more efficient than `AGAA` (−3.58 kcal/mol) and allow to reach high Ψ_*β*_ values within 10 minutes. This trend is observed at all time points (Figure 2.B, Supplementary Tables S3 and S4) for both the static and cycling assays. To further quantify this trend, we calculated Pearson correlation coefficients (shown in Figure 2.B, right panels) at each time point between Ψ_*β*_ and the overhang stability (upper panel) and the total free energy (lower panel). At 37°C, we observe significant (Supplementary Table S5) and high correlations (absolute values in excess of 0.8) between Ψ_*β*_ and the overhang stability (or the total free energy that accounts for stacking energies). For cycling experiments, the correlation is lower but non negligible (absolute values of ∼0.6). Most probably, lowering the temperature (as done in a cycling assay) prevents us from observing temperature-dependent characteristics (such as the stability of overhangs). These are the expected results when performing a simple ligation at low temperature (e.g. 16°C or 4°C) as recommended in ligation protocols. For comparison, we performed a statistical analysis on results obtained by high throughput assays (9) (Supplementary Figure S4). There, experiments have always been performed with temperature cycling, as recommended by GGA protocols for assemblies of 5 fragments and above, and show either unexpected correlation. However, it is important to note that those experiments were conducted at much lower substrate concentrations (below 1 nM), which may account for the observed differences. As seen in Figure 2.B, the difference in Ψ_*β*_ is time-dependent and pronounced at short times (30 minutes and less). In contrast, for longer reaction times, all overhangs show comparable Ψ_*β*_ (albeit still different and showing a significant correlation for static assays, Figure 2.B and Supplementary Tables S3 and S4). After 240 minutes, we obtain Ψ_*β*_ values of 80% to 90% for all overhangs, which is consistent with time-dependent GGA experiments of (6). This behavior is expected as the last reaction is irreversible (16) and so, for any overhang, Ψ_*β*_ values shall reach ∼100% when the reaction approaches completion. In other words, this reaction achieves a thermodynamic equilibrium where the ratio of on-rates to off-rates approaches infinity and it remains to be determined why ligation efficiencies values as reported by high throughput assays differ by up to 300% (9) after 300 minutes.

For strong overhangs, Figure 2 shows that static assays should be favored over cycling assays. In other words, for those overhangs, the loss in T4 DNA ligase activity at 16°C (and the complete loss of restriction endonuclease activity) is far more prejudicial than the possible gain in the dissociation constant when decreasing temperature. Here, this observation suggests that, for strong overhangs at the chosen DNA concentrations (21 nM for *αα*′ and of 7 nM for *β*′*β*), ligation should be performed at 37°C (as expected from the temperature-dependence activity of T4 DNA ligase (10; 17)). Finally, we observe that stacking energies may play a role for weak overhangs: while `TTTCTT` and `TAGAAT` have exactly the same overhang stability (as `TTCT` and `AGAA` are complementary to each other and not discernible in sequencing-based assays), `TAGAAT` that has a much weaker stacking energy (−1.53 as compared to -2.22 kcal/mol, Supplementary Table S2) and is more inefficient in a static assay. This result is in agreement with recent experiments (18) that have shown that base stacking affects the rate of ligation.

### Kinetic model

In the previous section, we demonstrated that there are experimental conditions where overhang stability (as determined by thermodynamics) strongly correlates with ligation efficiencies (as determined by Ψ_*β*_). Given the number of parameters that govern assembly, it is difficult, if not impossible, to predict the extent to which DNA stability (the opposite of DNA melting) affects GGA assembly. We therefore present in this section a kinetic model (Figure 3.A) that is able to capture experimental trends observed in Figure 2 and in Figure 3.C. We then extend our analysis to DNA molarities beyond the range feasible with conventional gel assays. Utilizing molarities typical of high-throughput assays (below 1 nM) and standard GGA protocols (5 nM and below), the model reveals that strong overhangs produce significantly higher ligation efficiencies compared to their non-stable counterparts.

**Fig. 3:**
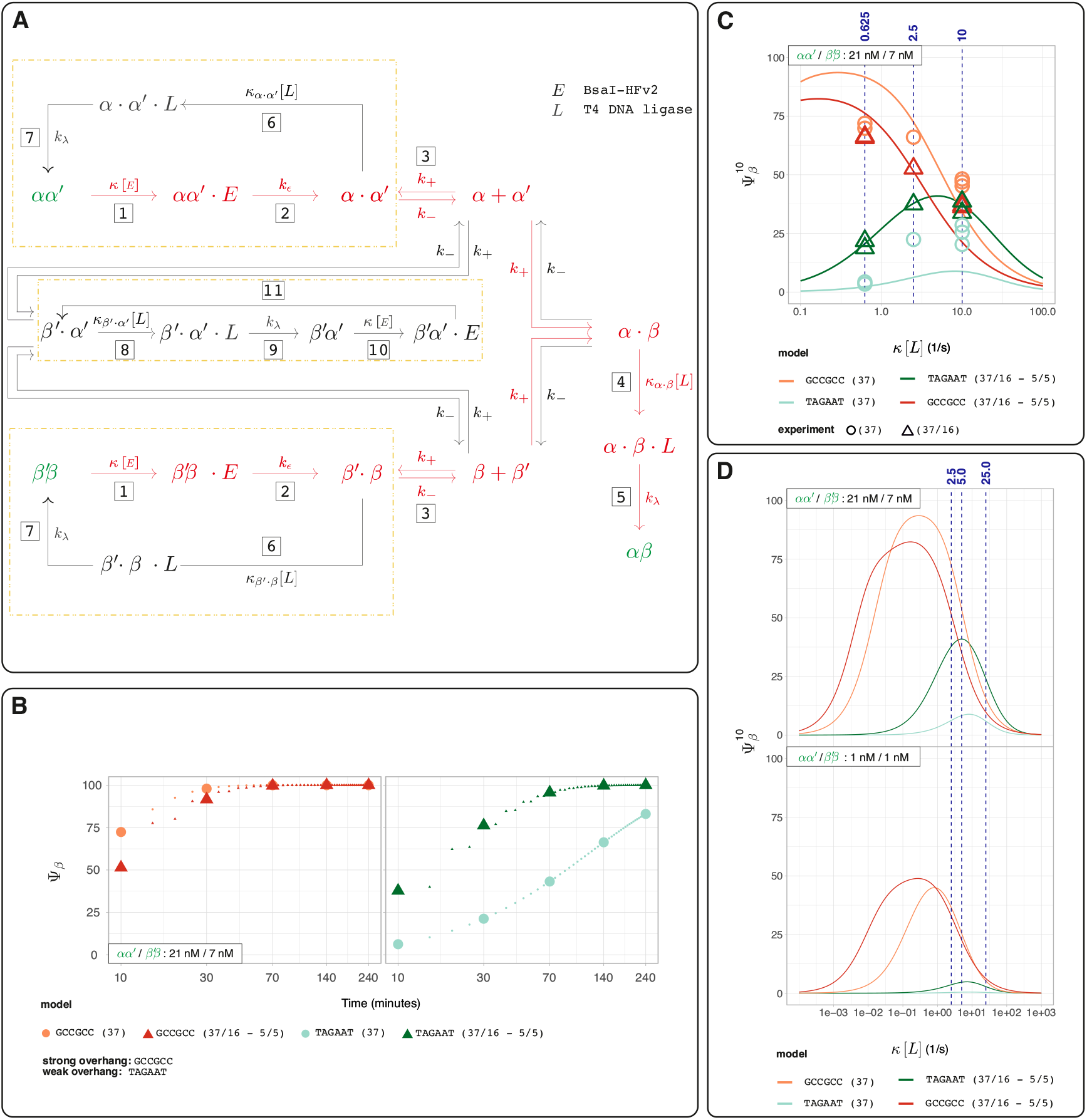
Kinetic model for our GGA assays. **A**. Initial reactants (*αα*′ and *β*′*β*) as well as the final (stable) product (*αβ*) are highlighted in green. Red arrows show the most direct route to form *αβ*. Enzymatic reactions are considered irreversible and are assumed to follow a two-steps Michaelis Menten process involving the binding of the enzyme to the substrate (represented by a centered dot) and a catalytic step (*k*_*ϵ*_, *k*_*λ*_). To account for a possible difference in stacking energy among the different nicked DNA molecules (*α* · *β, α* · *α*′, *β*′ · *β* and *β*′ · *α*′), different second order-reaction rates have been introduced (e.g. *κ*_*α*·*β*_). The concentration of enzymes is supposed constant so that *κ*[*E*], *κ*_*α*·*β*_[*L*], *κ*_*α*·*α*_′ [*L*], *κ*_*β*_′_·*β*_ [*L*] and *κ*_*β*_′_·*α*_′ [*L*] can be considered as first-order rates. Numbers in bold are used to identify the different reaction steps, additional details are given in the main text. **B**. Corresponding calculations (time dependence of Ψ_*β*_) for a static (37°C) and a cycling assays (37°C for 5 minutes and then 16°C for 5 minutes) using parameters from Supplementary Tables S6 and S7. Large symbols indicate times at which product concentrations were experimentally determined (10, 30, 70, 140 and 240 minutes). Calculations were performed for both a strong overhang (`GCCGCC`) and a weak overhang (`TAGAAT`). **C**. The dependence of Ψ_*β*_ as a function of the ligase concentration at 10 minutes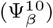. Solid lines represent calculated 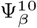 values (performed at [*αα*′] = 21 nM, [*β*′*β*] = 7 nM) for a weak (`TAGAAT`) and a strong overhang (`GCCGCC`) overhang while cycling the temperature (5 minutes at 37°C, 5 minutes at 16°C) or not (fixed temperature of 37°C). As in B, parameters given in Supplementary Table S6 and S7 were used. Symbols denote experimental points, and their colors match the simulations (e.g., orange circles are experimental points performed for a strong overhang at a static temperature of 37°C). Triangles represent cycling experiments (5 minutes at 37°C, 5 minutes at 16°C), whereas circles represent static experiments (fixed temperature of 37°C). Experimental points at *κ*[*L*]=2.5 *s*^−1^ are averaged values obtained from the data shown in Figure 2.A. Here a value of 2.5 *s*^−1^ for *κ*[*L*] corresponds to a final ligase concentration of 12 U/*µL*. **D**. 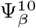as a function of *κ*[*L*] for different *αα*′ and *β*′*β* concentrations. Top panel: [*αα*′] = 21 nM, [*β*′*β*] = 7 nM, bottom panel: [*αα*′] = 1 nM, [*β*′*β*] = 1 nM (cycling assay : 37°C for 5 minutes and then 16°C for 5 minutes). Parameters for calculations as in B and C.

Importantly, this model is derived for one substrate at a time and ignores any competitive effects that may arise in presence of large number of competitive partners. Specifically, our model (Figure 3.A) assumes that enzymatic reactions follow the Michaelis–Menten kinetics, involving a binding step between the enzyme and the substrate (represented by a centered dot) and a catalytic step. At *t* = 0, the following species have non zero-molarities: *αα*′, *β*′*β, E* (endonuclease) and *L* (ligase). The formation of the final product *αβ* involves the binding of *α* and *β* (association rate *k*_+_, dissociation rate *k*_−_) and so the restriction endonuclease needs first to bind (rate constant *κ*[*E*]) and cleave (rate constant *k*_*ϵ*_) DNA (steps 1 and 2 for both *αα*′ and *β*′*β*). Ligase can then bind (rate constant *κ*_*α*·*β*_[*L*] for the nicked *α* · *β* intermediate) and form appropriate phosphodiester bonds (rate constant *k*_*λ*_). This results in joining *α* and *β* (steps 3 and 4) and in the formation of *αβ*. The sequence involving steps 1, 2, 3, 4, and 5 is the most direct route (highlighted in red in Figure 3) to form the target product, but other routes exist that may alter the overall kinetics. In particular, the hydrolysis of DNA fragments yields non-melted overhangs (steps 2 or 11) that can readily re-ligate (steps 6 or 8, dotted rectangles). It is challenging to estimate to what extent this affects the kinetics, as it depends on the concentration of the ligase, the molarities of the substrates, and the values of on- and off-rates (*i*.*e*., the stability of overhangs). Nonetheless, strong overhangs are slow-melting substrates with possibly lower off-rate values, making them more prone to re-ligation. Regarding the number of free parameters in the model, we have made several assumptions, which are detailed below:

i. *Irreversible Reactions*. We assume that all reactions catalyzed by enzymes are irreversible.
ii. *First-order rate constants*. First-order (*k*_*λ*_ and *k*_*ϵ*_) rate constants were chosen to be 10 s^−1^, in agreement with turn-overs reported in most enzymes (19).
iii. *Second-order rate constants*. We assume *κ* values not to depend on DNA substrates (which is an approximation as different substrates have different stacking energies that can possibly influence binding of ligase to the nicked substrate). This implies that *κ*_*α*·*β*_[*L*], *κ*_*α*·*α*_′ [*L*], *κ*_*β*_′_·*β*_ [*L*] and *κ*_*β*_′_·*α*_′ [*L*] (binding of the ligase *L*) are considered as first-order reaction rates. We furthermore impose that these *κ* values are substrate independent. In the Supplementary Text (Supplementary Figure S5), we show experiments that support this assumption. Similarly, we assume *κ*[*E*] (binding of the restriction endonuclease *E*) to be a first-order reaction rate.
iv. *On and off-rates constants*. The ratio of off-rates to on-rates has been chosen so that their ratio matches dissociation contants values at both 37°C and 16°C (see Supplementary Table S7). As a first approximation and to decrease the number of free parameters, we have further assumed that off- and on-rates are both substrate (e.g. *α* · *α*′ or *β*′ · *β*) and temperature independent. In the Supplementary Text, we present data obtained without imposing this constraint on the temperature (Supplementary Table S8 and Supplementary Figures S6, S7 and S8) that do not alter our conclusions.
v. *Enzyme concentrations*. We have assumed the concentration of enzymes to be constant. In other words, the concentration of enzymes is supposed to be much larger than substrate concentrations. In Supplementary Figure S9, we show that the above assumption holds.
vi. *Enzyme activity and temperature*. We finally assume the activity of enzymes to be temperature dependent with a change in activity of 100 % (60 %) for the restriction endonuclease (ligase) at 16°C (10; 17).

Considering the above assumptions, the remaining variables in our model are *κ*[*L*] and *κ*[*E*], along with the absolute values of the on- and off-rates (the ratio between these rates is already defined by free energy considerations). The model is expected to capture the trends observed in Figure 2 for both a strong (`GCCGCC`) and a weak (`TAGAAT`) overhang, whether cycling the temperature or not (see Figure 3.B, which reports calculated Ψ_*β*_ as a function of time). We further refined the model by fitting it to experiments conducted across a large range of ligase concentrations at 10 minutes, varying the concentration by a factor of 16 (as shown in Figure 3.C). Because GGA is typically performed with temperature cycling (as in (9) or when using the E1601 kit from NEB for assemblies containing five or more fragments), we prioritized experimental data obtained at 37°C/16°C when adjusting free parameters. Values of parameters are given in Supplementary Tables S6 and S7. We note finally that Ψ_*β*_at 10 minutes is sufficient to predict product formation efficiency, and that it shows the greatest differences across all experimental time points (Figure 2.A).

As shown in Figure 3.B and Figure 3.C, the model successfully captures all experimental trends observed in Figure 2: (*i*) cycling seems to be prejudicial for strong overhangs, (*ii*) strong overhangs are more efficient than weaker ones, (*iii*) there is a marked difference in Ψ_*β*_ for weak overhangs for static and cycling assays, (*iv*) the impact of ligase concentration on Ψ_*β*_ varies significantly between strong and weak overhangs. The latter is to be expected: for a strong overhang, forming the *α* · *β* intermediate is challenging as these overhangs are slow-melting that can re-ligate (see step 6 in Figure 3.A). However, once the intermediate *α* · *β* is formed, it is less likely to dissociate, allowing *αβ* to eventually form as long as *k*_−_ is not significantly larger than *κ*_*αβ*_[*L*]. In contrast, a weak overhang is more likely to reach *α* · *β*, but in this case, decreasing the ligase concentration will favor dissociation (as here *k*_−_ is significantly larger than *κ*_*αβ*_[*L*]).

Adjusting parameters for different overhang stabilities, temperatures, and ligase concentrations imposes many constraints, giving us confidence that the model can predict trends at molarities we have not yet explored due to the sensitivity of gel assays. Experiments shown in Figure 2 and Figure 3.C were conducted at molarities of 21 nM for *αα*′ and of 7 nM for *β*′*β*. These values significantly differ from those used in high-throughput assays, where the molarity of individual overhangs was lower than 1 nM (the total concentration of DNA substrate was 100 nM (9)). As discussed in the Supplementary Text, the concentration of T4 DNA ligase and BsaI-HFv2 is not specified in (9) as a kit was used for those multiplexed assays. However, we can assume the endonuclease concentration is similar to ours, and that the ligase concentration is approximately twice the concentration we used (see Supplementary Discussion). As shown in Figure 3.C, where Ψ_*β*_ is calculated at 10 minutes, increasing the ligase concentration significantly impacts the results. Under these conditions, during cycling, a strong overhang (`CCGC`) exhibits lower Ψ_*β*_ than a weak overhang (`AGAA`). We may then wonder if this effect is observed at other molarities, particularly those used in high-throughput experiments (lower than 1 nM). There, our model (Figure 3.D) predicts that, when cycling, strong overhangs yield higher GGA assembly (as evidenced by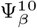) when *κ*[*L*] is increased by up to a factor 10. This result contradicts observations from high-throughput assays, which show that overhang stability inversely correlates with ligation efficiency (Supplementary Figure S4). One possible explanation is that the presence of competitive partners in high-throughput GGA experiments dramatically affects the reported ligation efficiencies. To test this hypothesis, we present in the following section an assembly of 10 fragments following the protocols recommended for GGA by NEB. There, the molarity of fragments is expected to be 2.5 nM (if using pre-cloned sequences) or 5 nM (if using PCR fragments). At those molarities and given simulations performed using a cycling period of 2 or 10 minutes, different molarities and different parameters, we expect strong overhangs to yield higher GGA efficiencies (Supplementary Figures S7, S8, S10 and S11) in assays involving the presence of a defined number of fragments exhibiting high fidelities (*i*.*e*., in conditions where mismatch ligations do not occur).

### 10 fragments GGA assembly

To test our model, which proposes that strong overhangs facilitate the rapid formation of GGA assemblies, we now present a GGA assembly involving 10 fragments. The key parameter in this analysis is fidelity, which refers to an overhang’s ability to avoid binding with additional partners. Low fidelity, as mathematically defined in the Materials and Methods section, can result in truncated assemblies (containing fewer than 10 fragments) or assemblies with fragments arranged incorrectly in terms of order and/or direction. Based on previous sequencing-based experiments, we have carefully chosen 9 overhangs with a typical stability of 4.5 kcal/mol or higher (absolute values) that exhibit a fidelity of ∼1 in a static or a cycling assay. We define this set as optimal. Additionally, we have selected a second set of overhangs without imposing any stability restrictions (see Materials and Methods). This second set, defined as non-optimal, also possesses a fidelity of ∼1 (see Table 1). When comparing both sets, we observe that the overhangs in the optimal set exhibit significantly lower ligation efficiencies, as determined by high-throughput assays (Figure 4.A). Consequently, the optimal set should result in a reduced number of full-length fragments compared to the non-optimal sets. We note finally that both assemblies exhibit high fidelities and are thus representative of the constraints (absence of competitive partners yielding mismatch ligations) imposed in our simulations.

**Table 1.**
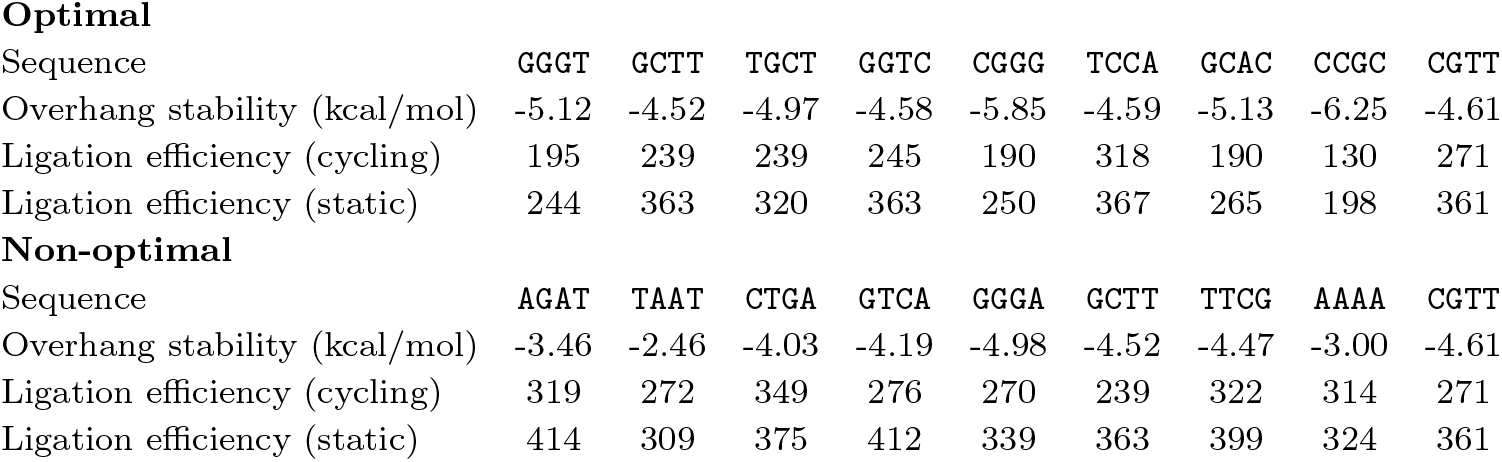
The chosen overhangs were selected for both optimal and non-optimal sets. Optimal overhangs had a stability of 4.5 kcal/mol or higher (in absolute values), while non-optimal sets had no restrictions on stability. Also shown are normalized ligation counts as provided by the NEBridge Ligase Fidelity Viewer from NEB for a static (fixed temperature of 37°C) and a cycling assay (37°C for 5 minutes and 16°C for 5 minutes) in the presence of BsaI-HFv2 and T4 DNA ligase (8). For both sets and temperature assays, the NEB Viewer reports fidelities equal or greater than 0.99 when using the pGGAselect vector. See also Supplementary Figure S16, showing the dispersion of ligation counts in a graph (similar to Figure 4.C).

**Optimal**
| Sequence | GGGT | GCTT | TGCT | GGTC | CGGG | TCCA | GCAC | CCGC | CGTT |
| --- | --- | --- | --- | --- | --- | --- | --- | --- | --- |
| Overhang stability (kcal/mol) | -5.12 | -4.52 | -4.97 | -4.58 | -5.85 | -4.59 | -5.13 | -6.25 | -4.61 |
| Ligation efficiency (cycling) | 195 | 239 | 239 | 245 | 190 | 318 | 190 | 130 | 271 |
| Ligation efficiency (static) | 244 | 363 | 320 | 363 | 250 | 367 | 265 | 198 | 361 |

**Table 1.** The chosen overhangs were selected for both optimal and non-optimal sets.
| Sequence | AGAT | TAAT | CTGA | GTCA | GGGA | GCTT | TTCG | AAAA | CGTT |
| --- | --- | --- | --- | --- | --- | --- | --- | --- | --- |
| Overhang stability (kcal/mol) | -3.46 | -2.46 | -4.03 | -4.19 | -4.98 | -4.52 | -4.47 | -3.00 | -4.61 |
| Ligation efficiency (cycling) | 319 | 272 | 349 | 276 | 270 | 239 | 322 | 314 | 271 |
| Ligation efficiency (static) | 414 | 309 | 375 | 412 | 339 | 363 | 399 | 324 | 361 |

**Fig. 4:**
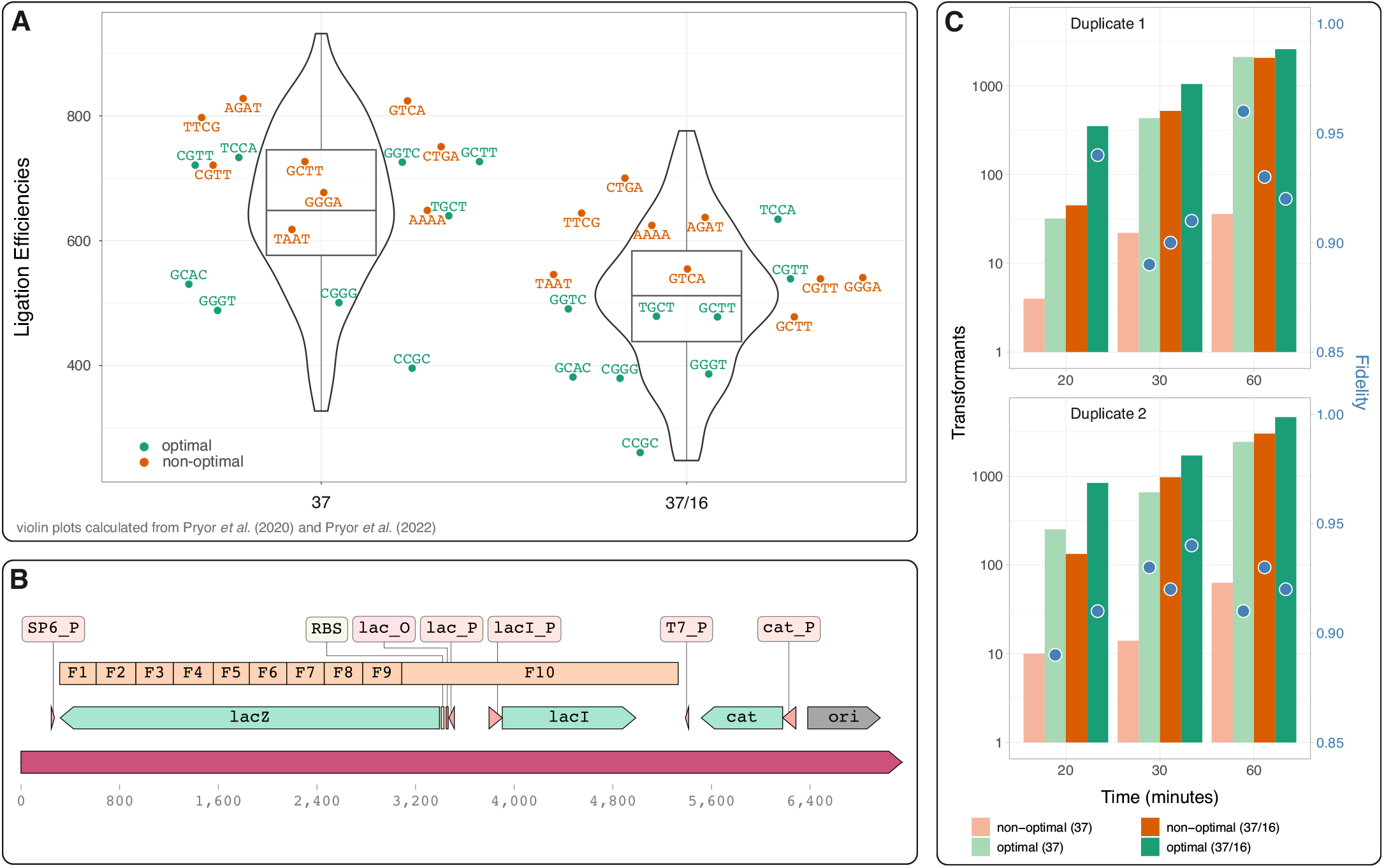
**A**. Violin plots (*i*.*e*., kernel density estimation) showing ligation efficiencies for a static (37°C) and a cycling (37/16°C, which corresponds to 5 minutes at 37°C and 5 minutes at 16°C) GGA assay. Data represents 120 diagonal elements of the 256 × 256 element matrix as determined by sequencing experiments (4) and (9). As in (9), reported values are normalized to 100,000 ligation events. Circles represent individual overhangs chosen to reconstitute the lac cassette operon into pGGAselect. Green circles indicate overhangs of the optimal set, while orange circles represent the non-optimal set. **B**. Diagram of the lac operon cassette (cloned from the pNJ2075 plasmid) reconstituted by the 10 fragments GGA assembly (named F1 to F10). The length of the fragments corresponds to that of the optimal set, with fragments of the non-optimal set being approximately the same size (Supplementary Table S10 and Supplementary Figure S14). Note that lacI is dispensable for lacZ production, thus the last fragment (F10) is much longer than the other fragments. The suffix `P` stands for promoter, `O` for operator, and `cat` stands for chloramphenicol acetyltransferase. **C**. Top panel: The number of transformants over time was measured for two assemblies performed under identical conditions, except for slight variations in transformation conditions (detailed in Supplementary Protocols). Transformant counts are normalized to the total outgrowth volume and are presented for both optimal and non-optimal conditions in two assays: a static assay (constant temperature of 37°C) and a cycling assay (alternating the temperature between 37°C and 16°C every 1 minute, as recommended in NEB protocols). Additionally, fidelity values, defined as the ratio of correct transformants to the total number of transformants, are presented. In assays where the number of transformants is low, fidelity values exhibit significant fluctuations and are not shown on this graph (see Supplementary Tables S11 and S12 for detailed values).

To test for the assembly, we have chosen an approach similar to that of (9), which essentially involves cloning a lac operon cassette into a vector (pGGAselect): in a medium supplemented with IPTG and X-Gal, transformants with the correct assembly (lac cassette, see Figure 4.B) will show a blue phenotype while those having errors (wrong number of fragments and/or wrong orientation or order and/or unrelated seqeunces) will show a white phenotype. For those experiments, the recommended protocol for the NEBridge Golden Gate Assembly Kit (BsaI-HFv2) (E1601) was followed as described in the Supplementary Information. This protocol involves cycling the temperature every 1 minute for a total of 60 minutes and using a total volume of NEB Golden Gate Assembly Mix of 0.5 *µ*L per 10 *µ*L. Furthermore, we conducted an assembly at a constant temperature of 37°C, as our model predicts that a static assay can achieve comparable efficiencies.

Figure 4.C shows the total number of transformants as well as the fidelity (defined as the ratio of the total number of blue transformants to the total number of transformants), see Supplementary Table S11 and S12 for exact numbers. The results are shown for both the optimal and non-optimal sets, with and without temperature cycling. First, we observe that the optimal set (supposed to have overhangs with low ligation efficiencies) yields a higher number of transformants for both the cycling and static assays. This effect is more pronounced at early time points, similar to what is observed in experiments performed on single substrates (see Figure 2). Specifically, as the reaction approaches completion (at 60 minutes), both the optimal and non-optimal assays tend to reach identical values in the cycling assay. This is expected, as the final step in a GGA assay involves irreversible ligation. We also observe that performing the experiments at a static temperature is detrimental for the non-optimal set. This set contains weak overhangs (e.g. `TAAT` and `AAAA`, see Table 1), which significantly lower the yield. These results demonstrate that static experiments should be avoided when performing assemblies involving weak overhangs. Finally and as shown in Figure 4.C, both sets achieve high fidelities, slightly below the expected value of 0.99 (Table 1). This slight discrepancy can be attributed to potential sub-PCR products, as indicated by faint bands visible when the gel was over-exposed (Supplementary Figure S15). In conclusion, our model has successfully predicted the observed experimental trends when conducting GGA assembly on a defined number of fragments (here 10), demonstrating that strong overhangs with low competitive effects should be prioritized for optimal assembly efficiency.

## OPTIMIZING GGA USING STRONG OVERHANGS

High-throughput DNA sequencing has recently provided a comprehensive analysis of GGA. For these assays, strong overhangs (determined from thermodynamic data) exhibit lower ligation efficiencies compared to weaker ones (see Supplementary Figures S4 and Figure 5.A). While this seems counterintuitive, as ligase activity is crucial for GGA, the authors propose that strong overhangs are slow-melting, which increases the probability of self-religation (see Figure 3.A). While this may be a valid assumption, our simulations and experimental data (performed on a low number of substrates having high fidelities) indicate that under the conditions used in high-throughput assays or recommended GGA protocols, strong overhangs result in higher ligation efficiencies. This discrepancy indicates that high-throughput GGA results cannot be generalized to experiments involving a specified number of high-fidelity fragments, which is typically the case when performing GGA. Still, recommended protocols and tools (8) report ligation efficiencies for assemblies involving a significantly lower number of fragments

**Fig. 5:**
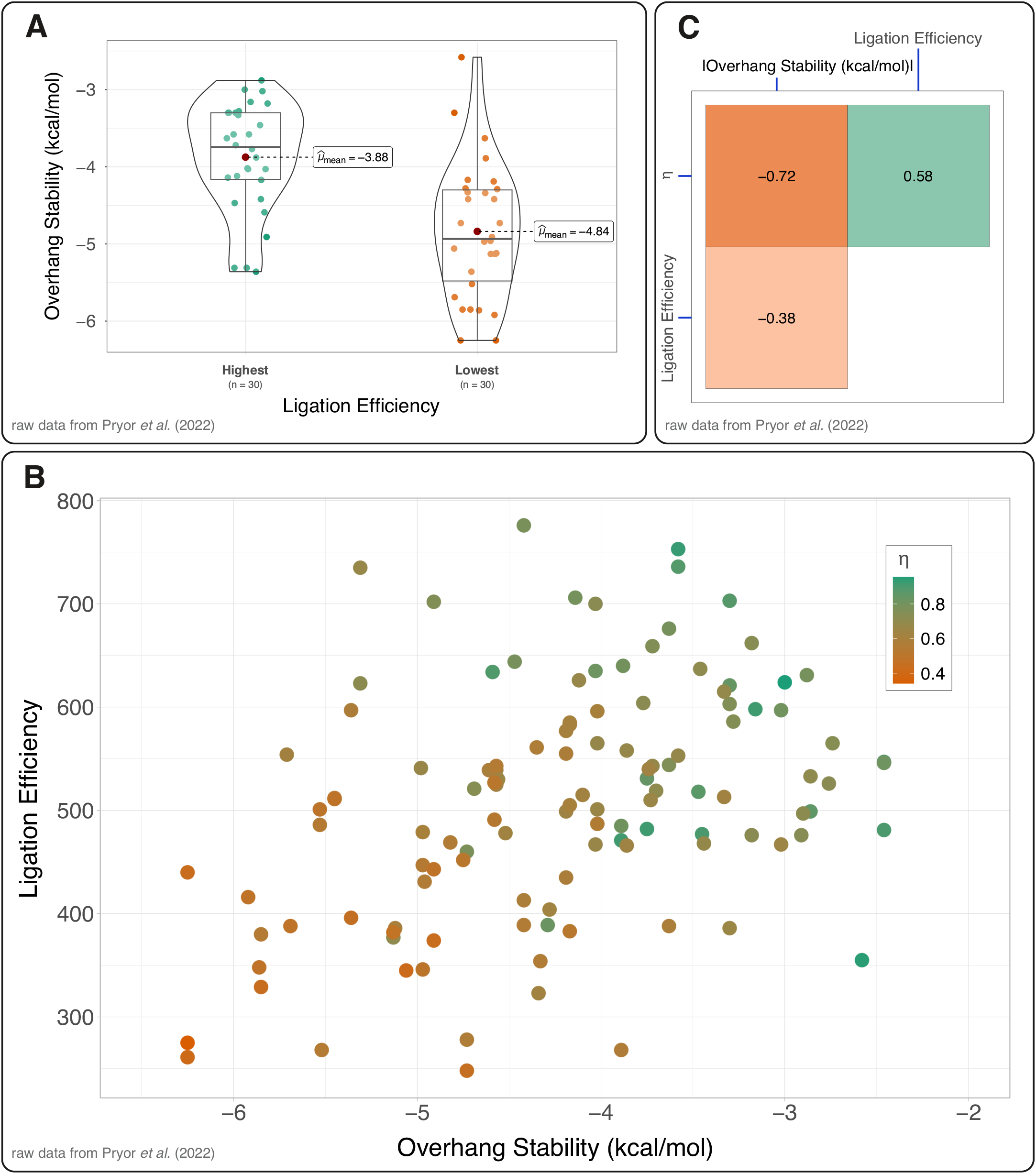
Analyzing high-throughput data from (4). **A**. Violin plots illustrating the distribution of ligation efficiencies achieved in the presence of T4 DNA ligase and BsaI-HFv2 during a cycling assay. The plots highlight the top 30 and bottom 30 ligation efficiency values. **B**. Ligation efficiency as a function of the overhang stability. For each point, we calculate *η*, defined as the ratio between the ligation efficiency (obtained from Watson-Crick base pairing) and total ligation count (including mismatches) for a given overhang. **C**. Correlation matrix showing Pearson correlation coefficients for *η*, ligation efficiency and overhang stability (e.g., the correlation between *η* and the ligation efficiency is 0.58).

Upon analyzing high-throughput GGA data, it becomes evident that these assays exhibit significant bias. In Figure 5.B, we introduce a new metric, *η*, defined as the ratio of the maximum count (*i*.*e*. ligation efficiency, associated with perfect Watson-Crick base pairing) to the total ligation count for a given overhang. Additionally, we calculate the correlation coefficients between *η*, ligation efficiency, and overhang stability (Figure 5.C). Notably, strong overhangs tend to have low ligation efficiencies and low *η* values. In other words, a strong overhang exhibits lower ligation efficiencies due to a higher occurrence of mismatches. A more subtle yet important point is the following: given that that phosphodiester bond formation is irreversible, one would expect overhangs to show comparable efficiencies at completion. Surprisingly, reported ligation efficiencies vary by 300% or more after 300 minutes. Either those experiments report on values that cannot represent end-time points or other effects (e.g. competitive inhibition-like effects arising from simultaneous mismatch ligations) have to be taken into account. In any case, normalizing to 100,000 events as in (9) seems to be incorrect. Additionally and given that ligation efficiencies are biased, it is crucial to determine if ligation fidelities are also affected. Finally, it remains to be seen if similar high-throughput data obtained in the presence of DNA ligases alone exhibit the same bias (20). For instance, using the Supplementary data from (20) (Table S4), we can estimate the correlation between initial velocities and GC content of hLig3 for substrates in isolation. We find a value of 0.69, which indicates a significant correlation. Using the data obtained by high-throughput assays (File S1 in (20)), we find a value of -0.39. Interestingly, among the ligases studied (T3 DNA ligase, T4 DNA ligase, T7 DNA ligase, hLig3 and PBCV-1 ligase), hLig3 has the lowest overall fidelity and so may be subjected to the strongest competitive effects, leading to potential biases when experiments are performed in the presence of all competitive partners. Data (absolute ligation counts) derived from high-throughput experiments (including both GGA and ligation assays) should be interpreted with great caution when extrapolated to practical assays. However, we acknowledge that those experiments have clearly benefited the development and optimization of GGA, particularly in enhancing the efficiency and reliability of large assemblies (4; 8) and/or facilitating reverse genetics (21).

Based on our data, we recommend using strong overhangs - those with binding energies of 4.5 kcal/mol or higher (in absolute terms) - for GGA. Implementing this approach, particularly when following the protocols provided by NEB, ensures a rapid and reliable assembly process. This strategy optimizes the efficiency and consistency of GGA, making it a robust choice for complex molecular cloning tasks. However, it is important to note that following NEB guidelines when assembling a low number of fragments and selecting overhangs, which are supposed to yield high efficiencies according to high-throughput GGA assays, may be detrimental. Specifically, the protocols suggest performing experiments at a static temperature of 37°C and we have demonstrated (Figure 4.C) that this approach results in poor yields.

## SUPPLEMENTARY DATA

Supplementary Data is available.

## Author contributions statement

W.G. planned the experiments. P.S., N.J., E.H., A.K., G.M.C, F.B. and W.G. performed GGA assays and/or contributed to molecular biology assays. P.S., F.B. and W.G. analyzed GGA data. F.B., P.H. and W.G. derived the kinetic model and W.G. performed calculations.

N.J. conducted 10 fragments GGA assembly and contributed to the preparation and analysis of the assays. W.G. performed statistical analysis and wrote the paper. All authors read the manuscript and contributed to comments or corrections.

## Acknowledgments

We thank Marie-Nöelle Lalloz-Vogel (IPCMS, Strasbourg) for her help with preparing the DNA substrates and Patricia Moussounda (IJM, Paris) for her help with media preparation.

This research was funded by the National Science Centre, Poland (grant SONATINA 2017/24/C/NZ1/00456 to G.M.C) and was supported by a French Government Scholarship (Campus France to P.S.). N.J. is supported by a grant from the Fondation ARC pour la recherche sur le cancer (ARCPJA2022050005002) and a grant from Université Paris-Cité IDEx https://u-paris.fr; ANR-18-IDEX-0001, Emergence en Recherche, RS30J23IDX64 KATAREP).

This study contributes to the IdEx Université Paris Cité (ANR-18-IDEX-0001).

## References

1. W Szybalski, SC Kim, N Hasan, and AJ Podhajska. Class-iis restriction enzymes–a review. Gene, 100:13–26, 1991. doi: 10.1016/0378-1119(91)90345-c.

2. A Pingoud and A Jeltsch. Structure and function of type ii restriction endonucleases. Nucleic Acids Res, 29(18):3705–27, 2001. doi: 10.1093/nar/29.18.3705.

3. Carola Engler, Romy Kandzia, and Sylvestre Marillonnet. A one pot, one step, precision cloning method with high throughput capability. PLoS One, 3(11):e3647, 2008. doi: 10.1371/journal.pone.0003647.

4. John M Pryor, Vladimir Potapov, Katharina Bilotti, Nilisha Pokhrel, and Gregory J S Lohman. Rapid 40 kb genome construction from 52 parts through data-optimized assembly design. ACS Synth Biol, 11(6):2036–2042, 2022. doi: 10.1021/acssynbio.1c00525.

5. Yingjia Tong, Jingwen Zhou, Liang Zhang, and Peng Xu. A golden-gate based cloning toolkit to build violacein pathway libraries in yarrowia lipolytica. ACS Synth Biol, 10(1):115–124, 2021. doi: 10.1021/acssynbio.0c00469.

6. Nicholas A W Bell and Justin E Molloy. Efficient golden gate assembly of dna constructs for single molecule force spectroscopy and imaging. Nucleic Acids Res, 50(13):e77, 2022. doi: 10.1093/nar/gkac300.

7. Sylvestre Marillonnet and Ramona Grützner. Synthetic dna assembly using golden gate cloning and the hierarchical modular cloning pipeline. Current protocols in molecular biology, 130(1):e115, 2020.

8. Andrew P. Sikkema, S. Kasra Tabatabaei, Yan-Jiun Lee, Sean Lund, and Gregory J. S. Lohman. High-complexity one-pot golden gate assembly. Current Protocols, 3(9):e882, 2023. doi: 10.1002/cpz1.882. URL https://currentprotocols.onlinelibrary.wiley.com/doi/abs/10.1002/cpz1.882.

9. John M Pryor, Vladimir Potapov, Rebecca B Kucera, Katharina Bilotti, Eric J Cantor, and Gregory JS Lohman. Enabling one-pot golden gate assemblies of unprecedented complexity using data-optimized assembly design. PLoS One, 15(9):e0238592, 2020. doi: 10.1371/journal.pone.0238592.

10. B Weiss, A Jacquemin-Sablon, TR Live, GC Fareed, and CC Richardson. Enzymatic breakage and joining of deoxyribonucleic acid. vi. further purification and properties of polynucleotide ligase from escherichia coli infected with bacteriophage t4. J Biol Chem, 243 (17):4543–55, 1968.

11. IR Lehman. Dna ligase: structure, mechanism, and function. Science, 186(4166):790–7, 1974. doi: 10.1126/science.186.4166.790.

12. Mohammad HamediRad, Scott Weisberg, Ran Chao, Jiazhang Lian, and Huimin Zhao. Highly efficient single-pot scarless golden gate assembly. ACS Synth Biol, 8(5):1047–1054, 2019. doi: 10.1021/acssynbio.8b00480.

13. Indrajeet Patil. Visualizations with statistical details: The’ggstatsplot’approach. Journal of Open Source Software, 6(61):3167, 2021.

14. J SantaLucia, Jr. A unified view of polymer, dumbbell, and oligonucleotide dna nearest-neighbor thermodynamics. Proc Natl Acad Sci U S A, 95(4):1460–5, 1998. doi: 10.1073/pnas.95.4.1460.

15. Ekaterina Protozanova, Peter Yakovchuk, and Maxim D Frank-Kamenetskii. Stacked-unstacked equilibrium at the nick site of dna. J Mol Biol, 342(3):775–85, 2004. doi: 10.1016/j.jmb.2004.07.075.

16. Gregory J S Lohman, Lixin Chen, and Thomas C Evans, Jr. Kinetic characterization of single strand break ligation in duplex dna by t4 dna ligase. J Biol Chem, 286(51):44187–44196, 2011. doi: 10.1074/jbc.M111.284992.

17. FM Pohl, R Thomae, and A Karst. Temperature dependence of the activity of dna-modifying enzymes: endonucleases and dna ligase. Eur J Biochem, 123(1):141–52, 1982. doi: 10.1111/j.1432-1033.1982.tb06510.x.

18. Jibin Abraham Punnoose, Kevin J Thomas, Arun Richard Chandrasekaran, Javier Vilcapoma, Andrew Hayden, Kacey Kilpatrick, Sweta Vangaveti, Alan Chen, Thomas Banco, and Ken Halvorsen. High-throughput single-molecule quantification of individual base stacking energies in nucleic acids. Nature Communications, 14(1):631, 2023.

19. Arren Bar-Even, Elad Noor, Yonatan Savir, Wolfram Liebermeister, Dan Davidi, Dan S Tawfik, and Ron Milo. The moderately efficient enzyme: evolutionary and physicochemical trends shaping enzyme parameters. Biochemistry, 50(21):4402–10, 2011. doi: 10.1021/bi2002289.

20. Katharina Bilotti, Vladimir Potapov, John M Pryor, Alexander T Duckworth, James L Keck, and Gregory J S Lohman. Mismatch discrimination and sequence bias during end-joining by dna ligases. Nucleic Acids Res, 50(8):4647–4658, 2022. doi: 10.1093/nar/gkac241.

21. Katharina Bilotti, Sarah Keep, A. Paul Skeldon, James M. Pryor, Sarah K. Taylor, and Erica Bickerton. One-pot golden gate assembly of an avian infectious bronchitis virus reverse genetics system. PLOS ONE, 19(1):e0307655, 2024. doi: 10.1371/journal.pone.0307655. URL https://journals.plos.org/plosone/article?id=10.1371/journal.pone.0307655.

