## Supplementary Information for "Enhanced Golden Gate Assembly: Evaluating Overhang Strength for Improved Ligation Efficiency"

### SUPPLEMENTARY DISCUSSION

#### Concentrations of enzymes

For high-throughput GGA fidelity and bias assays, as well as GGA of the lac cassette test systems, a kit was used for BsaI-HFv2 and T4 DNA ligase [1], so their exact concentrations are not known. However, experiments performed with other enzymes (Esp3I, BbsI-HF, and SapI) in the referenced study [1] utilized a final ligase concentration of 25 U/$\mu$L and a final endonuclease concentration of 0.75 U/$\mu$L. Additionally, in a separate study performed by NEB [2], GGA assemblies with 12 fragments were conducted using a final concentration of 25 U/$\mu$L for T4 DNA ligase and 0.75 U/$\mu$L for BsaI-HFv2. Finally, in their 24-Fragment Golden Gate Assembly using BsaI-HFv2 protocol, NEB recommends to use a final ligase concentration of 20 U/$\mu$L and a final endonuclease concentration of 0.6 U/$\mu$L for assemblies of 10 fragments of less. Therefore, we assume the endonuclease concentration in our experiments is similar, around 0.6 U/$\mu$L or 0.65 U/$\mu$L, to that used in high-throughput GGA assays. In contrast, our ligase concentration of 12 U/$\mu$L is significantly lower than the 25 U/$\mu$L reported in study [1].

Importantly, we employ molarities much higher than those used in sequencing-based GGA assays (approximately 0.5 nM per overhang) because we perform the analysis using standard gel assays, which can detect only a few tens of nano-grams of DNA per band. Reducing the concentration of DNA molecules from 21 nM to 0.5 nM while maintaining the same sensitivity achieved in this study would require increasing the reaction volume by approximately 40 times. Since we use about 1 $\mu$L of stock enzyme per reaction, we would need almost one tube of enzyme per experiment performed on a single overhang (see Supplementary Protocols, page 11 of this document). Considering that these assays were performed in triplicates for six different overhangs, we would need about 20 tubes of enzymes to reproduce the data presented in Figure 2.

Moreover, taking into account other experimental data presented in this paper (such as those in Figure 3.C), and considering the optimization steps for gel assays as well as potential failed experiments, we may need more than 100 tubes each of endonuclease and ligase to reproduce all the data presented in this study. Additionally, working with larger volumes and maintaining optimal temperature control would require splitting an approximate $\sim$400 $\mu$L volume into 40 tubes. While feasible, this approach is suboptimal when collecting data points at specific time intervals. Therefore, we have decided to use molarities of 21 nM and 8 nM for $\alpha\alpha'$ and $\beta'\beta$, respectively, for our experiments.

#### Primers and sequence context

Type IIP enzymes, composed of symmetric domains, interact with DNA on both sides of the recognition site sequence [3]. Consequently, nucleotides flanking the recognition site at the 5' and 3' ends may modulate binding, potentially altering enzyme activity [4, 5]. In contrast, Type IIS enzymes have a different structure, comprising a recognition domain and a catalytic domain located downstream of the recognition site, linked by a connector [6]. Due to the non-symmetric nature of the protein, the sequence downstream of the recognition should not affect binding. To our knowledge, the role of the upstream sequence has never been discussed in the context of GGA (and not discussed in protocols involving endonucleases). For BsaI-HFv2, the presence of just a single base upstream of the recognition sequence is sufficient to achieve activity levels between 50% and 100% (NEB: Cleavage Close to the End of DNA Fragments). In the context of simple digestion assays, NEB does not specify any particular nucleotide bases as being required for optimal enzyme activity. This suggests that, at the enzyme concentrations typically used in experiments, the sequence context likely plays a minor role. In fact, when assessing enzyme activity based on the number of bases upstream of the recognition sequence, the first base is not consistently conserved (NEB: Cleavage Close to the End of DNA Fragments (oligonucleotides)). In GGA assays, NEB also does not require specific sequences for optimal performance. Indeed, in sequencing-based GGA assays, the substrate precursor sequences upstream of the recognition sites differ between the enzymes BsaI-HFv2, Esp3I, and BbsI-HF. This suggests that the authors may not have accounted for the potential impact of these variations [1] and did not enforce specific upstream sequences in practical assays. In other words, the authors assume that their results are broadly applicable to other sequences and that variations in these upstream sequences do not affect ligation efficiency or fidelity counts. Finally, it is important to note that sequences upstream of the recognition site are not conserved when using commercial vectors like pGGAselect or when designing new vectors for GGA [7].

Still we may wonder whether our conclusions hold if the binding rate for $\alpha\alpha'$ is significantly higher than that for $\beta'\beta$ (and *vice versa*). To account for this potential effect, we can adjust the value of $\kappa[E]$ for reactions involving $\alpha\alpha'$ and $\beta'\alpha'$. Similarly, if the upstream sequence of $\beta'\beta$ shows significantly higher binding affinity than $\alpha\alpha'$, we can increase the value of $\kappa[E]$ for reactions involving $\beta\beta'$ and $\beta'\alpha'$. As shown in Supplementary Figure S12, adjusting for changes in the binding rate (by a factor 10) does not affect our conclusions. In other words, even if the sequences $\alpha\alpha'$ and $\beta'\beta$ exhibit different binding efficiencies for the endonuclease, and we apply the derived fitting parameters for identical upstream sequences (as in sequencing-based assays), we do not observe evidence of major changes in the dependence of $\Psi_\beta$ and thus we conclude that sequencing-based experiments have inherent biases.

We selected the primers to mimic potential scenarios encountered when designing primers for GGA, where the upstream sequence may be adjusted to prevent the formation of secondary structures or primer dimers.

2 Fragments Assembly: For $\alpha\alpha'$ fragments, the upstream sequence is conserved across all primers as `ATAGTA` (see Supplementary Table S9 and the DNA Sequences section, page 20 of this document). Similarly, the upstream sequence for the $\beta'\beta$ fragments is conserved as `ATGCGA`.

10 Fragments Assembly: Each reverse primer producing an overhang, whether from the optimal or non-optimal set has either 4, 7 or 8 nucleotides upstream of the recognition sequence, comprising `AGTA`. Each forward primer producing an overhang, whether from the optimal or non-optimal set, has either 4, 7 or 8 nucleotides upstream of the recognition sequence, comprising `CGCA`. For instance, OL1475, which produces the overhang `ATCT`, has `AATAGTA` upstream of the recognition sequence, while OL1476, which produces the complementary overhang `AGAT`, has `ATACGCA` upstream of the recognition site. Note that all reverse primers have a 3' `G` base adjacent to the recognition site (except OL1465, 3' `A`) and that all forward primers have a 3' `A` base adjacent to the recognition site. Note finally that OL1465 produces the `CCCG` overhang, which is a strong overhang for which we do not observe significant changes in $\Psi_\beta$ upon varying the stacking energy (Figure 2 of the main text and Supplementary Figure S5).

#### Estimating errors in percentages

As seen in the main text and in Supplementary Figure S3, we have defined the percentage of product formed $\Psi_\beta$ as:

$$100 \times \left(1 + \left(\frac{\beta'\beta}{\alpha\beta}\right) \times \gamma\right)^{-1}$$

where $\beta'\beta$ and $\alpha\beta$ are intensities and $\gamma$ (~1.34) accounts for the difference in molecular weight for both fragments. Assuming independent random errors on $\beta'\beta$ ( $\delta_{\beta'\beta}$) and $\alpha\beta$ ( $\delta_{\alpha\beta}$), the error on the percentage $\delta_{\Psi_\beta}$ reads:

$$100 \times \frac{\gamma}{\alpha\beta \times (1 + (\beta'\beta/\alpha\beta) \times \gamma)^2} \times \sqrt{\delta^2_{\beta'\beta} + \left(\frac{\beta'\beta}{\alpha\beta}\right)^2 \times \delta^2_{\alpha\beta}}$$

The error in the intensity determination is mainly associated to the choice of two parameters that influence background removal. Here, we use the rollingBall function of the R package baseline, which has 2 arguments : $wm$ (width of local window for minimization/maximization) and $ws$ (width local window for smoothing). We estimate a maximum relative error on both $\alpha\beta$ and $\beta'\beta$ of ~7%, which results in a typical absolute error not greater than ~2% on $\Psi_\beta$ .

#### Numerical calculations

We solved differential equations using the R package deSolve and the R function lsoda. According to our kinetic model (see main text and Figure 3), the differential equations read :

$$
\begin{aligned}
\frac{d[\alpha\beta]}{dt} &= +k_\lambda \times [\alpha \cdot \beta \cdot L] \\
\frac{d[\alpha\alpha' \cdot E]}{dt} &= +\kappa[E] \times [\alpha\alpha'] - k_\epsilon \times [\alpha\alpha' \cdot E] \\
\frac{d[\beta'\beta \cdot E]}{dt} &= +\kappa[E] \times [\beta'\beta] - k_\epsilon \times [\beta'\beta \cdot E] \\
\frac{d[\alpha \cdot \alpha' \cdot L]}{dt} &= +\kappa_{\alpha\cdot\alpha'}[L] \times [\alpha \cdot \alpha'] - k_\lambda \times [\alpha \cdot \alpha' \cdot L] \\
\frac{d[\beta' \cdot \beta \cdot L]}{dt} &= +\kappa_{\beta'\cdot\beta}[L] \times [\beta' \cdot \beta] - k_\lambda \times [\beta' \cdot \beta \cdot L] \\
\frac{d[\alpha \cdot \beta \cdot L]}{dt} &= +\kappa_{\alpha\cdot\beta}[L] \times [\alpha \cdot \beta] - k_\lambda \times [\alpha \cdot \beta \cdot L] \\
\frac{d[\alpha\alpha']}{dt} &= -\kappa[E] \times [\alpha\alpha'] + k_\lambda \times [\alpha \cdot \alpha' \cdot L] \\
\frac{d[\beta'\beta]}{dt} &= -\kappa[E] \times [\beta'\beta] + k_\lambda \times [\beta' \cdot \beta \cdot L] \\
\frac{d[\alpha \cdot \alpha']}{dt} &= -\kappa_{\alpha\cdot\alpha'}[L] \times [\alpha \cdot \alpha'] + k_\epsilon \times [\alpha\alpha' \cdot E] + k_+ \times [\alpha] \times [\alpha'] - k_- \times [\alpha \cdot \alpha'] \\
\frac{d[\beta' \cdot \beta]}{dt} &= -\kappa_{\beta'\cdot\beta}[L] \times [\beta' \cdot \beta] + k_\epsilon \times [\beta'\beta \cdot E] + k_+ \times [\beta'] \times [\beta] - k_- \times [\beta' \cdot \beta] \\
\frac{d[\alpha \cdot \beta]}{dt} &= -\kappa_{\alpha\cdot\beta}[L] \times [\alpha \cdot \beta] + k_+ \times [\alpha] \times [\beta] - k_- \times [\alpha \cdot \beta] \\
\frac{d[\alpha]}{dt} &= -k_+ \times [\alpha] \times \big([\alpha'] + [\beta]\big) + k_- \times \big([\alpha \cdot \alpha'] + [\alpha \cdot \beta]\big) \\
\frac{d[\beta]}{dt} &= -k_+ \times [\beta] \times \big([\beta'] + [\alpha]\big) + k_- \times \big([\beta' \cdot \beta] + [\alpha \cdot \beta]\big) \\
\frac{d[\alpha']}{dt} &= -k_+ \times [\alpha'] \times \big([\alpha] + [\beta']\big) + k_- \times \big([\alpha \cdot \alpha'] + [\beta' \cdot \alpha']\big) \\
\frac{d[\beta']}{dt} &= -k_+ \times [\beta'] \times \big([\beta] + [\alpha']\big) + k_- \times \big([\beta' \cdot \beta] + [\beta' \cdot \alpha']\big) \\
\frac{d[\beta' \cdot \alpha']}{dt} &= -\kappa_{\beta'\cdot\alpha'}[L] \times [\beta' \cdot \alpha'] + k_\epsilon \times [\beta'\alpha' \cdot E] + k_+ \times [\alpha'] \times [\beta'] - k_- \times [\beta' \cdot \alpha'] \\
\frac{d[\beta' \cdot \alpha' \cdot L]}{dt} &= +\kappa_{\beta'\cdot\alpha'}[L] \times [\beta' \cdot \alpha'] - k_\lambda \times [\beta' \cdot \alpha' \cdot L] \\
\frac{d[\beta'\alpha']}{dt} &= -\kappa[E] \times [\beta'\alpha'] + k_\lambda \times [\beta' \cdot \alpha' \cdot L] \\
\frac{d[\beta'\alpha' \cdot E]}{dt} &= +\kappa[E] \times [\beta'\alpha'] - k_\epsilon \times [\beta'\alpha' \cdot E]
\end{aligned}
$$

At each step, the value of all compounds is calculated and allows to determine both the percentage of product formed $\Psi_\beta$ and the efficiency $\xi$.

$$
\Psi_\beta = 100 \times \frac{[\alpha\beta]}{[\beta'\beta] + [\alpha\beta] + [\beta'\beta \cdot E] + [\beta' \cdot \beta \cdot L] + [\alpha \cdot \beta \cdot L] + [\beta' \cdot \beta] + [\alpha \cdot \beta] + [\beta]}
$$

$$
\xi = 100 \times \frac{[\alpha\beta]}{[\alpha\alpha'] + [\beta'\beta] + [\alpha\beta] + [\alpha\alpha' \cdot E] + [\beta'\beta \cdot E] + [\alpha \cdot \alpha' \cdot L] + [\beta' \cdot \beta \cdot L] + [\alpha \cdot \beta \cdot L] + [\alpha \cdot \alpha'] + [\beta' \cdot \beta] + [\alpha \cdot \beta] + [\alpha] + [\beta]}
$$

### SUPPLEMENTARY TABLES

#### Overhang stabilities and stacking energies

| `CCGC` | `CACC` | `AGGA` | `TTCT` | `AGAA` |
|---|---|---|---|---|
| -6.25 | -4.73 | -4.42 | -3.58 | -3.58 |

| | `CCGC` | `CCGC` | `CACC` | `AGGA` | `TTCT` | `AGAA` |
|---|---|---|---|---|---|---|
| $\alpha\beta$ | -9.86 (`GCCGCC`) | -8.74 (`TCCGCT`) | -7.22 (`TCACCT`) | -5.95 (`TAGGAT`) | -5.80 (`TTTCTT`) | -5.11 (`TAGAAT`) |
| $\alpha\alpha'$ | -9.86 (`GCCGCC`) | -9.12 (`TCCGCC`) | -7.60 (`TCACCC`) | -6.42 (`TAGGAC`) | -6.12 (`TTTCTC`) | -5.58 (`TAGAAC`) |
| $\beta'\beta$ | -9.86 (`GCCGCC`) | -9.48 (`GCCGCT`) | -7.96 (`GCACCT`) | - 7.19 (`GAGGAT`) | -6.50 (`GTTCTT`) | -6.35 (`GAGAAT`) |
| $\alpha'\beta'$ | -9.86 (`GCCGCC`) | -9.86 (`GCCGCC`) | -8.34 (`GCACCC`) | - 7.66 (`GAGGAC`) | -6.82 (`GTTCTC`) | -6.82 (`GAGAAC`) |

Supplementary Table S1: Top. Overhang stabilities in kcal/mol obtained from [8] (Table 1). Bottom. Total free energies in kcal/mol for all possible products. For a sequence of six nucleotides and a 4 bp overhang, total free energies are the sum of the overhang stability (determined by nucleotides in positions 2,3,4 and 5) and the stacking energies (determined by nucleotides in positions 1 and 2 and in positions 5 and 6 using values of [9]). Overhang stabilities are also presented in Figure 2 of the main text. For instance, the overhang `TCCGCT` has a stability of -6.25 kcal/mol. The value of -2.49kcal/mol appearing in Figure 2 for `TCCGCT` represents the stacking energy, as obtained from Supplementary Table S2. Note that the sum of -6.25 and -2.49 gives -8.74 kcal/mol, which is the value displayed in Figure 2 for the fragment $\alpha\beta$.

| | | N2-5 | | | |
|---|---|---|---|---|---|
| | | `CCGC` or `CACC` | | | |
| | | N6 | | | |
| | | `T` | `A` | `G` | `C` |
| N1 | `T` | -2.49 | -1.98 | -2.34 | -2.87 |
| | `A` | -2.87 | -2.36 | - 2.72 | -3.25 |
| | `G` | -3.23 | -2.72 | -3.08 | -3.61 |
| | `C` | -2.50 | -1.99 | -2.35 | -2.88 |

| | | N2-5 | | | |
|---|---|---|---|---|---|
| | | `AGGA` or `AGAA` | | | |
| | | N6 | | | |
| | | `T` | `A` | `G` | `C` |
| N1 | `T` | -1.53 | -1.30 | -1.25 | -2.00 |
| | `A` | -2.45 | -2.22 | -2.17 | -2.92 |
| | `G` | -2.77 | -2.54 | -2.49 | -3.24 |
| | `C` | -1.89 | -1.66 | -1.61 | -2.36 |

| | | N2-5 | | | |
|---|---|---|---|---|---|
| | | `TTCT` | | | |
| | | N6 | | | |
| | | `T` | `A` | `G` | `C` |
| N1 | `T` | -2.22 | -1.30 | -1.66 | - 2.54 |
| | `A` | -2.45 | - 1.53 | -1.89 | -2.77 |
| | `G` | -2.92 | -2.00 | -2.36 | - 3.24 |
| | `C` | -2.17 | -1.25 | -1.61 | -2.49 |

Supplementary Table S2: Stacking energies (in kcal/mol) calculated using values of [9] for a sequence of six nucleotides. Each table corresponds to specific 4 bp overhangs. Left: `CCGC` or `CACC` in positions 2, 3, 4 and 5. Middle: `AGGA` or `AGAA` in positions 2, 3, 4 and 5. Right: `TTCT` in positions 2, 3, 4 and 5. N1 (N6) indicates the nucleotide in position 1 (6). Choosing the `ACCGCA` sequence, one needs to obtain stacking energies for `AC` (nucleotides in positions 1 and 2) and `CA` (nucleotides in positions 5 and 6). Those are given by [9] and yield a total stacking energy of -(1.81+0.55) kcal/mol or -2.36 kcal/mol (highlighted in bold).

#### Percentage of product formed (Figure 2 of the main text)

| | | GCCGCC | TCCGCT | TCACCT | TAGGAT | TTTCTT | TAGAAT |
|---|---|---|---|---|---|---|---|
| Overhang stability (kcal/mol) | | -6.25 | -6.25 | -4.73 | -4.42 | -3.58 | -3.58 |
| Total free energy (kcal/mol) | | -9.86 | -8.74 | -7.22 | -5.95 | -5.80 | -5.11 |
| 10 minutes | | | | | | | |
| | 37 | 66.0 | 65.0 | 72.6 | 50.1 | 32.8 | 22.4 |
| | 37/16 | 52.7 | 50.6 | 62.0 | 55.8 | 35.8 | 37.4 |
| 30 minutes | | | | | | | |
| | 37 | 84.2 | 81.3 | 81.5 | 76.2 | 66.6 | 53.4 |
| | 37/16 | 81.7 | 73.0 | 81.1 | 79.7 | 69.6 | 67.7 |
| 70 minutes | | | | | | | |
| | 37 | 86.7 | 83.1 | 85.9 | 81.7 | 78.8 | 75.3 |
| | 37/16 | 85.8 | 82.4 | 82.6 | 84.5 | 80.9 | 77.6 |
| 140 minutes | | | | | | | |
| | 37 | 87.9 | 85.3 | 88.0 | 82.9 | 87.7 | 77.1 |
| | 37/16 | 87.2 | 83.7 | 86.3 | 85.5 | 86.9 | 80.6 |
| 240 minutes | | | | | | | |
| | 37 | 89.4 | 87.2 | 88.6 | 83.0 | 83.7 | 80.5 |
| | 37/16 | 88.8 | 85.8 | 87.9 | 86.7 | 85.2 | 79.6 |

Supplementary Table S3: The percentage of product formed $\Psi_\beta$ averaged over three replicates for the different overhangs used in this study. For GCCGCC and TAGAAT at 10 minutes, values were averaged over five replicates. $\Psi_\beta$ were measured for both static experiment (37°C) and a cycling experiment (37°C for 5 minutes and 16°C for 5 minutes). Also shown are overhang stabilities and total free energies (kcal/mol) from Supplementary Table S1.

| | 10' | 30' | 70' | 140' | 240' |
|---|---|---|---|---|---|
| Overhang stability | | | | | |
| 37 | -0.80 | -0.81 | -0.78 | -0.45 | -0.80 |
| 37/16 | -0.55 | -0.54 | -0.68 | -0.27 | -0.62 |
| Total free energy | | | | | |
| 37 | -0.79 | -0.81 | -0.82 | -0.59 | -0.89 |
| 37/16 | -0.48 | -0.56 | -0.69 | -0.42 | -0.68 |

Supplementary Table S4: The Pearson correlation coefficient calculated between averaged $\Psi_\beta$ values (Supplementary Table S3) and the overhang stabilities (or the total free energies, as obtained from Supplementary Table S1). See also Supplementary Table S5.

| Pearson coefficient Threshold | 0.9 | 0.8 | 0.7 | 0.6 | 0.5 | 0.4 | 0.3 |
|---|---|---|---|---|---|---|---|
| Probability | 0.015 | 0.056 | 0.122 | 0.208 | 0.312 | 0.432 | 0.563 |

Supplementary Table S5: The probability that six independent and identically distributed (i.i.d) variables give a Pearson correlation coefficient larger or equal than a given threshold. For instance, there is a 1.5% chance that six i.i.d variables give a Pearson coefficient larger than 0.9. Here we just experimentally calculated those values by generating a large number of events (1,000,000).

#### Parameters used for the kinetic model

| | | 37°C | 16°C |
|---|---|---|---|
| $k_\epsilon$ | (1/s) | 10 | 0 |
| $k_\lambda$ | (1/s) | 10 | 4 |
| $\kappa[L]$ | (1/s) | 2.5 | 2.5 |
| $\kappa[E]$ | (1/s) | 0.012 | 0.012 |

Supplementary Table S6: $k_\epsilon$ , $k_\lambda$, $\kappa$ are assumed to be temperature (37°C, 16°C), substrate ($\alpha\alpha'$, $\alpha \cdot \alpha'$, $\beta'\beta$, $\beta' \cdot \beta$, $\beta'\alpha'$, $\beta' \cdot \alpha'$, $\alpha \cdot \beta$) and overhang (`GCCGCC`, `TAGAAT`) independent. Here $k_\epsilon$ and $k_\lambda$ were set to 10 $s^{-1}$, in agreement with [10]. We assumed a 100 % and a 60% decrease in activity when changing the temperature from 37°C to 16°C [11]. $\kappa[L]$ and $\kappa[E]$ were chosen to fit experimental values (cycling) at 10 minutes.

| | $k_-$ (1/s) | | $k_+$ (1/(M·s)) | | $\Delta G^0$ (kcal/mol) | |
|---|---|---|---|---|---|---|
| | 37°C | 16°C | 37°C | 16°C | 37°C | 16°C |
| `GCCGCC` | 1 | 1 | $8.5 \times 10^6$ | $3.7 \times 10^8$ | -9.83 | -11.34 |
| `TAGAAT` | 6 | 6 | $2.5 \times 10^4$ | $5.1 \times 10^5$ | -5.15 | -6.52 |

Supplementary Table S7: $k_-$ and $k_+$ are assumed to be both temperature and overhang dependent and substrate independent. Here we set identical values for $k_-$ at 37°C and at 16°C. We then calculated dissociation constants at 37°C and 16°C using values form [8] (Table 2) and [9] assuming that the temperature dependence of staking energies was negligible, in agreement with [12]. $k_-$ are were chosen to fit experimental values (cycling) at 10 minutes with the additional constraint that the derived $k_+$ values could not exceed the diffusion limit. Moreover, $k_-$values were required to be lower for a strong overhang (`GCCGCC`), which seems intuitive, even though thermodynamics does not necessarily impose this constraint.

| | $k_-$ (1/s) | | $k_+$ (1/(M·s)) | | $\Delta G^0$ (kcal/mol) | |
|---|---|---|---|---|---|---|
| | 37°C | 16°C | 37°C | 16°C | 37°C | 16°C |
| `GCCGCC` | 0.9 | 0.05 | $8.0 \times 10^6$ | $2.0 \times 10^7$ | -9.83 | -11.34 |
| `TAGAAT` | 18.9 | 2.4 | $8.0 \times 10^4$ | $2.0 \times 10^5$ | -5.15 | -6.52 |

.

Supplementary Table S8: In contrast to Supplementary Table S7, we did not set identical values for $k_-$ at 37°C and at 16°C. $k_+$ are were chosen to fit experimental values (cycling) at 10 minutes and $k_-$ were calculated from the dissociation constants at 37°C and at 16°C using values form [8] using the same approximations as in Supplementary Table S7. Moreover, $k_-$values were required to be lower for a strong overhang (`GCCGCC`), which seems intuitive, even though thermodynamics does not necessarily impose this constraint.

#### Primers

| | $\alpha\alpha'$ | $\beta'\beta$ |
|---|---|---|
| GCCGCC | OL082-OL106 | OL090-OL103 |
| TCCGCT | OL082-OL098 | OL090-OL093 |
| TCACCT | OL082-OL099 | OL090-OL094 |
| TAGGAT | OL082-OL097 | OL090-OL092 |
| TTTCTT | OL082-OL105 | OL090-OL102 |
| TAGAAT | OL082-OL111 | OL090-OL110 |
| TCCGCT* | OL082-OL131 | OL090-OL132 |
| TAGAAT* | OL082-OL135 | OL090-OL136 |

Supplementary Table S9: Primers used for generating different overhangs in GGA assemblies with isolated substrates. Exact sequences are given in the Supplementary Section DNA sequences.

| Optimal Set | | Non-optimal Set | | Fragment |
|---|---|---|---|---|
| Pair | Length | Pair | Length | |
| OL1388-OL1457 | 325 | OL1388-OL1475 | 348 | 1 |
| OL1458-OL1459 | 357 | OL1476-OL1477 | 345 | 2 |
| OL1460-OL1461 | 337 | OL1478-OL1479 | 335 | 3 |
| OL1462-OL1463 | 359 | OL1480-OL1481 | 325 | 4 |
| OL1464-OL1465 | 325 | OL1482-OL1483 | 349 | 5 |
| OL1466-OL1467 | 341 | OL1484-OL1485 | 334 | 6 |
| OL1468-OL1469 | 334 | OL1486-OL1487 | 352 | 7 |
| OL1470-OL1471 | 346 | OL1488-OL1489 | 328 | 8 |
| OL1472-OL1473 | 347 | OL1490-OL1491 | 352 | 9 |
| OL1492-OL1387 | 2275 | OL1492-OL1387 | 2275 | 10 |

Supplementary Table S10: Primer pairs used to generate either short PCR fragments (approximately 300 bp in length, fragments numbered 1 to 9) or a long fragment (approximately 2,200 bp in length, fragment 10) for the optimal and non-optimal sets. PCR have been performed on the pNJ2075 plasmid and allow to reconstitute a lac operon cassette (Supplementary Figure S14). Exact sequences are given in the Supplementary Section DNA sequences.

#### Estimating the number of transformants

| Total | Fidelity | Condition | Temperature | Time |
|---|---|---|---|---|
| 4 | 0 | Non-Optimal | 37 | 20 |
| 32 | 0.57 | Optimal | 37 | 20 |
| 45 | 0.7 | Non-Optimal | 37/16 | 20 |
| 351 | 0.94 | Optimal | 37/16 | 20 |
| 22 | 0 | Non-Optimal | 37 | 30 |
| 432 | 0.89 | Optimal | 37 | 30 |
| 522 | 0.9 | Non-Optimal | 37/16 | 30 |
| 1048 | 0.91 | Optimal | 37/16 | 30 |
| 36 | 0.12 | Non-Optimal | 37 | 60 |
| 2110 | 0.96 | Optimal | 37 | 60 |
| 2066 | 0.93 | Non-Optimal | 37/16 | 60 |
| 2588 | 0.92 | Optimal | 37/16 | 60 |

Supplementary Table S11: Results for the 10 fragments GGA assembly, duplicate 1. Shown here are the total number of colonies, including both white and blue colonies (Total), as well as the ratio of the number of blue colonies to the total number of colonies (Fidelity) for different conditions (either by cycling the temperature or using a static temperature) obtained at different time points (Time, in minutes). The number of colonies was obtained by summing the number of transformants on two different plates (each containing 100 $\mu$L of the outgrowth) and normalizing by the ratio of the total outgrowth volume (900 $\mu$L) to the total plating volume (200 $\mu$L). For details, see the protocol page 17 of this document.

| Total | Fidelity | Condition | Temperature | Time |
|---|---|---|---|---|
| 10 | 0 | Non-Optimal | 37 | 20 |
| 252 | 0.89 | Optimal | 37 | 20 |
| 133 | 0.68 | Non-Optimal | 37/16 | 20 |
| 840 | 0.91 | Optimal | 37/16 | 20 |
| 14 | 0 | Non-Optimal | 37 | 30 |
| 658 | 0.93 | Optimal | 37 | 30 |
| 973 | 0.92 | Non-Optimal | 37/16 | 30 |
| 1712 | 0.94 | Optimal | 37/16 | 30 |
| 63 | 0.5 | Non-Optimal | 37 | 60 |
| 2443 | 0.91 | Optimal | 37 | 60 |
| 3020 | 0.93 | Non-Optimal | 37/16 | 60 |
| 4634 | 0.92 | Optimal | 37/16 | 60 |

Supplementary Table S12: Results for the 10 fragments GGA assembly, duplicate 2. Same as Supplementary Table S11 using a total outgrowth volume of 700 $\mu$L. While this duplicate assembly was performed under the exact same conditions as the previous one, the transformation was carried out using slightly different experimental conditions (see the protocol page 17 of this document). This variation could explain the observed differences in the total number of colonies.

### PROTOCOLS

#### • 2 FRAGMENTS GGA ASSEMBLY

##### – PCR reactions

###### ∗ PCR Reactions for $\alpha\alpha'$ fragments

**Note** $\alpha\alpha'$ are 565 bp in length.

**Note** $\alpha\alpha'$ share the same primer (OL082).

**Master Mix for up to 9 reactions in parallel**

| Stock | Volume ($\mu$L) | Final |
|---|---|---|
| 10X Standard Taq Reaction Buffer | 46 | 1.15X |
| pEXK4pSAM001 @ 25 ng/$\mu$L | 3 | 0.19 ng/$\mu$L |
| 10 mM dNTPs | 9 | 225 $\mu$M |
| Taq DNA Polymerase (NEB M0495) - Hot Start | 2.4 | 0.03 U/$\mu$L |
| OL082 @ 10 $\mu$M | 9 | 225 nM |
| H20 | 330.6 | |
| Total | 400 | |

**Reaction Mix**

| Stock | Volume ($\mu$L) | Final |
|---|---|---|
| Master Mix | 44 | 1X Taq Buffer, 200 $\mu$M dNTP, 200 nM OL082, 0.026 U/$\mu$L Taq |
| Primer @ 10 $\mu$M | 1 | 200 nM |
| H20 | 5 | |
| Total | 50 | |

**Reaction**

Split in 4 or 5 tubes (12.5 or 10 $\mu$L).

- 1X : 30 s @ 95°C.
- 25 X : 30 s @ 95°C, 30 s @53°C, 60 s @68°C.
- 1X : 300 s @ 68°C.
- Hold @ 14°C.

###### ∗ PCR Reactions for $\beta'\beta$ fragments

**Note** $\beta'\beta$ are 1678 bp in length.

**Note** $\beta'\beta$ share the same primer (OL090).

**Master Mix for up to 8 reactions in parallel**

| Stock | Volume ($\mu$L) | Final |
|---|---|---|
| 5X Q5 Reaction Buffer | 89 | 1.11X |
| 10 mM dNTPs | 8.9 | 222 $\mu$M |
| OL090 @ 10 $\mu$M | 22.2 | 555 nM |
| pEXK4pSAM001 @ 25 ng/$\mu$L | 3 | 0.19 ng/$\mu$L |
| Q5 High-Fidelity DNA Polymerase (NEB M0493) - Hot Start | 4.5 | 0.0225 U/$\mu$L |
| H20 | 272.4 | |
| Total | 400 | |

**Reaction Mix**

| Stock | Volume ($\mu$L) | Final |
|---|---|---|
| Master Mix | 45 | 1X Q5 Buffer, 200 $\mu$M dNTP, 500 nM OL090, 0.02 U/$\mu$L Q5 |
| Primer @ 10 $\mu$M | 2.5 | 500 nM |
| H20 | 2.5 | |
| Total | 50 | |

**Reaction**

Split in 4 or 5 tubes (12.5 or 10 $\mu$L).

- 1X : 30 s @ 98°C.
- 25 X : 10 s @ 98°C, 30 s @54°C, 45 s @54°C.
- 1X : 120 s @ 72°C.
- Hold @ 14°C.

#### – Assemblies at a fixed Ligase concentration (12U/$\mu$L)

**Prepare Master Mix for 2 reactions (e.g. 2 sequences in parallel)**

| Stock | Volume ($\mu$L) |
|---|---|
| Ligase (M0202 NEB) | 1.8 |
| BsaI-HFv2 (R3733 NEB) | 1.8 |
| ATP (10 mM) | 6 |
| DTT (100 mM) | 6 |
| rCutsmart 10 X | 6 |
| H20 | 7.2 |
| Total | 28.8 |

**Reaction Mix (below is for 1 sequence only)**

| Stock | Volume ($\mu$L) | Final |
|---|---|---|
| $\beta'\beta$ @ 37 ng/$\mu$L (1678 bp) | 5.2 | ~7 nM |
| $\alpha\alpha'$ @ 25 ng/$\mu$L (565 bp) | 7.8 | ~21 nM |
| Master Mix | 12 | 1 mM ATP, 10 mM DTT, rCutsmart 1X, Enzymes : 0.75 $\mu$L |
| Total | 25 | |

**Reaction (below is for 1 sequence only)**

- ∗ Take 2$\mu$L of Reaction Mix, add 5 $\mu$L SDS (Stock 0.3%) and heat at 65°C for at least 10 minutes.
- ∗ Split remaining 23 $\mu$L in 2 tubes.
- ∗ 1 tube in dry bath (37°C, static experiment), 1 tube in thermocycler (37°C/16°C, 5 minutes at each temperature, cycling experiment).
- ∗ At 10, 30,70, 140 and 240 minutes, take 2 $\mu$L, add 5 $\mu$L SDS (Stock 0.3%) and heat at 65°C for at least 10 minutes.

**Agarose Gel**

* Ladder (SM1173, Fisher Scientific), 3 $\mu$L
* 1/2 S0 (lane to determine stoichiometry)
* 10 Cycling
* 10 Static
* 30 Cycling
* 30 Static
* 70 Cycling
* 70 Static
* 140 Cycling
* 140 Static
* 240 Cycling
* 240 Static
* 1/2 S0 (lane to determine stoichiometry)
* Ladder

Staining

* 1.2% in Tris Acetate EDTA (TAE).
* Staining in EtBr (5$\mu$L in 100 mL TAE) for 15 minutes.
* Destain twice in nanopure water for 10 minutes.

#### – Assemblies at varying Ligase concentrations

**Prepare Master Mix - MASTER MIX-ENZ**

| Stock | Volume ($\mu$L) |
|---|---|
| BsaI-HFv2 (R3733 NEB) | 6 |
| ATP (10 mM) | 18 |
| DTT (100 mM) | 18 |
| rCutsmart 10 X | 18 |
| H20 | 6 |
| Total | 66 |

**Prepare Master Mix - MASTER MIX-DNA-1 (strong overhang, `GCCGCC`)**

| Stock | Volume ($\mu$L) |
|---|---|
| $\beta'\beta$ @ 37 ng/$\mu$L (1678 bp) | 18 |
| $\alpha\alpha'$ @ 25 ng/$\mu$L (565 bp) | 27 |
| Total | 45 |

**Prepare Master Mix - MASTER MIX-DNA-2 (weak overhang, `TAGAAT`)**

| Stock | Volume ($\mu$L) |
|---|---|
| $\beta'\beta$ @ 37 ng/$\mu$L (1678 bp) | 18 |
| $\alpha\alpha'$ @ 25 ng/$\mu$L (565 bp) | 27 |
| Total | 45 |

**Reaction 1 - strong overhang - Ligase @ 48 U/$\mu$L**

| Stock | Volume ($\mu$L) |
|---|---|
| MASTER MIX-ENZ | 9 |
| MASTER MIX-DNA-1 | 13 |
| Ligase (M0202 NEB) | 3 |
| Total | 25 |

- ∗ Take 4 μL in 10 μL SDS (Stock 0.3%) and Heat at 65°C for 20': $\text{Sample}_0$.
- ∗ Split remaining 2 x 10.5 μL.
- ∗ 10.5 μL will cycle (37°C for 5 minutes and 16°C for 5 minutes) for a total of 10 minutes.
- ∗ 10.5 μL will not cycle.
- ∗ At 10 minutes: take 2 μL in 5 uL SDS 0.3% + Heat Shock at 65°C for at least 10 minutes. Do that for cycling and no cycling.

[Final : ATP @ 1mM, CutSmart 1X, BsaI-HFv2 : 0.65 U/μL, DTT @ 10 mM]
[Main fragment (1678 bp) @ 7 nM, Extension (565 bp)@ 21 nM]
[Ligase @ 48 U/μL]

**Reaction 2 - weak overhang - Ligase @ 48 U/μL**

| Stock | Volume (μL) |
|---|---|
| MASTER MIX-ENZ | 9 |
| MASTER MIX-DNA-2 | 13 |
| Ligase (M0202 NEB) | 3 |
| Total | 25 |

**Reaction 3 - strong overhang - Ligase @ 12 U/μL**

| Stock | Volume (μL) |
|---|---|
| MASTER MIX-ENZ | 9 |
| MASTER MIX-DNA-1 | 13 |
| Ligase (M0202 NEB) | 0.75 |
| H20 | 2.25 |
| Total | 25 |

**Reaction 4 - weak overhang - Ligase @ 12 U/μL**

| Stock | Volume (μL) |
|---|---|
| MASTER MIX-ENZ | 9 |
| MASTER MIX-DNA-2 | 13 |
| Ligase (M0202 NEB) | 0.75 |
| H20 | 2.25 |
| Total | 25 |

**Ligase dilution** Take 3 μL of Ligase and mix with 45 μL Cutsmart 1X

**Reaction 5 -strong overhang - Ligase @ 3 U/μL**

| Stock | Volume (μL) |
|---|---|
| MASTER MIX-ENZ | 9 |
| MASTER MIX-DNA-1 | 13 |
| Ligase (M0202 NEB, diluted) | 3 |
| Total | 25 |

**Reaction 6 - weak overhang - Ligase @ 3 U/μL**

| Stock | Volume (μL) |
|---|---|
| MASTER MIX-ENZ | 9 |
| MASTER MIX-DNA-2 | 13 |
| Ligase (M0202 NEB, diluted) | 3 |
| Total | 25 |

### • 10 FRAGMENTS GGA ASSEMBLY

#### – PCR reactions

**Master Mix**

| Stock | Volume (μL) | Final |
|---|---|---|
| 5X Q5 Reaction Buffer | 120 | 1.11X |
| 10 mM dNTPs | 12 | 222 μM |
| pNJ2075 @ 10 ng/μL | 5 | 50 ng |
| Q5 High-Fidelity DNA Polymerase (NEB M0493) - Hot Start | 6 | 12 U |
| H20 | 397 | |
| Total | 540 | |

**Reaction Mix**

| Stock | Volume (μL) | Final |
|---|---|---|
| Master Mix | 22.5 | 1X Q5 Buffer, 200 μM dNTP, 0.02 U/μL Q5 |
| Primer1 @ 5 μM | 1.25 | 250 nM |
| Primer2 @ 5 μM | 1.25 | 250 nM |
| Total | 25 | |

**Reaction**

* 1X : 30 s @ 98°C.
* 25 X : 10 s @ 98°C, 20 s @72°C [2-steps PCR]
* Hold @ 14°C.

**Note**

* For the longer fragment involving OL1474 and OL1487, the elongation was set to 40 seconds.
* For the fragment involving OL1462 and OL1463, the final concentration of primers was set to 125 nM to reduce primer-dimer formation.

#### – Assembly

We used a protocol similar to that proposed by NEB using the NEBridge Golden Gate Assembly Kit (BsaI-HFv2 (E1601).

* Prepare GGA assembly

**Step 0**

Resuspend all oligos to 60ng/μL

| A | B | AB |
|---|---|---|
| 1388-1475 | 1388-1457 | 1492-1387 |
| 1476-1477 | 1458-1459 | |
| 1478-1479 | 1460-1461 | |
| 1480-1481 | 1462-1463 | |
| 1482-1483 | 1464-1465 | |
| 1484-1485 | 1466-1467 | |
| 1486-1487 | 1468-1469 | |
| 1488-1489 | 1470-1471 | |
| 1490-1491 | 1472-1473 | |

**A** is the non-optimal set, **B** is the optimal set. The prefix `OL` has been omitted.

Take 8.5 $\mu$L of 1492-1387 (AB) and add 1.5 $\mu$L $H_2O$ (total is 10 $\mu$L).

**Step 1**

Prepare 2 tubes (labelled A and B).
- A contains 0.6 $\mu$L of each A short PCR fragments (75-78 $\rightarrow$ 90-91) and 4.6 $\mu$L of 92-87 (step 0).
- B contains 0.6 $\mu$L of each B short PCR fragments (57-88 $\rightarrow$ 72-73) and 4.6 $\mu$L of 92-87 (step 0).
[Short fragments are 340 bp long on average, Long fragment (AB) is 2275 bp]
[In A and B, the final concentration of all PCR fragments is about 16 nM]

**Step 2**

Using E1601S (Link) kit from NEB that contains:
- NEBridge Golden Gate Enzyme Mix (BsaI-HFv2) - M2616AVIAL
- T4 DNA Ligase Reaction Buffer - B0202SVIAL
- pGGAselect DNA - N0309AAVIAL

Prepare a MIX (Total is: 35 $\mu$L)

| | Volume ($\mu$L) |
|---|---|
| Enzyme Mix (M2616AVIAL) | 2.5 |
| BufferT4 (B0202SVIAL, 10X) | 5 |
| pGGAselect | 2.5 |
| $H_2O$ | 25 |

Then

| | Volume ($\mu$L) |
|---|---|
| A | 6 |
| MIX | 14 |

[In A+MIX, the final concentration of all PCR fragments is about 4.8 nM. Plasmid is about 2.6 nM]
[In A+MIX, Buffer is 1X, and there is 1 $\mu$L of Enzyme Mix]

| | Volume ($\mu$L) |
|---|---|
| B | 6 |
| MIX | 14 |

[In B+MIX, the final concentration of all PCR fragments is about 4.8 nM. Plasmid is about 2.6 nM]
[In B+MIX, Buffer is 1X, and there is 1 $\mu$L of Enzyme Mix]

**Step 3**

Split A (+MIX) and B (+MIX) into 2 tubes of each 10 $\mu$L.

From tube A (+MIX), you get 2 new tubes → A3716 and A37
From tube B (+MIX), you get 2 new tubes → B3716 and B37
[For each tube A and B of 10 $\mu$L, the final concentration of all PCR fragments is about 4.8 nM; Plasmid is about 2.6nM; Buffer is 1X and there is 0.5 $\mu$L of Enzyme Mix]

∗ Reaction

A3716: (37°C, 1 min → 16°C, 1 min) x 30
A37: (37°C for 60 min)
B3716: (37°C, 1 min → 16°C, 1 min) x 30
B37: (37°C for 60 min)

At 20 min:
- Take 2 $\mu$L of A3716 → 60°C for 5 min: Id# A3716_20
- Take 2 $\mu$L of A37 → 60°C for 5 min: Id# A37_20
- Take 2 $\mu$L of B3716 → 60°C for 5 min: Id# B3716_20
- Take 2 $\mu$L of B37 → 60°C for 5 min: Id# B37_20

At 30 min:
- Take 2 $\mu$L of A3716 → 60°C for 5 min: Id# A3716_30
- Take 2 $\mu$L of A37 → 60°C for 5 min: Id# A37_30
- Take 2 $\mu$L of B3716 → 60°C for 5 min: Id# B3716_30
- Take 2 $\mu$L of B37 → 60°C for 5 min: Id# B37_30

At 60 min:
- Take 2 $\mu$L of A3716 → 60°C for 5 min: Id# A3716_60
- Take 2 $\mu$L of A37 → 60°C for 5 min: Id# A37_60
- Take 2 $\mu$L of B3716 → 60°C for 5 min: Id# B3716_60
- Take 2 $\mu$L of B37 → 60°C for 5 min: Id# B37_60

#### – Transformation

Once the reactions were completed, the bacteria were transformed as follows. The assemblies were done in duplicate (starting from raw PCR fragments), and the two transformations conducted were slightly modified (see below).

| **Transformation** | **Assay 1** | **Assay 2** |
|---|---|---|
| T7 Express Competent *E. coli* (C2566I, NEB) | 50 µl | 30 µl |
| GGA reaction | 2 µl | 1.8 µl |
| Thermal shock time at 42°C | 20 sec | 10 sec |
| LB addition | 900 µl | 700 µl |
| Spreading | 2 x 100 µl | 2 x 100 µl |

### DNA SEQUENCES

#### Plasmid

**pNJ2075** CGGCCCCACGGGTGCGCATGATCGTGCTCCTGTCGTTGAGGACCCGGCTAGGCTGGCGGGGTTGCCTTACTGGTTAGCAGAATGAATCACCGATACGCGAGCGAAC
GTGAAGCGACTGCTGCTGCAAAACGTCTGCGACCTGAGCAACAACATGAATGGTCTTCGGTTTCCGTGTTTCGTAAAGTCTGGAAACGCGGAAGTCAGCGCCCTGCACCATTATGTTCC
GGATCTGCATCGCAGGATGCTGCTGGCTACCCTGTGGAACACCTACATCTGTATTAACGAAGCGCTGGCATTGACCCTGAGTGATTTTTCTCTGGTCCCGCCGCATCCATACCGCCAGT
TGTTTACCCTCACAACGTTCCAGTAACCGGGCATGTTCATCATCAGTAACCCGTATCGTGAGCATCCTCTCTCGTTTCATCGGTATCATTACCCCCATGAACAGAAATCCCCCTTACAC
GGAGGCATCAGTGACCAAACAGGAAAAAACCGCCCTTAACATGGCCCGCTTTATCAGAAGCCAGACATTAACGCTTCTGGAGAAACTCAACGAGCTGGACGCGGATGAACAGGCAGACA
TCTGTGAATCGCTTCACGACCACGCTGATGAGCTTTACCGCAGCTGCCTCGCGCGTTTCGGTGATGACGGTGAAAACCTCTGACACATGCAGCTCCCGGAGACGGTCACAGCTTGTCTG
TAAGCGGATGCCGGGAGCAGACAAGCCCGTCAGGGCGCGTCAGCGGGTGTTGGCGGGTGTCGGGGCGCAGCCATGACCCAGTCACGTAGCGATAGCGGAGTGTATACTGGCTTAACTAT
GCGGCATCAGAGCAGATTGTACTGAGAGTGCACCATTGCGGTGTGAAATACCGCACAGATGCGTAAGGAGAAAATACCGCATCAGGCGCTCTTCCGCTTCCTCGCTCACTGACTCGCTG
CGCTCGGTCGTTCGGCTGCGGCGAGCGGTATCAGCTCACTCAAAGGCGGTAATACGGTTATCCACAGAATCAGGGGATAACGCAGGAAAGAACATGTGAGCAAAAGGCCAGCAAAAGGC
CAGGAACCGTAAAAAGGCCGCGTTGCTGGCGTTTTTCCATAGGCTCCGCCCCCCTGACGAGCATCACAAAAATCGACGCTCAAGTCAGAGGTGGCGAAACCCGACAGGACTATAAAGAT
ACCAGGCGTTTCCCCCTGGAAGCTCCCTCGTGCGCTCTCCTGTTCCGACCCTGCCGCTTACCGGATACCTGTCCGCCTTTCTCCCTTCGGGAAGCGTGGCGCTTTCTCATAGCTCACGC
TGTAGGTATCTCAGTTCGGTGTAGGTCGTTCGCTCCAAGCTGGGCTGTGTGCACGAACCCCCCGTTCAGCCCGACCGCTGCGCCTTATCCGGTAACTATCGTCTTGAGTCCAACCCGGT
AAGACACGACTTATCGCCACTGGCAGCAGCCACTGGTAACAGGATTAGCAGAGCGAGGTATGTAGGCGGTGCTACAGAGTTCTTGAAGTGGTGGCCTAACTACGGCTACACTAGAAGGA
CAGTATTTGGTATCTGCGCTCTGCTGAAGCCAGTTACCTTCGGAAAAAGAGTTGGTAGCTCTTGATCCGGCAAACAAACCACCGCTGGTAGCGGTGGTTTTTTTGTTTGCAAGCAGCAG
ATTACGCGCAGAAAAAAAGGATCTCAAGAAGATCCTTTGATCTTTTCTACGGGGTCTGACGCTCAGTGGAACGAAAACTCACGTTAAGGGATTTTGGTCATGAACAATAAAACTGTCTG
CTTACATAAACAGTAATACAAGGGGTGTTATGAGCCATATTCAACGGGAAACGTCTTGCTCTAGGCCGCGATTAAATTCCAACATGGATGCTGATTTATATGGGTATAAATGGGCTCGC
GATAATGTCGGGCAATCAGGTGCGACAATCTATCGATTGTATGGGAAGCCCGATGCGCCAGAGTTGTTTCTGAAACATGGCAAAGGTAGCGTTGCCAATGATGTTACAGATGAGATGGT
CAGACTAAACTGGCTGACGGAATTTATGCCTCTTCCGACCATCAAGCATTTTATCCGTACTCCTGATGATGCATGGTTACTCACCACTGCGATCCCCGGGAAAACAGCATTCCAGGTAT
TAGAAGAATATCCTGATTCAGGTGAAAATATTGTTGATGCGCTGGCAGTGTTCCTGCGCCGGTTGCATTCGATTCCTGTTTGTAATTGTCCTTTTAACAGCGATCGCGTATTTCGTCTC
GCTCAGGCGCAATCACGAATGAATAACGGTTTGGTTGATGCGAGTGATTTTGATGACGAGCGTAATGGCTGGCCTGTTGAACAAGTCTGGAAAGAAATGCATAAACTTTTGCCATTCTC
ACCGGATTCAGTCGTCACTCATGGTGATTTCTCACTTGATAACCTTATTTTTGACGAGGGGAAATTAATAGGTTGTATTGATGTTGGACGAGTCGGAATCGCAGACCGATACCAGGATC
TTGCCATCCTATGGAACTGCCTCGGTGAGTTTTCTCCTTCATTACAGAAACGGCTTTTTCAAAAATATGGTATTGATAATCCTGATATGAATAAATTGCAGTTTCATTTGATGCTCGAT
GAGTTTTTCTAAGAATTAATTCATGAGCGGATACATATTTGAATGTATTTAGAAAAATAAACAAATAGGGGTTCCGCGCACATTTCCCCGAAAAGTGCCACCTAAATTGTAAGCGTTAA
TATTTTGTTAAAATTCGCGTTAAATTTTTGTTAAATCAGCTCATTTTTTAACCAATAGGCCGAAATCGGCAAAATCCCTTATAAATCAAAAGAATAGACCGAGATAGGGTTGAGTGTTG
TTCCAGTTTGGAACAAGAGTCCACTATTAAAGAACGTGGACTCCAACGTCAAAGGGCGAAAAACCGTCTATCAGGGCGATGGCCCACTACGTGAACCATCACCCTAATCAAGTTTTTTG
GGGTCGAGGTGCCGTAAAGCACTAAATCGGAACCCTAAAGGGAGCCCCCGATTTAGAGCTTGACGGGGAAAGCCGGCGAACGTGGCGAGAAAGGAAGGGAAGAAAGCGAAAGGAGCGGG
CGCTAGGGCGCTGGCAAGTGTAGCGGTCACGCTGCGCGTAACCACCACACCCGCCGCGCTTAATGCGCCGCTACAGGGCGCGTCCCATTCGCCAATCCGGATATAGTTCCTCCTTTCAG
CAAAAAACCCCTCAAGACCCGTTTAGAGGCCCCAAGGGGTTATGCTAGTTATTGCTCAGCGGTGGCAGCAGCCAACTCAGCTTCCTTTCGGGCTTTGTTAGCAGCCGGATCTCAGTGGT
GGTGGTGGTGGTGCTCGAGTGCGGCCGCTCATTATTTTTGACACCAGACCAACTGGTAATGGTAGCGACCGGCGCTCAGCTGGAATTCCGCCGATACTGACGGGCTCCAGGAGTCGTCG
CCACCAATCCCCATATGGAAACCGTCGATATTCAGCCATGTGCCTTCTTCCGCGTGCAGCAGATGGCGATGGCTGGTTTCCATCAGTTGCTGTTGACTGTAGCGGCTGATGTTGAACTG
GAAGTCGCCGCGCCACTGGTGTGGGCCATAATTCAATTCGCGCGTCCCGCAGCGCAGACCGTTTTCGCTCGGGAAGACGTACGGGGTATACATGTCTGACAATGGCAGATCCCAGCGGT
CAAAACAGGCGGCAGTAAGGCGGTCGGGATAGTTTTCTTGCGGCCCTAATCCGAGCCAGTTTACCCGCTCTGCTACCTGCGCCAGCTGGCAGTTCAGGCCAATCCGCGCCGGATGCGGT
GTATCGCTCGCCACTTCAACATCAACGGTAATCGCCATTTGACCACTACCATCAATCCGGTAGGTTTTCCGGCTGATAAATAAGGTTTTCCCCTGATGCTGCCACGCGTGAGCGGTCGT
AATCAGCACCGCATCAGCAAGTGTATCTGCCGTGCACTGCAACAACGCTGCTTCGGCCTGGTAATGGCCCGCCGCCTTCCAGCGTTCGACCCAGGCGTTAGGGTCAATGCGGGTCGCTT
CACTTACGCCAATGTCGTTATCCAGCGGTGCACGGGTGAACTGATCGCGCAGCGGCGTCAGCAGTTGTTTTTTATCGCCAATCCACATCTGTGAAAGAAAGCCTGACTGGCGGTTAAAT
TGCCAACGCTTATTACCCAGCTCGATGCAAAAATCCATTTCGCTGGTGGTCAGATGCGGGATGGCGTGGGACGCGGCGGGGAGTGTCACGCTGAGGTTTTCAGCCAGACGCCACTGCTG
CCAGGCGCTGATGTGTCCGGCTTCTGACCATGCGGTCGCGTTCGGTTGCACTACGCGTACTGTGAGCCAGAGTTGCCCGGCGCTCTCCGGCTGCGGTAGTTCAGGCAGTTCAATCAACT
GTTTACCTTGTGGAGCGACATCCAGAGGCACTTCACCGCTTGCCAGCGGCTTACCATCCAGCGCCACCATCCAGTGCAGGAGCTCGTTATCGCTATGACGGAACAGGTATTCGCTGGTC
ACTTCGATGGTTTGCCCGGATAAACGGAACTGGAAAAACTGCTGCTGGTGTTTTGCTTCCGTCAGCGCTGGATGCGGCGTGCGGTCGGCAAAGACCAGACCGTTCATACAGAACTGGCG
ATCGTTCGGCGTATCGCCAAAATCACCGCCGTAAGCCGACCACGGGTTGCCGTTTTCATCATATTTAATCAGCGACTGATCCACCCAGTCCCAGACGAAGCCGCCCTGTAAACGGGGAT
ACTGACGAAACGCCTGCCAGTATTTAGCGAAACCGCCAAGACTGTTACCCATCGCGTGGGCGTATTCGCAAAGGATCAGCGGGCGCGTCTCTCCAGGTAGCGAAAGCCATTTTTTGATG
GACCATTTCGGCACAGCCGGGAAGGGCTGGTCTTCATCCACGCGCGCGTACATCGGGCAAATAATATCGGTGGCCGTGGTGTCGGCTCCGCCGCCTTCATACTGCACCGGGCGGGAAGG
ATCGACAGATTTGATCCAGCGATACAGCGCGTCGTGATTAGCGCCGTGGCCTGATTCATTCCCCAGCGACCAGATGATCACACTCGGGTGATTACGATCGCGCTGCACCATTCGCGTTA
CGCGTTCGCTCATCGCCGGTAGCCAGCGCGGATCATCGGTCAGACGATTCATTGGCACCATGCCGTGGGTTTCAATATTGGCTTCATCCACCACATACAGGCCGTAGCGGTCGCACAGC
GTGTACCACAGCGGATGGTTCGGATAATGCGAACAGCGCACGGCGTTAAAGTTGTTCTGCTTCATCAGCAGGATATCCTGCACCATCGTCTGCTCATCCATGACCTGACCATGCAGAGG
ATGATGCTCGTGACGGTTAACGCCTCGAATCAGCAACGGCTTGCCGTTCAGCAGCAGCAGACCATTTTCAATCCGCACCTCGCGGAAACCGACATCGCAGGCTTCTGCTTCAATCAGCG
TGCCGTCGGCGGTGTGCAGTTCAACCACCGCACGATAGAGATTCGGGATTTCGGCGCTCCACAGTTTCGGGTTTTCGACGTTCAGACGTAGTGTGACGCGATCGGCATAACCACCACGC
TCATCGATAATTTCACCGCCGAAAGGCGCGGTGCCGCTGGCGACCTGCGTTTCACCCTGCCATAAAGAAACTGTTACCCGTAGGTAGTCACGCAACTCGCCGCACATCTGAACTTCAGC
CTCCAGTACAGCGCGGCTGAAATCATCATTAAAGCGAGTGGCAACATGGAAATCGCTGATTTGTGTAGTCGGTTTATGCAGCAACGAGACGTCACGGAAAATGCCGCTCATCCGCCACA
TATCCTGATCTTCCAGATAACTGCCGTCACTCCAGCGCAGCACCATCACCGCGAGGCGGTTTTCTCCGGCGCGTAAAAATGCGCTCAGGTCAAATTCAGACGGCAAACGACTGTCCTGG

CCGTAACCGACCCAGCGCCCGTTGCACCACAGATGAAACGCCGAGTTAACGCCATCAAAAATAATTCGCGTCTGGCCTTCCTGTAGCCAGCTTTCATCAACATTAAATGTGAGCGAGTA
ACAACCCGTCGGATTCTCCGTGGGAACAAACGGCGGATTGACCGTAATGGGATAGGTCACGTTGGTGTAGATGGGCGCATCGTAACCGTGCATCTGCCAGTTTGAGGGGACGACGACAG
TATCGGCCTCAGGAAGATCGCACTCCAGCCAGCTTTCCGGCACCGCTTCTGGTGCCGGAAACCAGGCAAAGCGCCATTCGCCATTCAGGCTGCGCAACTGTTGGGAAGGGCGATCGGTG
CGGGCCTCTTCGCTATTACGCCAGCTGGCGAAAGGGGGATGTGCTGCAAGGCGATTAAGTTGGGTAACGCCAGGGTTTTCCCAGTCACGACGTTGTAAAACGACGGCCAGTGAATCCGT
AATCATGGTCATATGCCCATGGTATATCTCCTTCTTAAAGTTAAACAAAATTATTTCTAGATCACCTTGTTATCCGCTCACAATTTCACACAACATACGAGCCGGAAGCATAAAGTGTA
AAGCCTGGGGTGCCTAATGAGTGAGCGCGGCAGATCTCGATCCTCTACGCCGGACGCATCGTGGCCGGCATCACCGGCGCCACAGGTGCGGTTGCTGGCGCCTATATCGCCGACATCAC
CGATGGGGAAGATCGGGCTCGCCACTTCGGGCTCATGAGCGCTTGTTTCGGCGTGGGTATGGTGGCAGGCCCCGTGGCCGGGGGACTGTTGGGCGCCATCTCCTTGCATGCACCATTCC
TTGCGGCGGCGGTGCTCAACGGCCTCAACCTACTACTGGGCTGCTTCCTAATGCAGGAGTCGCATAAGGGAGAGCGTCGAGATCCCGGACACCATCGAATGGCGCAAAACCTTTCGCGG
TATGGCATGATAGCGCCCGGAAGAGAGTCAATTCAGGGTGGTGAATGTGAAACCAGTAACGTTATACGATGTCGCAGAGTATGCCGGTGTCTCTTATCAGACCGTTTCCCGCGTGGTGA
ACCAGGCCAGCCACGTTTCTGCGAAAACGCGGGAAAAAGTGGAAGCGGCGATGGCGGAGCTGAATTACATTCCCAACCGCGTGGCACAACAACTGGCGGGCAAACAGTCGTTGCTGATT
GGCGTTGCCACCTCCAGTCTGGCCCTGCACGCGCCGTCGCAAATTGTCGCGGCGATTAAATCTCGCGCCGATCAACTGGGTGCCAGCGTGGTGGTGTCGATGGTAGAACGAAGCGGCGT
CGAAGCCTGTAAAGCGGCGGTGCACAATCTTCTCGCGCAACGCGTCAGTGGGCTGATCATTAACTATCCGCTGGATGACCAGGATGCCATTGCTGTGGAAGCTGCCTGCACTAATGTTC
CGGCGTTATTTCTTGATGTCTCTGACCAGACACCCATCAACAGTATTATTTTCTCCCATGAAGACGGTACGCGACTGGGCGTGGAGCATCTGGTCGCATTGGGTCACCAGCAAATCGCG
CTGTTAGCGGGCCCATTAAGTTCTGTCTCGGCGCGTCTGCGTCTGGCTGGCTGGCATAAATATCTCACTCGCAATCAAATTCAGCCGATAGCGGAACGGGAAGGCGACTGGAGTGCCAT
GTCCGGTTTTCAACAAACCATGCAAATGCTGAATGAGGGCATCGTTCCCACTGCGATGCTGGTTGCCAACGATCAGATGGCGCTGGGCGCAATGCGCGCCATTACCGAGTCCGGGCTGC
GCGTTGGTGCGGATATCTCGGTAGTGGGATACGACGATACCGAAGACAGCTCATGTTATATCCCGCCGTTAACCACCATCAAACAGGATTTTCGCCTGCTGGGGCAAACCAGCGTGGAC
CGCTTGCTGCAACTCTCTCAGGGCCAGGCGGTGAAGGGCAATCAGCTGTTGCCCGTCTCACTGGTGAAAAGAAAAACCACCCTGGCGCCCAATACGCAAACCGCCTCTCCCCGCGCGTT
GGCCGATTCATTAATGCAGCTGGCACGACAGGTTTCCCGACTGGAAAGCGGGCAGTGAGCGCAACGCAATTAATGTAAGTTAGCTCACTCATTAGGCACCGGGATCTCGACCGATGCCC
TTGAGAGCCTTCAACCCAGTCAGCTCCTTCCGGTGGGCGCGGGGCATGACTATCGTCGCCGCACTTATGACTGTCTTCTTTATCATGCAACTCGTAGGACAGGTGCCGGCAGCGCTCTG
GGTCATTTTCGGCGAGGACCGCTTTCGCTGGAGCGCGACGATGATCGGCCTGTCGCTTGCGGTATTCGGAATCTTGCACGCCCTCGCTCAAGCCTTCGTCACTGGTCCCGCCACCAAAC
GTTTCGGCGAGAAGCAGGCCATTATCGCCGGCATGG

**pEXK4pSAM001** GTGGGCGATCGCTCTAGAGCTAGCGAATTCCTGTGTGAAATTGTTATCCGCTCACAATTCCACACAACATACGAGCCGGAAGCATAAAGTGTAAAG
CCTGGGGTGCCTAATGAGTGAGCTAACTCACATTAATTGCGTTGCGCTCACTGCCCGCTTTCCAGTCGGGAAACCTGTCGTGCCAGCTGCATTAATGAATCGGCCAACGCGCGGGGAGA
GGCGGTTTGCGTATTGGGCGCTCTTCCGCTTCCTCGCTCACTGACTCGCTGCGCTCGGTCGTTCGGCTGCGGCGAGCGGTATCAGCTCACTCAAAGGCGGTAATACGGTTATCCACAGA
ATCAGGGGATAACGCAGGAAAGAACATGTGAGCAAAAGGCCAGCAAAAGGCCAGGAACCGTAAAAAGGCCGCGTTGCTGGCGTTTTTCCATAGGCTCCGCCCCCCTGACGAGCATCACA
AAAATCGACGCTCAAGTCAGAGGTGGCGAAACCCGACAGGACTATAAAGATACCAGGCGTTTCCCCCTGGAAGCTCCCTCGTGCGCTCTCCTGTTCCGACCCTGCCGCTTACCGGATAC
CTGTCCGCCTTTCTCCCTTCGGGAAGCGTGGCGCTTTCTCATAGCTCACGCTGTAGGTATCTCAGTTCGGTGTAGGTCGTTCGCTCCAAGCTGGGCTGTGTGCACGAACCCCCCGTTCA
GCCCGACCGCTGCGCCTTATCCGGTAACTATCGTCTTGAGTCCAACCCGGTAAGACACGACTTATCGCCACTGGCAGCAGCCACTGGTAACAGGATTAGCAGAGCGAGGTATGTAGGCG
GTGCTACAGAGTTCTTGAAGTGGTGGCCTAACTACGGCTACACTAGAAGAACAGTATTTGGTATCTGCGCTCTGCTGAAGCCAGTTACCTTCGGAAAAAGAGTTGGTAGCTCTTGATCC
GGCAAACAAACCACCGCTGGTAGCGGTGGTTTTTTTGTTTGCAAGCAGCAGATTACGCGCAGAAAAAAAGGATCTCAAGAAGATCCTTTGATCTTTTCTACGGGGTCTGACGCTCAGTG
GAACGAAAACTCACGTTAAGGGATTTTGGTCATGAGATTATCAAAAAGGATCTTCACCTAGATCCTTTTAAATTAAAAATGAAGTTTTAAATCAATCTAAAGTATATATGAGTAAAAAT
ATTCCGGAATTGCCAGCTGGGGCGCCCTCTGGTAAGGTTGGGAAGCCCTGCAAAGTAAACTGGATGGCTTTCTTGCCGCCAAGGATCTGATGGCGCAGGGGATCAAGATCTGATCAAGA
GACAGGATGAGGATCGTTTCGCATGATTGAACAAGATGGATTGCACGCAGGTTCTCCGGCCGCTTGGGTGGAGAGGCTATTCGGCTATGACTGGGCACAACAGACAATCGGCTGCTCTG
ATGCCGCCGTGTTCCGGCTGTCAGCGCAGGGGCGCCCGGTTCTTTTTGTCAAGACCGACCTGTCCGGTGCCCTGAATGAACTGCAGGACGAGGCAGCGCGGCTATCGTGGCTGGCCACG
ACGGGCGTTCCTTGCGCAGCTGTGCTCGACGTTGTCACTGAAGCGGGAAGGGACTGGCTGCTATTGGGCGAAGTGCCGGGGCAGGATCTCCTGTCATCCCACCTTGCTCCTGCCGAGAA
AGTATCCATCATGGCTGATGCAATGCGGCGGCTGCATACGCTTGATCCGGCTACCTGCCCATTCGACCACCAAGCGAAACATCGCATCGAGCGAGCACGTACTCGGATGGAAGCCGGTC
TTGTCGATCAGGATGATCTGGACGAAGAGCATCAGGGGCTCGCGCCAGCCGAACTGTTCGCCAGGCTCAAGGCGCGCATGCCCGACGGCGAGGATCTCGTCGTGACCCATGGCGATGCC
TGCTTGCCGAATATCATGGTGGAAAATGGCCGCTTTTCTGGATTCATCGACTGTGGCCGGCTGGGTGTGGCGGACCGCTATCAGGACATAGCGTTGGCTACCCGTGATATTGCTGAAGA
GCTTGGCGGCGAATGGGCTGACCGCTTCCTCGTGCTTTACGGTATCGCCGCTCCCGATTCGCAGCGCATCGCCTTCTATCGCCTTCTTGACGAGTTCTTCTGAACCGGTAATATTATTG
AAGCATTTATCAGGGTTATTGTCTCATGAGCGGATACATATTTGAATGTATTTAGAAAAATAAACAAATAGGGGTTCCGCGCACATTTCCCCGAAAAGTGCCACCTGACGTCTAAGAAA
CCATTATTATCATGACATTAACCTATAAAAATAGGCGTATCACGAGGCCCTTTCGTCTCGCGCGTTTCGGTGATGACGGTGAAAACCTCTGACACATGCAGCTCCCGGAGACGGTCACA
GCTTGTCTGTAAGCGGATGCCGGGAGCAGACAAGCCCGTCAGGGCGCGTCAGCGGGTGTTGGCGGGTGTCGGGGCTGGCTTAACTATGCGGCATCAGAGCAGATTGTACTGAGAGTGCA
CCAATTGGTCGACCTCGAGTTAATTAACGTAACGGCCAGTATGAGCAAAACTAATTTCTATCCTTTCTATTGCCAAAGCTAATATGGTCTTGGATTTTTCAAAGGCAAGGGAATCAGTA
TAAATGAACATATTTATAATTCCTTAAGAAATATAATATAAGAAATATGAAACCTGACTGAATTATAAATTGTAGATTGTAAGATAGTTAGTTTTATAAAACAATCTATTTATCCAGGC
AGCTAGACTTCTAGGCCTGATGCAAAAGGGGGGCACCCTGTTATTTCTCTATCAGAATTGGTCAGCTAGAGCCCTACCCCTGATGTAGCCACAAAGCCCCAGAACCTCACTTAGCCATG
TTTCATTTGGCTATTTGTTGAACTTTAACCCTGTTCTCTGCTTACAGAGCTAGAACCAAAGGGCTTCTTAACCGTAACTAAATGTTACTTCTAGTCTGTATGAAATAGTCCAGCAGGAG
TCCAACTGTTGGGGCAACAACGTAAAGAGGAGTATTTACTGTGCAGGGTCACAATCACTCCCTCTTAGGAAGGGATTATATGGAGTCTTTCAAAGGACAATATTGCGCACATCCTCACA
AGCACGACTCTCACAGCATCTTAAGTTGATGTACTACAGAGAGATGCTGGGCAACAATCTACTCCAAGACTATCTGCCTGGACTATAACTGAGTCTAACAGTCCACTGCTGGAAATATC
TTCCCCTACATAAAGGAATTTTCCCGCACGCTTGCTTTCAGGAAAATTTTTCTGTATAATGTCTCCATAAATTTGAGAGAGAGTTGATCAATGGCTGGTTCTCGCGAGAATTCCGAATA
GCCATCCCAATCGAACAGGGCCTGCTGGTAATCGCAGGCCTTTTTATTTCCTCAGCGAGTTTAAAATCTCCATATAAAACTGATTAAATCAGAAGAAATCATCTTCTTAAGTTGTTTCC
TTTGACAAGTTGCTGTGAATTTGTTTCAAAATGGTTAAATATTCAAATTCAAGTCGACGTTCTTAAAATGAAGAGTGAGCCTTAAGTCACAAAACCCCTGTGGGTAATGTGATGATTAG
TCGGATTTAAATGCTGGTTATTTTAGTAAATGTGAGAGTAACTAAATATATTCAAAATTTAAAATTTTCCATAATACCAAATTACAAGATACTATGCATAGGAGAGAGGGTCTAAAAGG
ACATTCATTTAAACCTTGTTTACTTTGCCCTCACAGAAACTTTTCCGGAAGGACCACATACCTATTGTGCTTAGTTAAAAATTAAAGCTGTAAGCCACCCACATCTGTCGCAAGTGCAG

CATATGATTAATGGCACATGTATTTTCCCTTCTCTTCTATTTTGTGCAGATGTTTTCCCTTGAGTTTTTGACCACTGGAGAGCTCAGCATGCCCATTGTTCCTGGCACAAGTTGGCTTG
TTCTCTTTCCCCTGTGGCTTAAATTCCTGTGCACAGTTTTCTTCCTAGGTATAAGAGTACAAGCTGTCACTTAGCCTACCTGGACTCAACTATGGGAGGTAACAAGTCCTAGGACACAA
AGGATAAGAAGTTCCTGGCCTGAGAACTGGGTTGACCTAGCAAACATGATTGAGAGCTTGAATCCTGAGATAGGCAAGGCTGAAGAACCAGGACTAAGGTTCAGAAGTTCTGGTGTCCA
AGACAAAACTGACATTAAGTATAAGTGTATGGGTTCAGTGCTCAAGCTAACTTAAATCTAGTTTCCAGAAAGGCAAGCCTGCATCTGAAGTGTGCCCCTGACAGGTTTGAAGCATTTTT
CCCCTTTGAAACACGTGATACAACAAATATGATTTTCTCCCTCTCCATATTCACTAATATCTGGCCCAGGTTTTTAACCCCATGAAGTTATGGGAAGGCCATTGAATTCTAAACAAGCT
TTCTCCAGTCAGTCCTGGACATTGTATTGAGTGATTTAGGCACATCTATGCAGATAGGTGACCTGGTAGTATCAGTGAGTATATATCACAAAATCGACTATGTTTAACAAATAGGCTGA
AGAGAAGACCCTTAGG

#### Primers for PCR reactions involving substrates in isolation

| | |
|---|---|
| **OL082** | GATTGCACGCAGGTTCTCC |
| **OL090** | CAATGGCCTTCCCATAAC |
| **OL092** | ATGCGAGGTCTCGAGGATTATCCAGGCAGCTAGAC |
| **OL093** | ATGCGAGGTCTCGCCGCTTATCCAGGCAGCTAGAC |
| **OL094** | ATGCGAGGTCTCGCACCTTATCCAGGCAGCTAGAC |
| **OL097** | ATAGTAGGTCTCGTCCTATGGGTCACGACGAGATCC |
| **OL098** | ATAGTAGGTCTCGGCGGATGGGTCACGACGAGATCC |
| **OL099** | ATAGTAGGTCTCGGGTGATGGGTCACGACGAGATCC |
| **OL102** | ATGCGAGGTCTCGTTCTTTATCCAGGCAGCTAGAC |
| **OL103** | ATGCGAGGTCTCGCCGCCTATCCAGGCAGCTAGAC |
| **OL105** | ATAGTAGGTCTCGAGAAATGGGTCACGACGAGATCC |
| **OL106** | ATAGTAGGTCTCGGCGGCTGGGTCACGACGAGATCC |
| **OL110** | ATGCGAGGTCTCGAGAATTATCCAGGCAGCTAGAC |
| **OL111** | ATAGTAGGTCTCGTTCTATGGGTCACGACGAGATCC |
| **OL131** | ATAGTAGGTCTCAGCGGATGGGTCACGACGAGATCC |
| **OL132** | ATGCGAGGTCTCTCCGCTTATCCAGGCAGCTAGAC |
| **OL135** | ATAGTAGGTCTCATTCTATGGGTCACGACGAGATCC |
| **OL136** | ATGCGAGGTCTCTAGAATTATCCAGGCAGCTAGAC |

#### Primers for PCR reactions involving the assembly of 10 fragments (optimal set)

| ID | Sequence | Overhang, BsaI-HFv2 digestion |
|---|---|---|
| **OL1388** | CGCAGGTCTCAGGAGATTATTTTTGACACCAGACC | GGAG |
| **OL1457** | CAATAGTAGGTCTCGACCCCGTACGTCTTCCCGAG | ACCC |
| **OL1458** | CATACGCAGGTCTCAGGGTATACATGTCTGACAAT | GGGT |
| **OL1459** | CAATAGTAGGTCTCGAAGCAGCGTTGTTGCAGTGC | AAGC |
| **OL1460** | CATACGCAGGTCTCAGCTTCGGCCTGGTAATGGCC | GCTT |
| **OL1461** | CAATAGTAGGTCTCGAGCAGTGGCGTCTGGCTGAA | AGCA |
| **OL1462** | CATACGCAGGTCTCATGCTGCCAGGCGCTGATGTG | TGCT |
| **OL1463** | CAATAGTAGGTCTCGGACCGCACGCCGCATCCAGC | GACC |
| **OL1464** | CATACGCAGGTCTCAGGTCGGCAAAGACCAGACCG | GGTC |
| **OL1465** | AATAGTAGGTCTCACCCGGCTGTGCCGAAATGGTC | CCCG |
| **OL1466** | CATACGCAGGTCTCACGGGAAGGGCTGGTCTTCAT | CGGG |
| **OL1467** | CAATAGTAGGTCTCGTGGATGAAGCCAATATTGAA | TGGA |
| **OL1468** | CATACGCAGGTCTCATCCACCACATACAGGCCGTA | TCCA |
| **OL1469** | CAATAGTAGGTCTCGGTGCGGTGGTTGAACTGCAC | GTGC |
| **OL1470** | CATACGCAGGTCTCAGCACGATAGAGATTCGGGAT | GCAC |
| **OL1471** | CAATAGTAGGTCTCGGCGGCATTTTCCGTGACGTC | GCGG |
| **OL1472** | CATACGCAGGTCTCACCGCTCATCCGCCACATATC | CCGC |
| **OL1473** | CAATAGTAGGTCTCGAACGTGACCTATCCCATTAC | AACG |
| **OL1492** | CATACGCAGGTCTCACGTTGGTGTAGATGGGCGCA | CGTT |
| **OL1387** | ATAGTAGGTCTCGATGGGGCCGCCATGCCGGCGAT | ATGG |

#### Primers for PCR reactions involving the assembly of 10 fragments (non-optimal set)

| ID | Sequence | Overhang, BsaI-HFv2 digestion |
|---|---|---|
| **OL1388** | CGCAGGTCTCAGGAGATTATTTTTGACACCAGACC | GGAG |
| **OL1475** | CAATAGTAGGTCTCGATCTGCCATTGTCAGACATG | ATCT |
| **OL1476** | ATACGCAGGTCTCAAGATCCCAGCGGTCAAAACAG | AGAT |
| **OL1477** | CAATAGTAGGTCTCGATTACCAGGCCGAAGCAGCG | ATTA |
| **OL1478** | CATACGCAGGTCTCATAATGGCCCGCCGCCTTCCA | TAAT |
| **OL1479** | ATAGTAGGTCTCGTCAGCGCCTGGCAGCAGTGGCG | TCAG |
| **OL1480** | CATACGCAGGTCTCACTGATGTGTCCGGCTTCTGA | CTGA |
| **OL1481** | CAATAGTAGGTCTCGTGACGGAAGCAAAACACCAG | TGAC |
| **OL1482** | CATACGCAGGTCTCAGTCAGCGCTGGATGCGGCGT | GTCA |
| **OL1483** | CAATAGTAGGTCTCGTCCCGGCTGTGCCGAAATGG | TCCC |
| **OL1484** | CATACGCAGGTCTCAGGGAAGGGCTGGTCTTCATC | GGGA |
| **OL1485** | CAATAGTAGGTCTCGAAGCCAATATTGAAACCCAC | AAGC |
| **OL1486** | CATACGCAGGTCTCAGCTTCATCCACCACATACAG | GCTT |
| **OL1487** | CAATAGTAGGTCTCGCGAATCTCTATCGTGCGGTG | CGAA |
| **OL1488** | CATACGCAGGTCTCATTCGGGATTTCGGCGCTCCA | TTCG |
| **OL1489** | CAATAGTAGGTCTCGTTTTCCGTGACGTCTCGTTG | TTTT |
| **OL1490** | CATACGCAGGTCTCAAAAAATGCCGCTCATCCGCCA | AAAA |
| **OL1491** | CAATAGTAGGTCTCGAACGTGACCTATCCCATTAC | AACG |
| **OL1492** | CATACGCAGGTCTCACGTTGGTGTAGATGGGCGCA | CGTT |
| **OL1387** | ATAGTAGGTCTCGATGGGGCCGCCATGCCGGCGAT | ATGG |

### References


[1] John M Pryor, Vladimir Potapov, Rebecca B Kucera, Katharina Bilotti, Eric J Cantor, and Gregory J S Lohman. Enabling one-pot golden gate assemblies of unprecedented complexity using data-optimized assembly design. *PLoS One*, 15(9):e0238592, 2020.

[2] Vladimir Potapov, Jennifer L Ong, Rebecca B Kucera, Bradley W Langhorst, Katharina Bilotti, John M Pryor, Eric J Cantor, Barry Canton, Thomas F Knight, Thomas C Evans, Jr, and Gregory J S Lohman. Comprehensive profiling of four base overhang ligation fidelity by t4 dna ligase and application to dna assembly. *ACS Synth Biol*, 7(11):2665–2674, 11 2018.

[3] Alfred Pingoud and Albert Jeltsch. Structure and function of type ii restriction endonucleases. *Nucleic acids research*, 29(18):3705–3727, 2001.

[4] Linda Jen-Jacobson. Protein—dna recognition complexes: Conservation of structure and binding energy in the transition state. *Biopolymers: Original Research on Biomolecules*, 44(2):153–180, 1997.

[5] Nick Kamps-Hughes, Aine Quimby, Zhenyu Zhu, and Eric A Johnson. Massively parallel characterization of restriction endonucleases. *Nucleic acids research*, 41(11):e119–e119, 2013.

[6] Waclaw Szybalski, Sun C Kim, Noaman Hasan, and Anna J Podhajska. Class-iis restriction enzymes—a review. *Gene*, 100:13–26, 1991.

[7] Sylvestre Marillonnet and Ramona Grützner. Synthetic dna assembly using golden gate cloning and the hierarchical modular cloning pipeline. *Current protocols in molecular biology*, 130(1):e115, 2020.

[8] J SantaLucia, Jr. A unified view of polymer, dumbbell, and oligonucleotide dna nearest-neighbor thermodynamics. *Proc Natl Acad Sci U S A*, 95(4):1460–5, Feb 1998.

[9] Ekaterina Protozanova, Peter Yakovchuk, and Maxim D Frank-Kamenetskii. Stacked-unstacked equilibrium at the nick site of dna. *J Mol Biol*, 342(3):775–85, Sep 2004.

[10] Arren Bar-Even, Elad Noor, Yonatan Savir, Wolfram Liebermeister, Dan Davidi, Dan S Tawfik, and Ron Milo. The moderately efficient enzyme: evolutionary and physicochemical trends shaping enzyme parameters. *Biochemistry*, 50(21):4402–10, 2011.

[11] F M Pohl, R Thomae, and A Karst. Temperature dependence of the activity of dna-modifying enzymes: endonucleases and dna ligase. *Eur J Biochem*, 123(1):141–52, 1982.

[12] Peter Yakovchuk, Ekaterina Protozanova, and Maxim D. Frank-Kamenetskii. Base-stacking and base-pairing contributions into thermal stability of the DNA double helix. *Nucleic Acids Research*, 34(2):564–574, 01 2006.

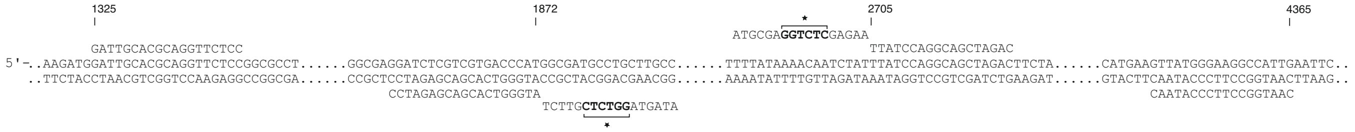


Supplementary Figure S1: DNA sequences showing the local binding of primers on the plasmid (`TAGAAT` sequence found in the $\alpha\beta$ product).

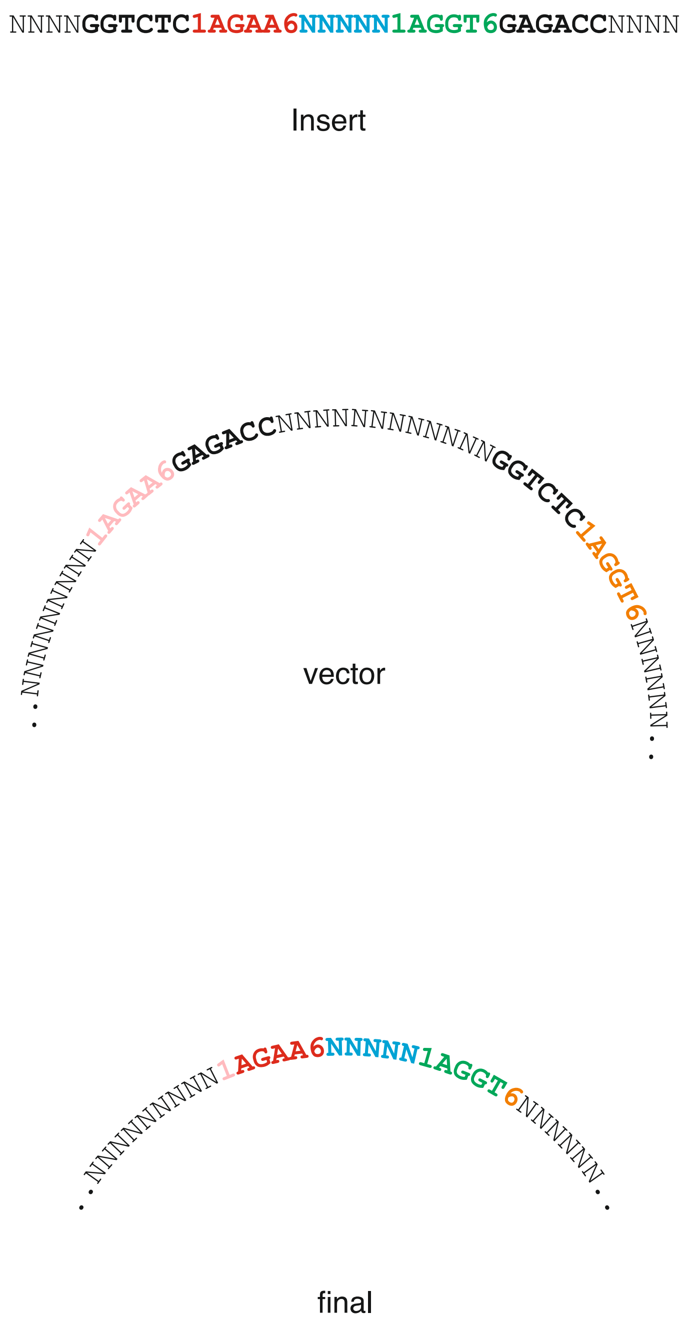


Supplementary Figure S2: A typical GGA experiment where a fragment is inserted into a vector using BsaI-HFv2 (recognizing the `GGTCTC` sequence). The vector has given nucleotides 1 and 6 that determine stacking energies (and possibly different than that found in the insert). Shown here are sens strands only.

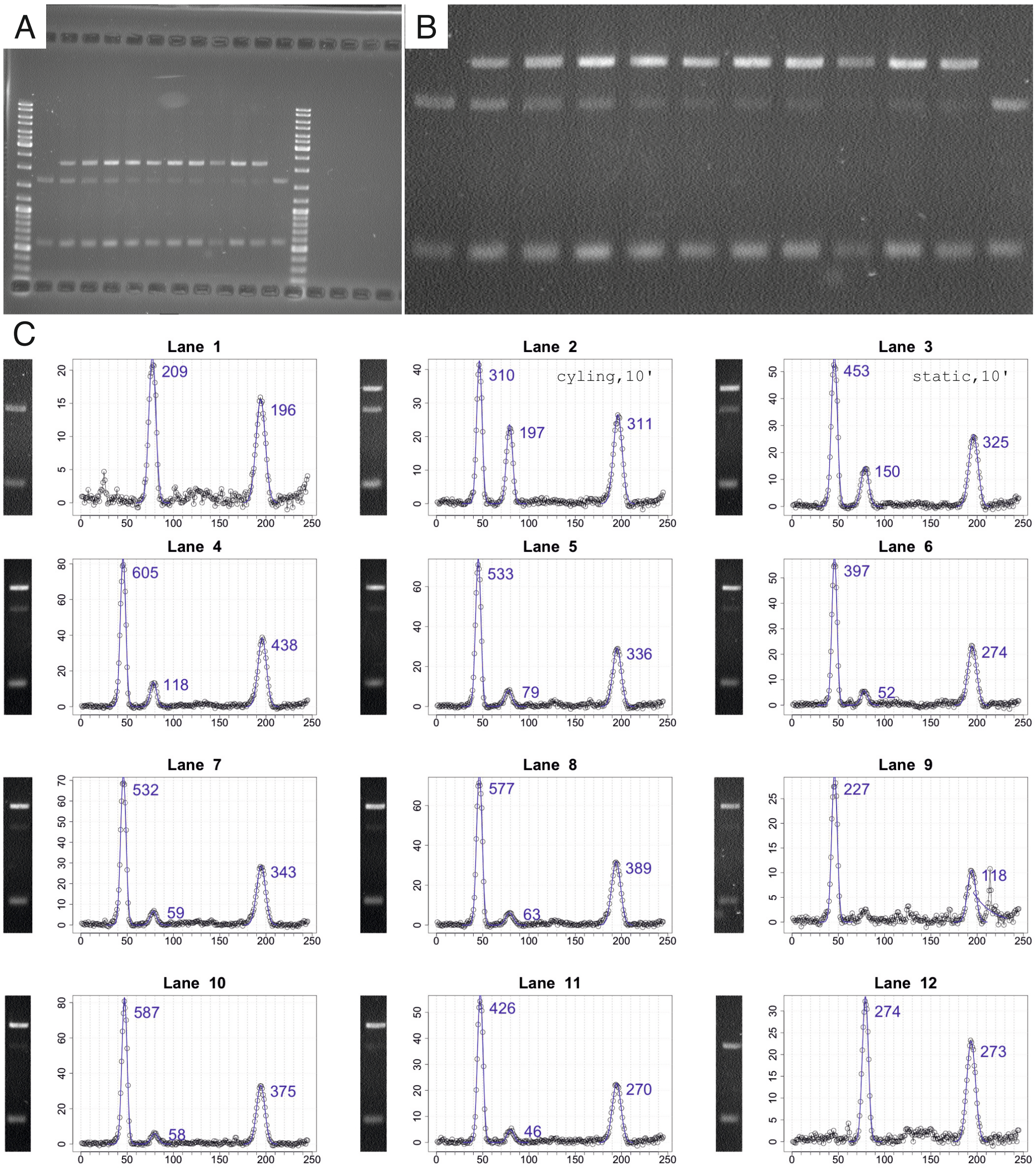


Supplementary Figure S3: A. A typical gel obtained after a GGA experiment (14 lanes), corresponding to the data shown in Figure 2 of the main text (`GCCGCC`, black symbols). B. Before the automatic analysis, the raw image is manually cropped to remove the ladders. C. The R routine automatically finds the different lanes (12 in total, see protocol page 12 of this document). We then determine the intensity of each band by fitting a gaussian to the projected image (along the axis perpendicular to the direction of migration, which allows to remove the noise and evidence for minute changes in intensity). Values on the $x$ axis are expressed in pixels, values on the $y$ axis denote intensities (arbitrary units). Due to the presence of artifacts (e.g. debris in lane 9), some manual processing is sometimes needed. Lanes 1 and 12 allow to calculate the stoichiometry $S_\beta$ as $100 \times \beta'\beta/(\beta'\beta + \alpha\alpha')$ where reactants are expressed in intensity (mass). Here, we find values of 209/(209+196)$\sim$51.6 and 274/(274+273)$\sim$50.1 which gives an average of $\sim$50.8 and an error of $\sim$0.8 (the half difference between the two calculated stoichiometries). Because $\alpha\alpha'$ is 565 bp and $\beta'\beta$ is 1678 bp, a theoretical stoichiometry of 3 (that of the experiments shown in Figure 2 of the main text for $[\alpha\alpha'] = 21$ nM and $[\beta'\beta] = 7$ nM) yields an experimental stoichiometry of 100/(1+3$\times$(565/1678)) or 49.7. The percentage of product formed is calculated using $100 \times \alpha\beta/(\alpha\beta + \beta'\beta \times \gamma)$, with $\gamma=$ (565+1678)/1678 . For Lane 2 (cycling, 10 minutes), this gives a value of 100/(1+(197/310)$\times\gamma$) or $\sim$54.1.

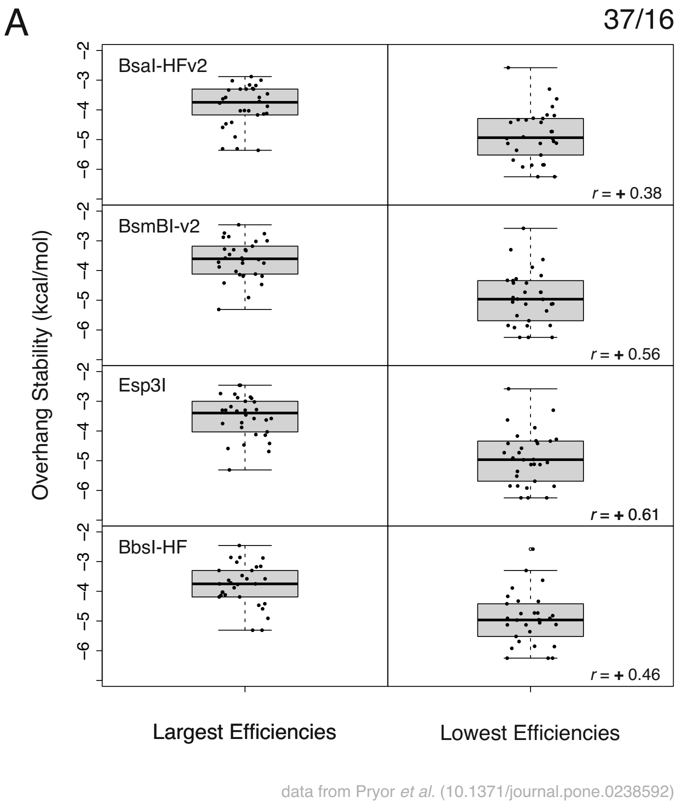


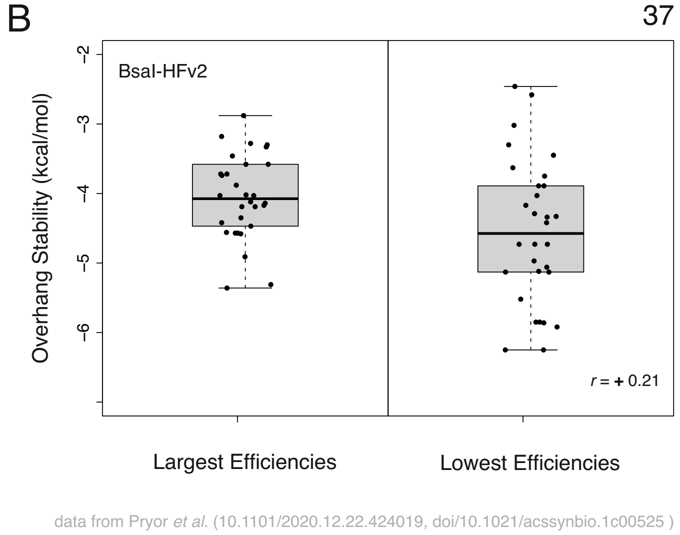


Supplementary Figure S4: Boxplots showing the relation between ligation efficiencies obtained by high throughput GGA experiments [1] and overhang stabilities (determined from [8]). Here the 30 largest (left panels) and lowest ligation (right panels) efficiency values were selected among 120 values: among the $4^4 = 256$ efficiency values, we removed palindromic and non-discernible sequences (e.g. `TAAC` and `GTTA` sequences are not discernible in those high throughput assays as they are complementary to each other) for each enzyme (BsaI-HFv2, BsmBI-v2, Esp3I and BbsI-HF). Ligation values (as .xlsx files) given in the Supplementary Information of [1] have been used. Also given are corresponding Pearson coefficients (denoted as r on the graph, calculated for the 120 values) between the ligation efficiencies and the overhang stabilities. The values are all positive suggesting that ligation efficiencies (as reported in high throughput experiments) may negatively correlate with thermodynamics data : weak overhangs show a larger ligation efficiencies at concentrations used in these assays (less than 1 nM). A. Cycling the temperature (5 minutes at 37°C, 5 minutes at 16°C). B. Static temperature of 37°C.

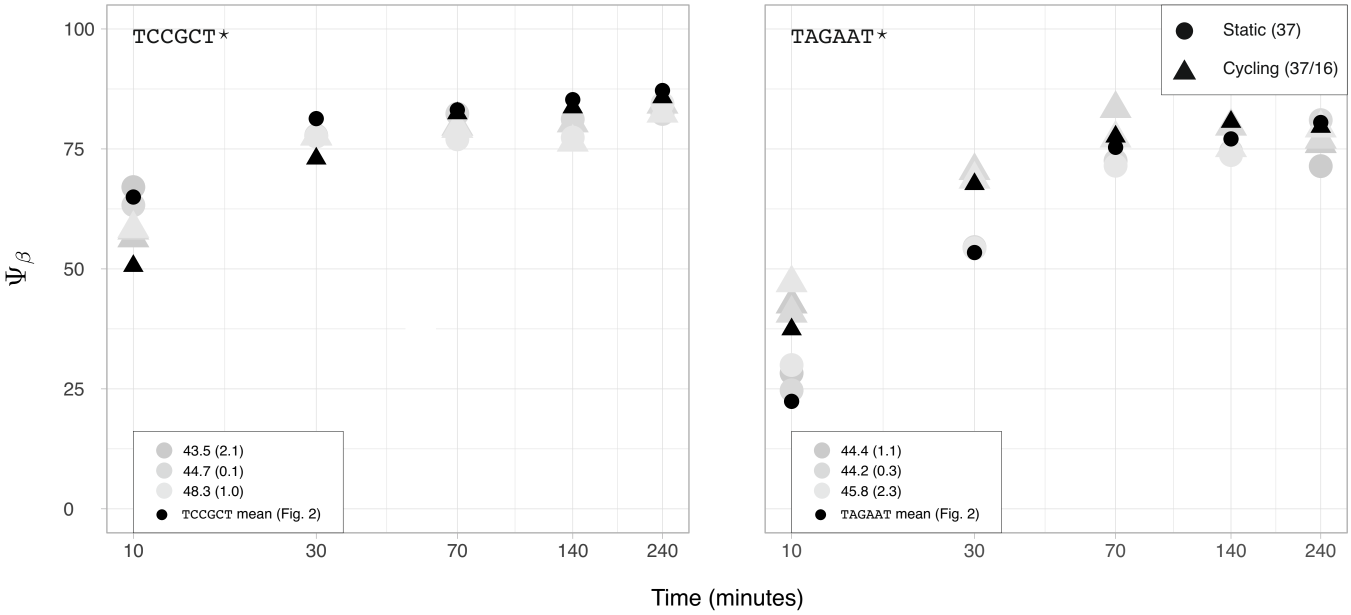


Supplementary Figure S5: Golden-gate assembly using conversed nucleotides in positions 1 and 6 for a given sequence (denoted as $\star$). Experiments for TCCGCT$^\star$ and TAGAAT$^\star$ sequences, where nucleotides in positions 1 and 6 are conserved through possible intermediates ($\alpha{\cdot}\beta$, $\alpha{\cdot}\alpha'$, $\beta'{\cdot}\beta$ and $\beta'{\cdot}\alpha'$, see Figure 3 of the main text). Experiments are performed in triplicates (grey circles: static assay, upper grey triangles : cycling assay). Black symbols are experimental averaged $\Psi_\beta$ values for TCCGCT and TAGAAT (Figure 2 of the main text), *i.e.* sequences in which stacking energies are not conserved. Finally and as in Figure 2 of the main text, the ratio $S_\beta$ is also given (the value in bracket is the corresponding error).

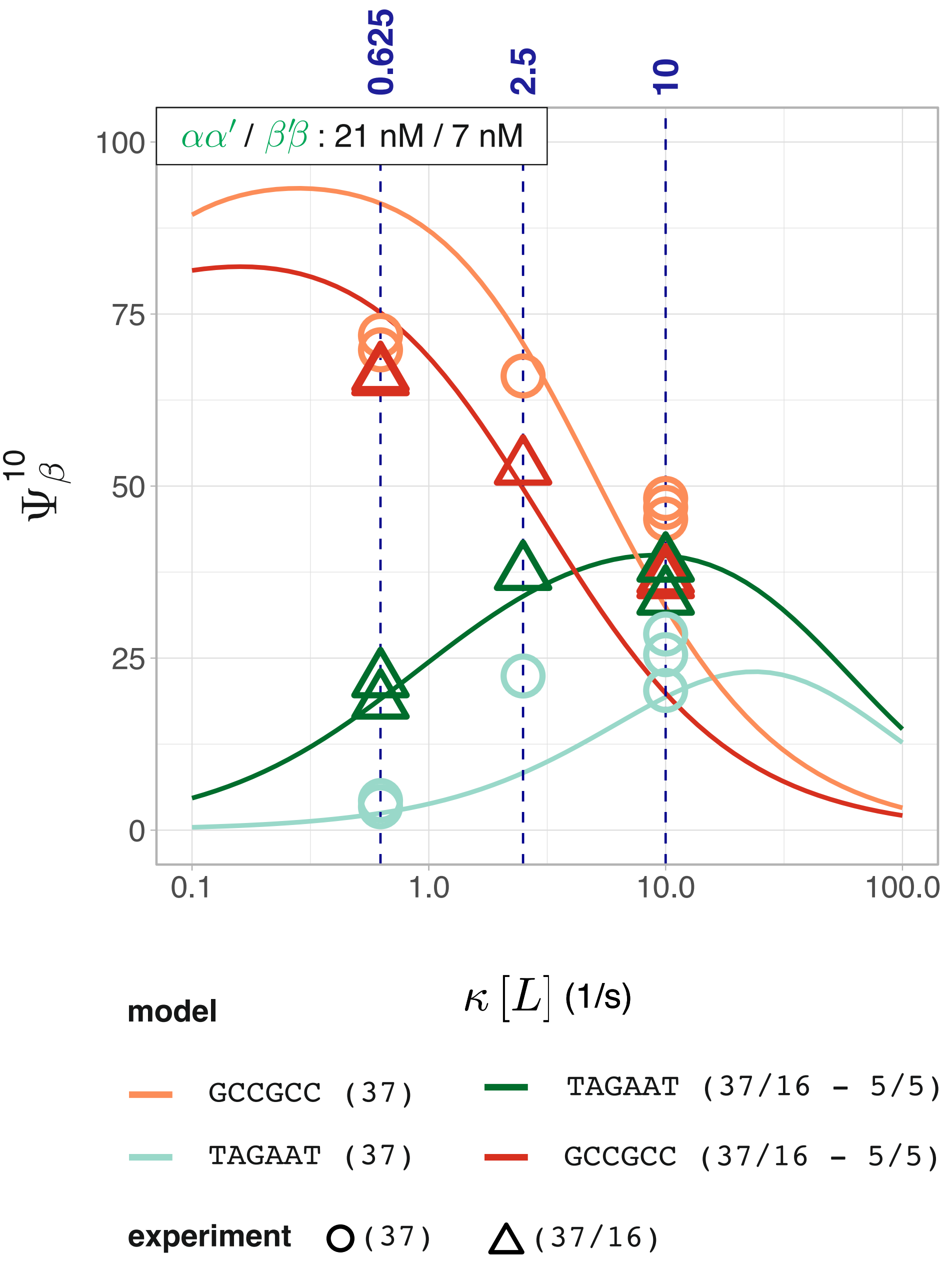


Supplementary Figure S6: The dependence of product formed $\Psi_\beta$ as a function of the ligase concentration at 10 minutes ($\Psi_\beta^{10}$) using $k_+$ values and $k_-$ values from Supplementary Table S8. Solid lines represent calculated $\Psi_\beta^{10}$ values (performed at $[\alpha\alpha'] = 21$ nM, $[\beta'\beta] = 7$ nM) for a weak (`TAGAAT`) and a strong overhang (`GCCGCC`) overhang while cycling the temperature (5 minutes at 37°C, 5 minutes at 16°C) or not (fixed temperature of 37°C). Here, parameters given in Supplementary Table S6 and S8 were used. Symbols denote experimental points, and their colors match the simulations (e.g., orange circles are experimental points performed for a strong overhang at a static temperature of 37°C). Triangles represent cycling experiments (5 minutes at 37°C, 5 minutes at 16°C), whereas circles represent static experiments (fixed temperature of 37°C). Experimental points at $\kappa[L]$=2.5 $s^{-1}$ are averaged values obtained from the data shown in Figure 2.A. Here a value of 2.5 $s^{-1}$ for $\kappa[L]$ corresponds to a final ligase concentration of 12 U/$\mu L$. This figure is identical to Figure 3.C of the main text; the only difference is that the kinetic parameters from Supplementary Table S8 have been used here. In contrast, Figure 3.C uses parameters from Supplementary Table S7.

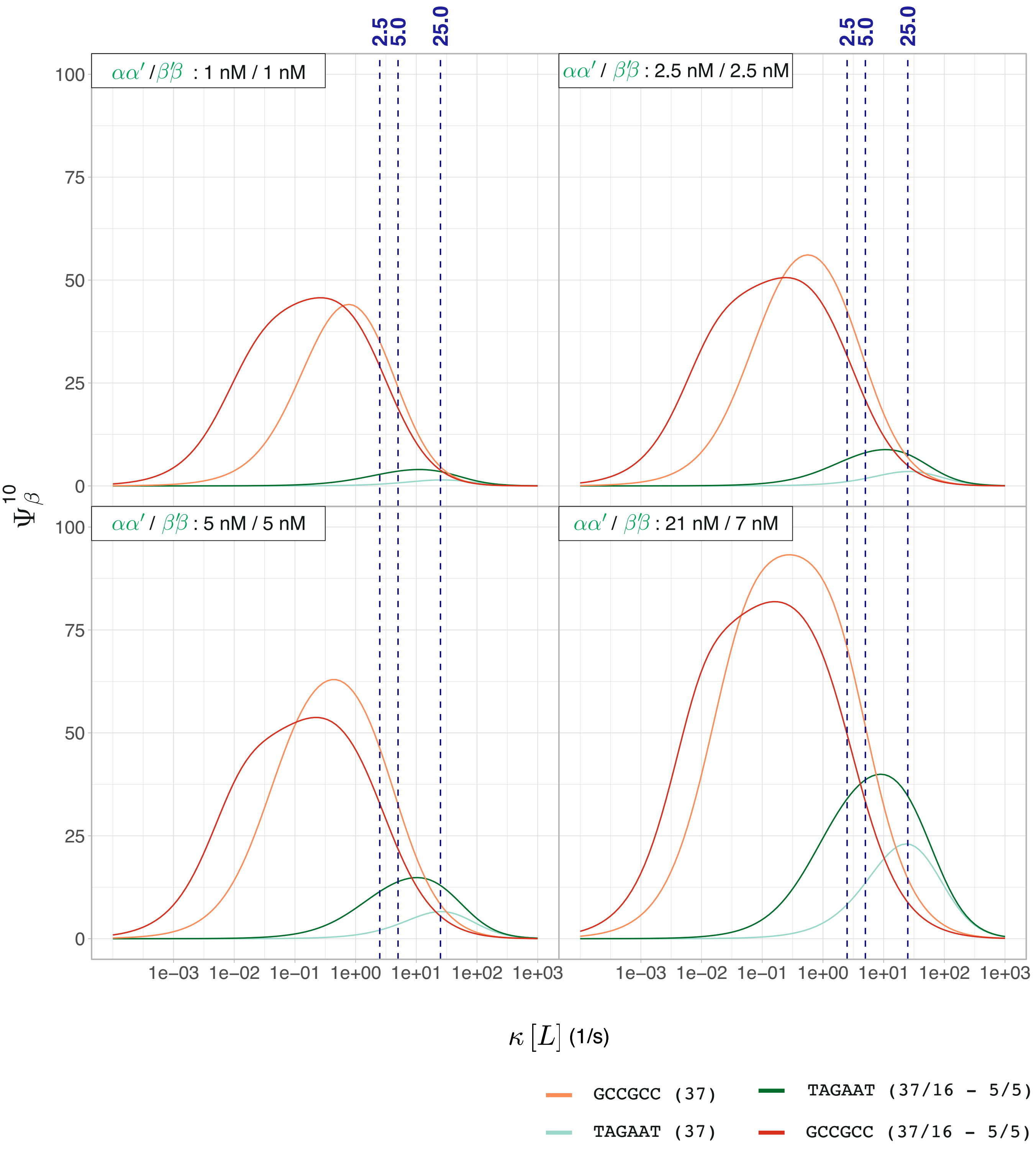


Supplementary Figure S7: Calculated $\Psi_{\beta}^{10}$ (percentage of product formed at 10 minutes) as a function of $\kappa[L]$ for different $\alpha\alpha'$ and $\beta'\beta$ concentrations ( $[\alpha\alpha']$=1, 5, 2.5, 5 nM, $[\beta'\beta]$=1, 5, 2.5, 21 nM) for a weak (`TAGAAT`) and a strong (`GCCGCC`) overhang while cycling the temperature (5 minutes at 37°C, 5 minutes at 16°C) or not (fixed temperature of 37°C). For those calculations, parameters given in Supplementary Tables S6 and S8 were used. Here a value of 2.5 $s^{-1}$ for $\kappa[L]$ corresponds to a final ligase concentration of 12 U/$\mu L$.

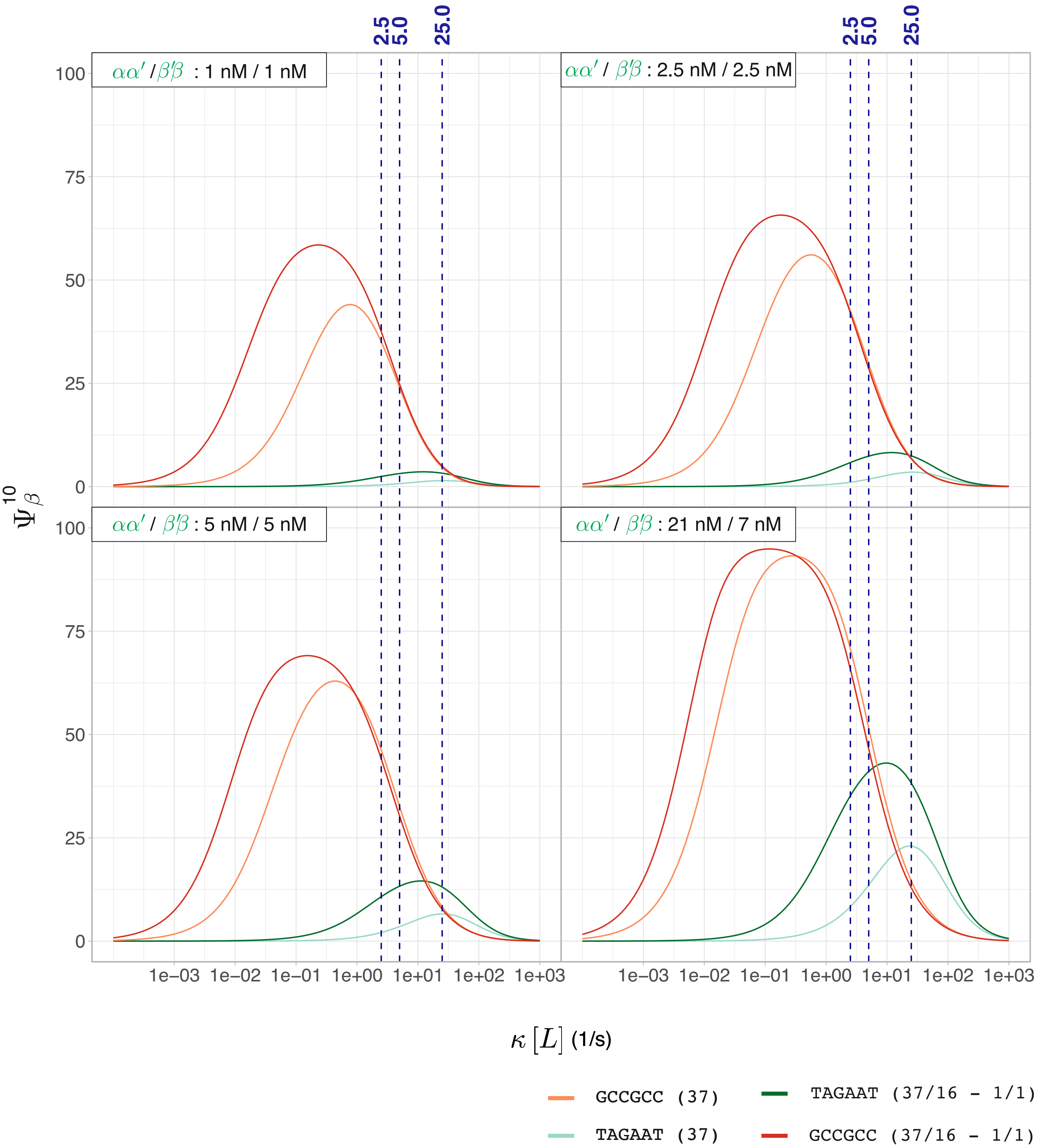


Supplementary Figure S8: Calculated $\Psi_\beta^{10}$ (percentage of product formed at 10 minutes) as a function of $\kappa[L]$ for different $\alpha\alpha'$ and $\beta'\beta$ concentrations ( $[\alpha\alpha']$=1, 5, 2.5, 5 nM, $[\beta'\beta]$=1, 5, 2.5, 21 nM) for a weak (`TAGAAT`) and a strong (`GCCGCC`) overhang while cycling the temperature (1 minute at 37°C, 1 minute at 16°C) or not (fixed temperature of 37°C). For those calculations, parameters given in Supplementary Tables S6 and S8 were used. Here a value of 2.5 $s^{-1}$ for $\kappa[L]$ corresponds to a final ligase concentration of 12 U/$\mu L$. This figure is identical to Supplementary Figure S7, except that the cycling duration has been altered from 5 minutes to 1 minute.

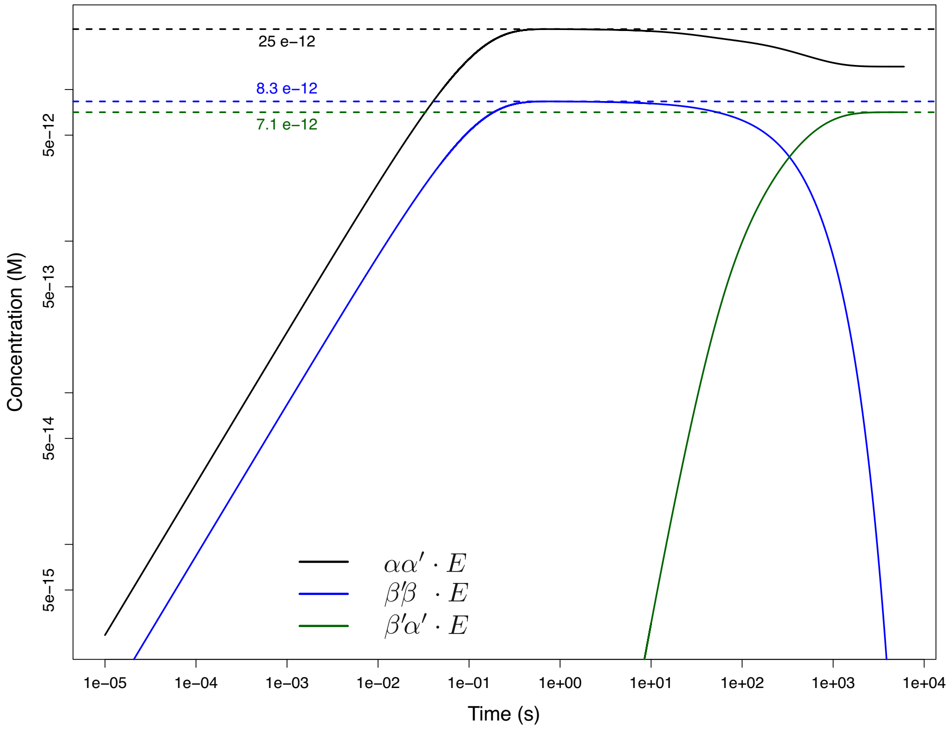


Supplementary Figure S9: The time-dependence of all three intermediates bound to the restriction endonuclease ($\alpha\alpha' \cdot E$, $\beta'\beta \cdot E$ and $\beta'\alpha' \cdot E$). Simulations performed at a static temperature of 37°C and for $[\alpha\alpha'] = 21$ nM, $[\beta'\beta] = 7$ nM. Parameters are given in Supplementary Tables S6 and S7. As stated in the main text, we have assumed the concentration of enzymes to be constant. This approximation holds as long as the concentration of the free enzyme is comparable to that of the total enzyme. Taking a value of 7 $\mu$M for the stock concentration of T4 DNA ligase [2], we obtain a value of 210 nM in our assay (see Supplementary Text and Protocols). As this is much larger than the concentration of our reactants, we can assume that the concentration of ligase is constant and close to that of the total concentration. The stock concentration of the restriction endonuclease is however not available. From the time-dependence of $\alpha\alpha' \cdot E$, $\alpha\beta' \cdot E$ as well as $\beta'\alpha' \cdot E$, we determine the maximum concentrations of those intermediates (and so of the bound enzyme). Here the total concentration of intermediates is much less $\sim$1 nM. We can then assume, to a good approximation, that our approximation holds as long as the concentration of restriction endonuclease is $\sim$1 nM.

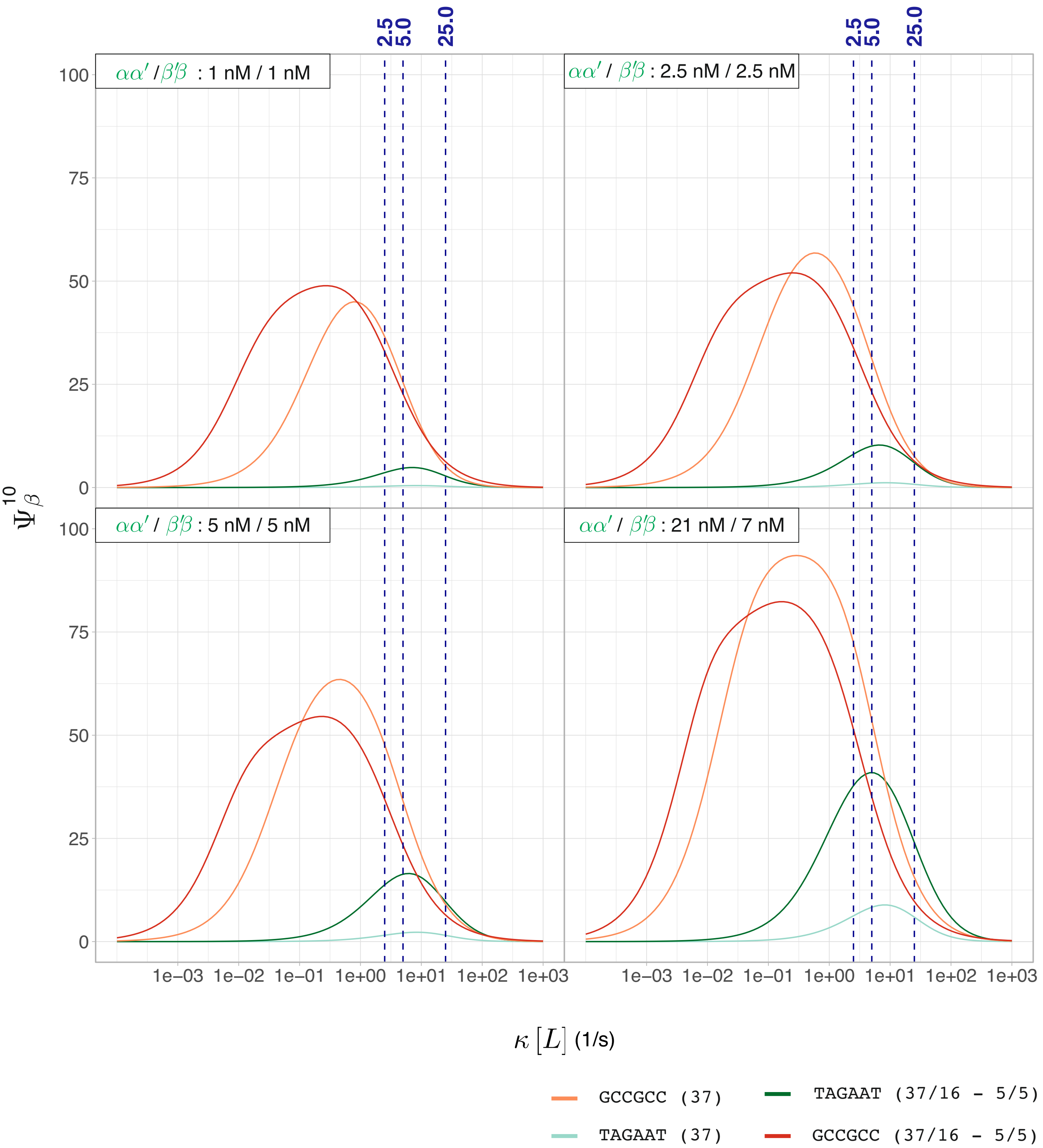


Supplementary Figure S10: Calculated $\Psi_\beta^{10}$ (percentage of product formed at 10 minutes) as a function of $\kappa[L]$ for different $\alpha\alpha'$ and $\beta'\beta$ concentrations ( $[\alpha\alpha']$=1, 5, 2.5, 5 nM, $[\beta'\beta]$=1, 5, 2.5, 21 nM) for a weak (TAGAAT) and a strong (GCCGCC) overhang while cycling the temperature (5 minutes at 37°C, 5 minutes at 16°C) or not (fixed temperature of 37°C). For those calculations, parameters given in Supplementary Tables S6 and S7 were used. Here a value of 2.5 $s^{-1}$ for $\kappa[L]$ corresponds to a final ligase concentration of 12 U/$\mu L$. This figure is identical to Supplementary Figure S7, except that parameters given in Supplementary Table S7 have been used instead of those given in Supplementary Table S8.

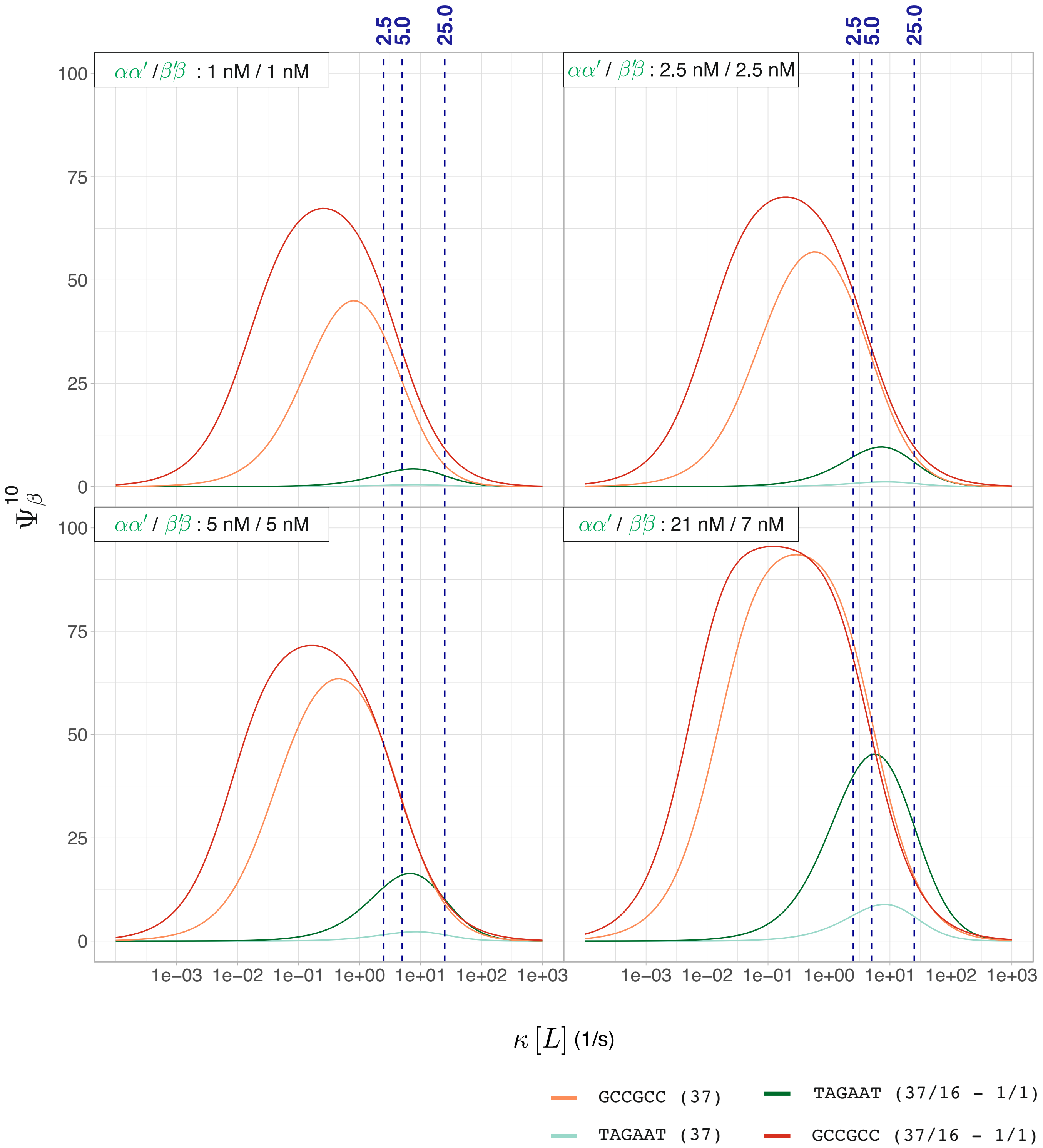


Supplementary Figure S11: Calculated $\Psi_\beta^{10}$ (percentage of product formed at 10 minutes) as a function of $\kappa[L]$ for different $\alpha\alpha'$ and $\beta'\beta$ concentrations ( $[\alpha\alpha']$=1, 5, 2.5, 5 nM, $[\beta'\beta]$=1, 5, 2.5, 21 nM) for a weak (`TAGAAT`) and a strong (`GCCGCC`) overhang while cycling the temperature (1 minute at 37°C, 1 minute at 16°C) or not (fixed temperature of 37°C). For those calculations, parameters given in Supplementary Tables S6 and S7 were used. Here a value of 2.5 $s^{-1}$ for $\kappa[L]$ corresponds to a final ligase concentration of 12 U/$\mu L$. This figure is identical to Supplementary Figure S10, except that the cycling duration has been altered from 5 minutes to 1 minute.

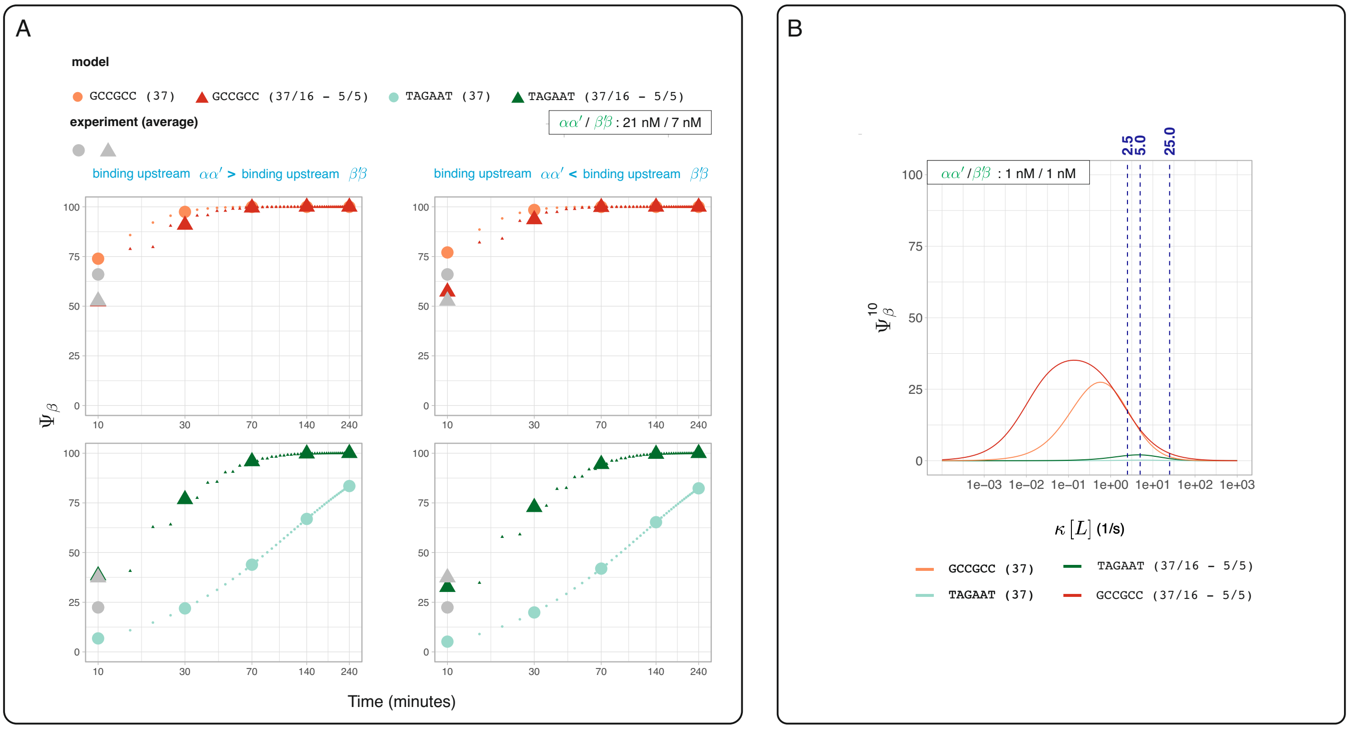


Supplementary Figure S12: A. The dependence of product formed $\Psi_\beta$ as a function of the binding efficiency of the endonuclease. This Figure is similar to Figure 3.B of the main text; except that the binding of the endonuclease was altered. When deriving differential equations, the value of $\kappa[E]$ was increased by a factor 10 for reactions involving binding to $\alpha\alpha'$ and $\beta'\alpha'$ (left panel) or for reactions involving binding to $\beta'\beta$ and $\beta'\alpha'$ (right panel). As for Figure 3.B of the main text, we prioritized experimental data obtained at 37°C/ 16°C when adjusting free parameters (gray circles and upper triangles represent the averaged of experimental values obtained for a strong overhang (`GCCGCC`) and a weak overhang (`TAGAAT`), respectively). Parameters given in Supplementary Table S6 and S7 were used with the exception that $\kappa[E]$ was found to be 0.010 $s^{-1}$ (left panel) or 0.005 $s^{-1}$ (right panel). B. After identifying these new parameter sets, we calculated the dependence of the product formed, denoted as $\Psi_\beta$, on ligase concentration at 10 minutes ($\Psi_\beta^{10}$). This calculation was performed similarly to the analysis shown in Figure 3.D of the main text. In this scenario, we assumed that the upstream sequence remained identical (consistent with sequencing-based experiments and the differential equations discussed in this document), and we updated the value of $\kappa[E]$ to 0.005 $s^{-1}$. As shown, the simulation results closely match those found in Figure 3.D of the main text. Indeed, varying $\kappa[E]$ does not significantly affect $\Delta\Psi_\beta^{10}$ (Supplementary Figure S13).

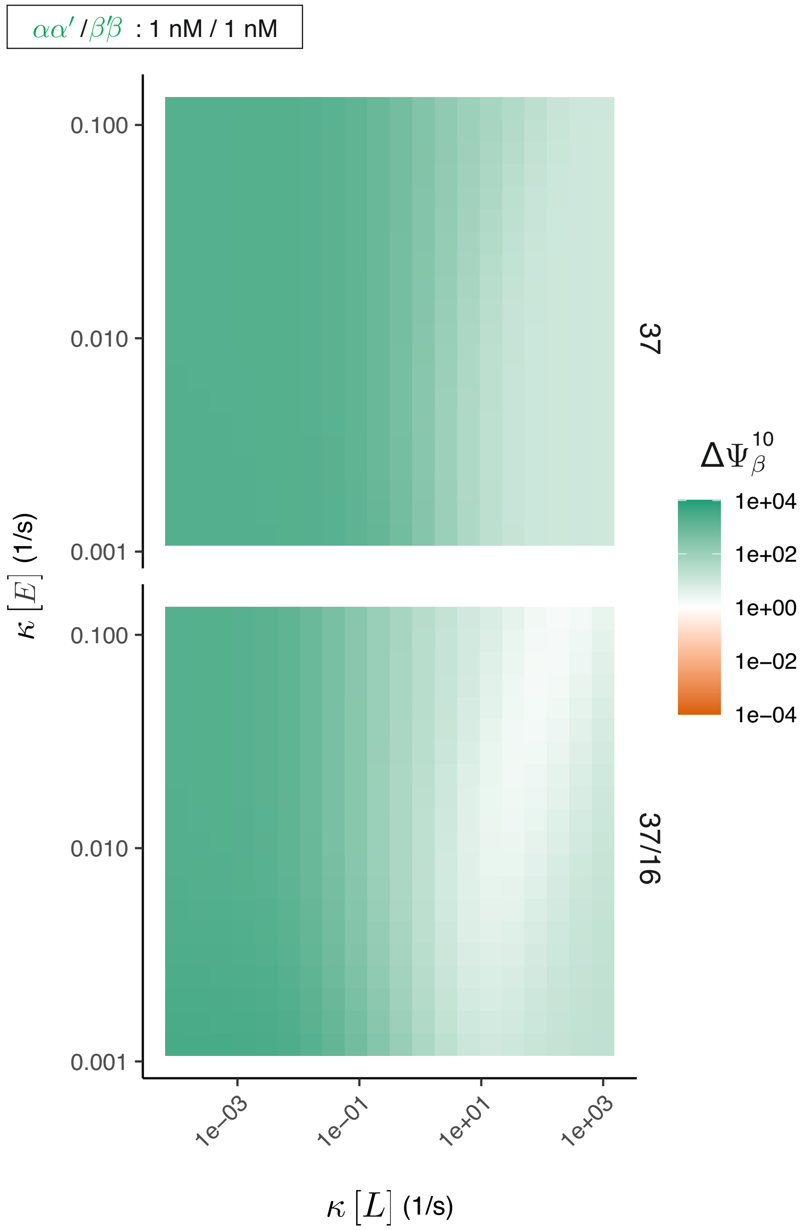


Supplementary Figure S13: Influence of the binding of the endonuclease to the GGA assembly. Here, using parameters given in Supplementary Table S6 and S7, we varied the value of $\kappa[E]$ by 2 orders of magnitude and determine the ratio in percentage of product formed $\Psi_\beta$ at 10 minutes, $\Delta\Psi_\beta^{10}$ between a strong overhang and a weak overhang (for both a cycling assay, 5 minutes at 37°C and 5 minutes at 16°C, and a static assay at 37°C assuming $\alpha\alpha'$ and $\beta'\beta'$ at 1 nM). Values greater than 1 (*i.e.*, a strong overhang shows a higher $\Psi_\beta^{10}$ value) are indicated in green (logarithmic scale). This simulation demonstrates that varying the concentration of the endonuclease (*i.e.*, the enzyme's binding capability to a given sequence) does not significantly affect $\Delta\Psi_\beta^{10}$.

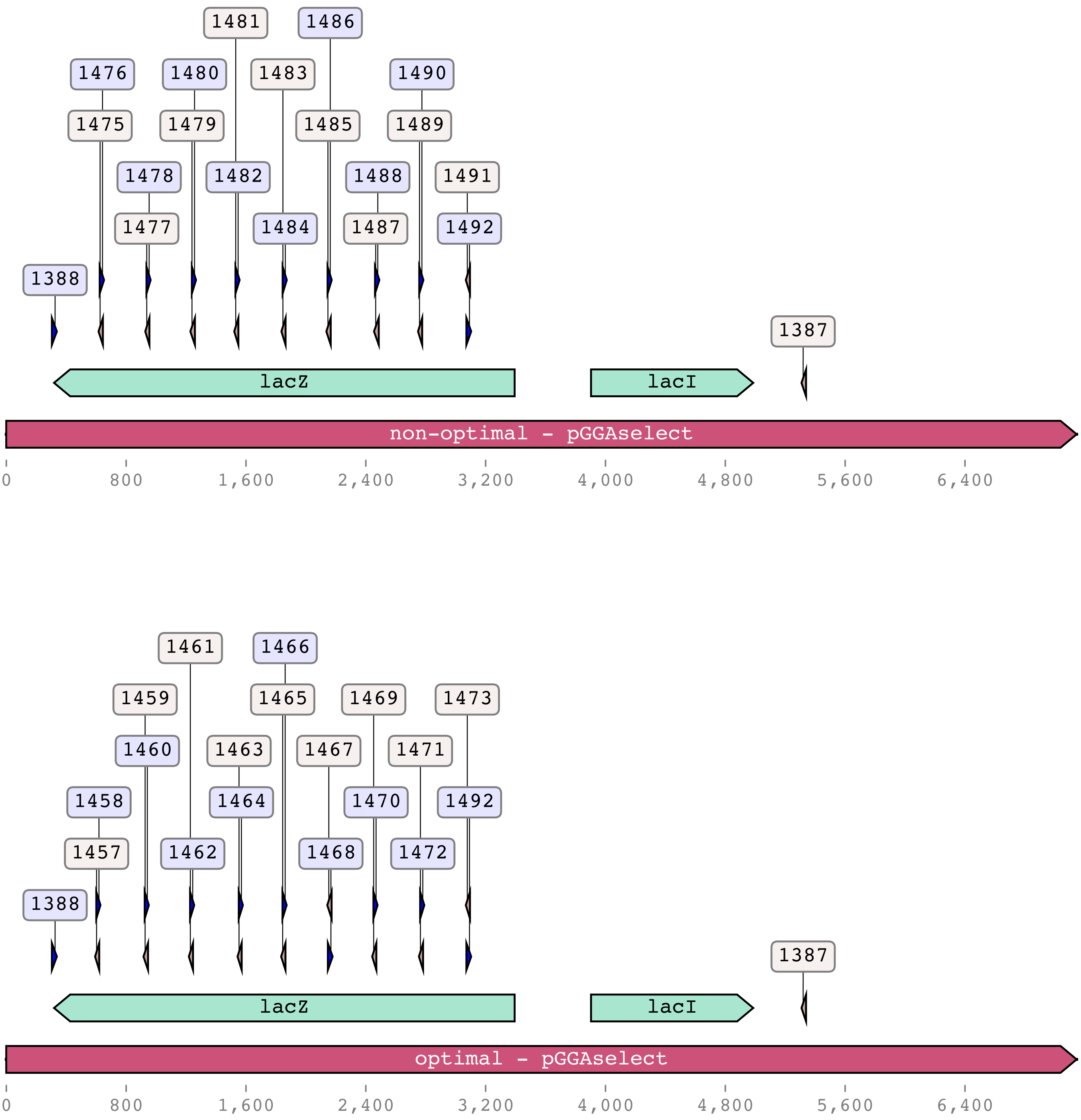


Supplementary Figure S14: Locations of primers used to reconstitute the lac operon cassette in pGGAselect. For clarity, the prefix `OL` has been omitted. Upper panel: non-optimal set of overhangs. Lower panel: optimal set of overhangs. See Supplementary Figure S15 for a gel showing PCR products.

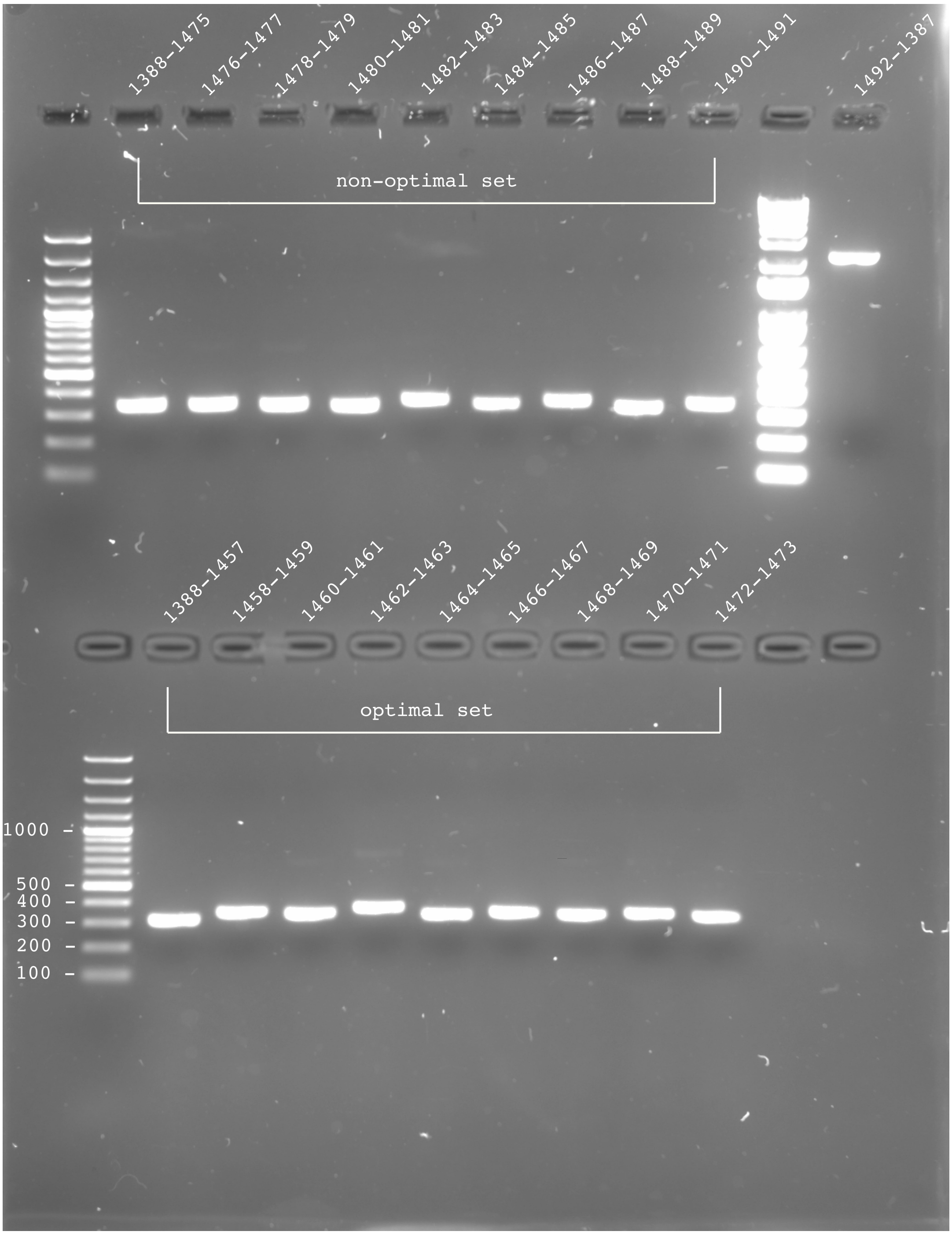


Supplementary Figure S15: 2% agarose gel displaying PCR products post-purification using the Monarch PCR & DNA Cleanup Kit (5 $\mu$g). The fragment generated by primers OL1492 and OL1387 is shared between both sets.

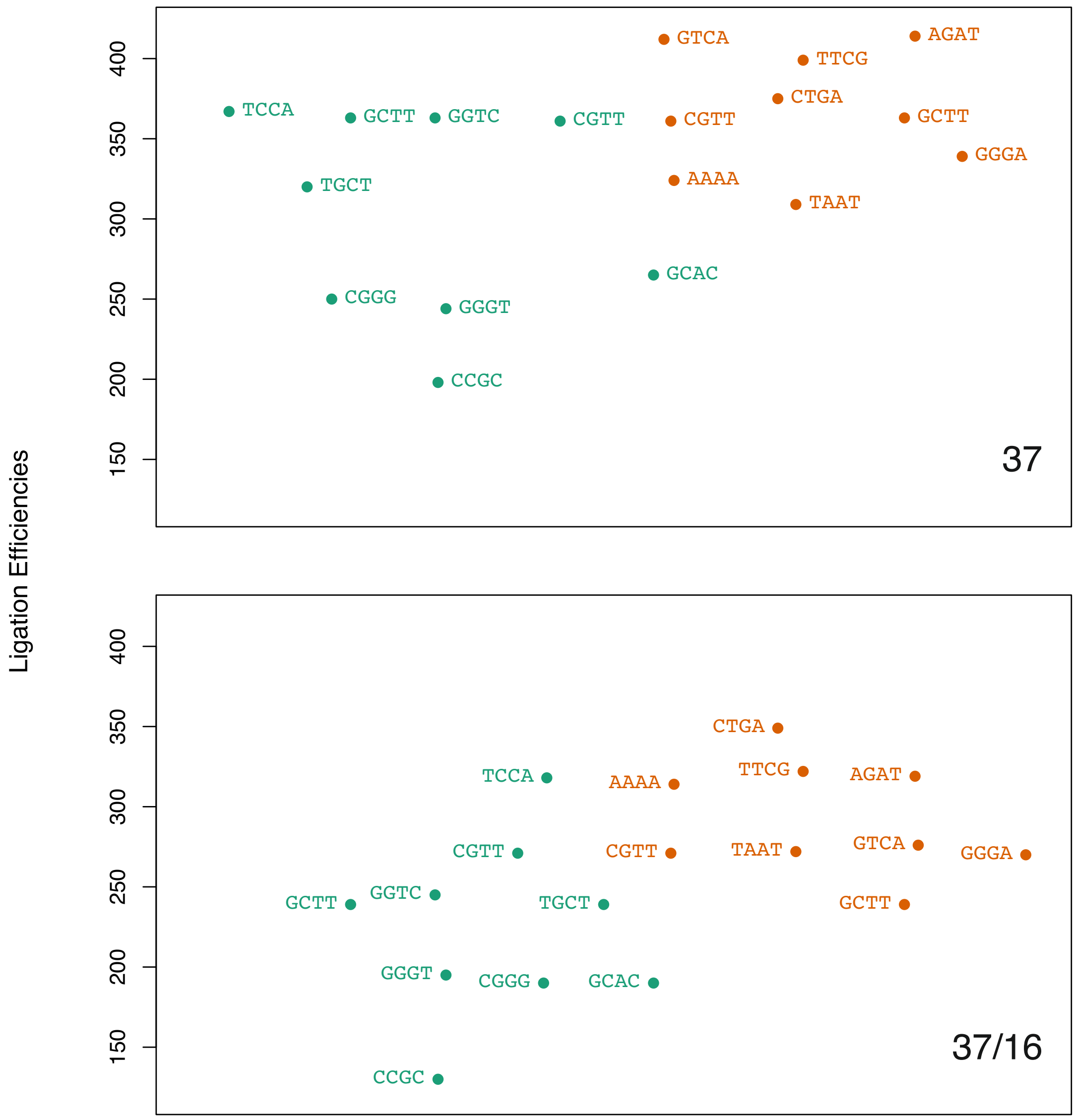


Supplementary Figure S16: Ligation efficiencies obtained from the NEBridge Ligase Fidelity Viewer. Values are reported for the optimal set (green) and the non-optimal set (orange) for both a static assay and a cycling assay (BsaI-HFv2).
